# Continuous cell type diversification throughout the embryonic and postnatal mouse visual cortex development

**DOI:** 10.1101/2024.10.02.616246

**Authors:** Yuan Gao, Cindy T. J. van Velthoven, Changkyu Lee, Emma D. Thomas, Darren Bertagnolli, Daniel Carey, Tamara Casper, Anish Bhaswanth Chakka, Rushil Chakrabarty, Michael Clark, Marie J. Desierto, Rebecca Ferrer, Jessica Gloe, Jeff Goldy, Nathan Guilford, Junitta Guzman, Carliana R. Halterman, Daniel Hirschstein, Windy Ho, Katelyn James, Rachel McCue, Emma Meyerdierks, Beagan Nguy, Nick Pena, Trangthanh Pham, Nadiya V. Shapovalova, Josef Sulc, Amy Torkelson, Alex Tran, Herman Tung, Justin Wang, Kara Ronellenfitch, Boaz Levi, Michael J. Hawrylycz, Chelsea Pagan, Nick Dee, Kimberly A. Smith, Bosiljka Tasic, Zizhen Yao, Hongkui Zeng

## Abstract

The mammalian cortex is composed of a highly diverse set of cell types and develops through a series of temporally regulated events that build out the cell type and circuit foundation for cortical function. The mechanisms underlying the development of different cell types remain elusive. Single-cell transcriptomics provides the capacity to systematically study cell types across the entire temporal range of cortical development. Here, we present a comprehensive and high-resolution transcriptomic and epigenomic cell type atlas of the developing mouse visual cortex. The atlas was built from a single-cell RNA-sequencing dataset of 568,674 high-quality single-cell transcriptomes and a single-nucleus Multiome dataset of 194,545 high-quality nuclei providing both transcriptomic and chromatin accessibility profiles, densely sampled throughout the embryonic and postnatal developmental stages from E11.5 to P56. We computationally reconstructed a transcriptomic developmental trajectory map of all excitatory, inhibitory, and non-neuronal cell types in the visual cortex, identifying branching points marking the emergence of new cell types at specific developmental ages and defining molecular signatures of cellular diversification. In addition to neurogenesis, gliogenesis and early postmitotic maturation in the embryonic stage which gives rise to all the cell classes and nearly all subclasses, we find that increasingly refined cell types emerge throughout the postnatal differentiation process, including the late emergence of many cell types during the eye-opening stage (P11-P14) and the onset of critical period (P21), suggesting continuous cell type diversification at different stages of cortical development. Throughout development, we find cooperative dynamic changes in gene expression and chromatin accessibility in specific cell types, identifying both chromatin peaks potentially regulating the expression of specific genes and transcription factors potentially regulating specific peaks. Furthermore, a single gene can be regulated by multiple peaks associated with different cell types and/or different developmental stages. Collectively, our study provides the most detailed dynamic molecular map directly associated with individual cell types and specific developmental events that reveals the molecular logic underlying the continuous refinement of cell type identities in the developing visual cortex.

## Introduction

The cerebral cortex of the mammalian brain is considered the newest invention of evolution, endowed with the ability to control a wide range of flexible and motivated behaviors, and dramatically expanded in species with more sophisticated cognitive functions (including human). As such, the cortex has been a prime subject for the investigation of the diverse cell types it contains and how these cell types form functionally specific neural circuits^1–3^. The cortex is a six-layered sheet structure, with specific glutamatergic excitatory and GABAergic inhibitory neuron types occupying different layers. These excitatory and inhibitory neurons are locally connected with each other across layers, forming vertically integrated canonical circuits called “cortical columns”^4^. Cortical columns are similarly repeated across the entire cortical sheet and are grouped into multiple different cortical areas each defined by its unique input/output connectivity and function. In addition to within-area local connections, excitatory neurons also extend their axons to other specific cortical areas (as well as subcortical areas) and form extensive interareal networks^4,5^. The expansion of cortex during species evolution mainly involves the emergence of new cortical areas and thus more complex interareal networks^6,7^.

Cell types in the cortex can be defined by multiple cellular properties, including gene expression, morphology, physiology, connectivity, or the various combinations of these properties^3,8–10^. Over the past decade, single-cell transcriptomics has provided the most comprehensive and detailed cell type classification, defining ∼100 transcriptomic cell types (T-types) in each cortical area of the *adult* brain that is largely consistent across areas and across species (e.g., from mouse to human)^11–14^. These T-types can be hierarchically organized into classes and subclasses, reflecting their varied relatedness that is likely rooted in the evolutionary and developmental histories of the cell types^10^. Specifically, in each cortical area, ∼28 cell subclasses are defined. These include 9 glutamatergic subclasses organized by layers and long-range projections: L2/3 IT (intratelencephalic projecting), L4/5 IT, L5 IT, L6 IT, L6 Car3, L5 ET (extratelencephalic projecting), L5/6 NP (near-projecting), L6 CT (corticothalamic projecting), and L6b; 8 GABAergic subclasses organized by developmental origins: Lamp5, Sncg, Vip, Pvalb, Pvalb chandelier, Sst, Sst Chodl, and Lamp5 Lhx6; 3 glial subclasses: astrocytes, oligodendrocytes, and oligodendrocyte precursor cells (OPC); 3 immune subclasses: microglia, border associated macrophages (BAM), and lymphoid cells; and 5 vascular subclasses: vascular leptomeningeal cells (VLMC), arachnoid barrier cells (ABC), endothelial cells, pericytes, and smooth muscle cells (SMC)^14,15^.

Multi-modal integrative approaches have been used to align the different levels of transcriptomic cell types to morphology, physiology and connectivity, and in some cases to refine cell type definition^14,16,17^. For example, using Patch-seq, the GABAergic neurons in the mouse visual cortex are classified into 28 morpho-electric-transcriptomic types (MET-types), representing a coarser resolution from the original 61 T-types but with higher cross-modality concordance within each MET-type^16^. Importantly from the perspective of circuitry, most of these GABAergic MET-types exhibit layer specificity. Computational matching of local dendritic and axonal morphology has allowed assigning T-type identities to reconstructed neurons in mouse visual cortex that have long-range projection patterns or synaptic connectivity profiles derived from light or electron microscopy data^18,19^. Cell-type targeting genetic tools, barcoded viruses and spatial transcriptomic approaches have also been used to relate transcriptomic identities to connectional or functional properties^20–22^.

A fundamental question in neuroscience is how the extraordinary cell type diversity and neural circuit specificity emerges during brain development. Uncovering the precise developmental processes and concomitant changes at molecular, cellular, connectional and functional levels, and identifying key factors driving these changes, will enable a better understanding of the mechanisms underlying brain development and further, how the process goes awry in neurodevelopmental disorders.

The development of the mammalian cortex has been extensively investigated over the years^23–27^. It is now well known that glutamatergic neurons as well as astrocytes and oligodendrocytes are generated within the dorsal pallium (which becomes the cortex later), whereas GABAergic neurons are generated in the subpallium and undergo long-distance migration into the cortex following specific routes^28,29^, and immune and vascular cell types originate outside the brain *per se*. In both pallium and subpallium, progenitors in the ventricular and subventricular zones (VZ and SVZ) progressively give rise to radial glia (RG) cells, intermediate progenitors (IP) and immature neurons (IMN). In the developing cortex, glutamatergic neurons in different layers are thought to be generated sequentially and migrate radially to reach their target layers in an inside-out manner^30,31^. After neurogenesis is complete, RGs switch to gliogenesis and generate astrocytes and OPCs/oligodendrocytes (though some oligodendrocytes also come from subpallium regions). Postmitotically, all cell types go through specific maturation processes. Glutamatergic and GABAergic neurons go through an extensive series of dendritic and axonal arborization, synapse formation, and activity-dependent circuit refinement. In particular, visual cortex goes through a series of experience-independent and experience-dependent circuit development to acquire increasingly refined visual response properties^32^.

There remain substantial gaps in our understanding of the developmental processes and mechanisms. It is still unclear when specific cell type identities are established, to what extent are cell types observed in the adult cortex established during the embryonic stage, and how lineage bifurcation decisions occur. In the postnatal developmental period, many processes are at play with overlapping time courses, such as intrinsic neuronal activities, influence of external sensory inputs, incoming and outgoing long-range connections, formation of local excitatory and inhibitory circuit motifs, and neuronal and non-neuronal cell-cell interactions. Consequently, cells are undergoing rapid state transitions. Despite the discovery of many genes, proteins and epigenetic signatures involved in these processes, we have very little systematic knowledge about what cell-type specific dynamics exist, how cell-type specific circuits are formed, and what mechanisms drive cell type differentiation and maturation. To address these questions, it is critical to investigate developmental changes at the single cell level and link these changes across time with cell type specificity.

Here, we report a comprehensive and high-resolution single-cell transcriptomic and epigenomic atlas of the developing mouse visual cortex, with unprecedented dense temporal profiling throughout the embryonic and postnatal developmental stages. Dense temporal profiling allowed us to build a transcriptomic developmental trajectory map of all excitatory, inhibitory, and non-neuronal cell types in the visual cortex, identifying branching points demarking the emergence of new cell types at specific developmental ages and defining molecular signatures of cellular diversification. In addition to neurogenesis, gliogenesis and early postmitotic maturation in the embryonic stage which gives rise to all the cell classes and nearly all subclasses, we observe that more refined cell types emerge throughout the postnatal differentiation process, including the late emergence of many cell types during the eye-opening stage (P11 to P14 days) and around the time of weaning (P21) which is also the start of critical period of experience-dependent plasticity, suggesting continuous cell type diversification at different stages of cortical development. We also derived a chromatin accessibility map across the cell-type development trajectories and identified key transcription factor regulators for cell-type specific epigenetic changes. The high-resolution transcriptomic and epigenomic cell type atlas and trajectory map also allowed us to identify many gene co-expression modules and chromatin accessibility peak modules for specific cell types and developmental ages, which collectively provides a dynamic molecular map directly associated with individual cell types and specific developmental events that will facilitate many future mechanistic studies of the different aspects of cortical development.

## Results

### Creation of a mouse visual cortex developmental cell-type atlas

We generated two types of large-scale, single-cell-resolution datasets for the developing mouse visual cortex, using single-cell RNA-sequencing (scRNA-seq) and single-nucleus Multiome (combination of snRNA-seq and snATAC-seq). We used the scRNA-seq data to generate a transcriptomic cell-type atlas and developmental trajectory map. We then used the Multiome data to reconstruct the epigenetic chromatin accessibility landscape across developmental cell type atlas and trajectory (described later).

We first generated 92 scRNA-seq libraries using 10x Genomics Chromium v3 (10xv3), resulting in a dataset of 919,547 single-cell transcriptomes (**Supplementary Table 1**). The scRNA-seq data densely covers the embryonic and postnatal periods with a total of 35 time points: embryonic day E11.5, E12.5, E13.5, E14.5, E15.5, E16.5, E17.0, E17.5, E18.0, E18.5, postnatal day P0, P1, P2, P3, P4, P5, P6, P7, P8, P9, P10, P11, P12, P13, P14, P15, P16, P17, P19, P20, P21, P23, P25, P28, plus adult stage P54-68 (collectively simplified as P56) (**Fig. 1a**). We established a series of stringent quality control (QC) metrics (e.g., gene detection, QC score, and doublet score, see **Methods, Supplementary Table 2**), which were also adopted by our previous studies^15^ to identify low-quality single-cell transcriptomes. After the QC-filtering, we obtained 761,419 high-quality cells (**Extended Data Fig. 1**).

**Figure 1.**
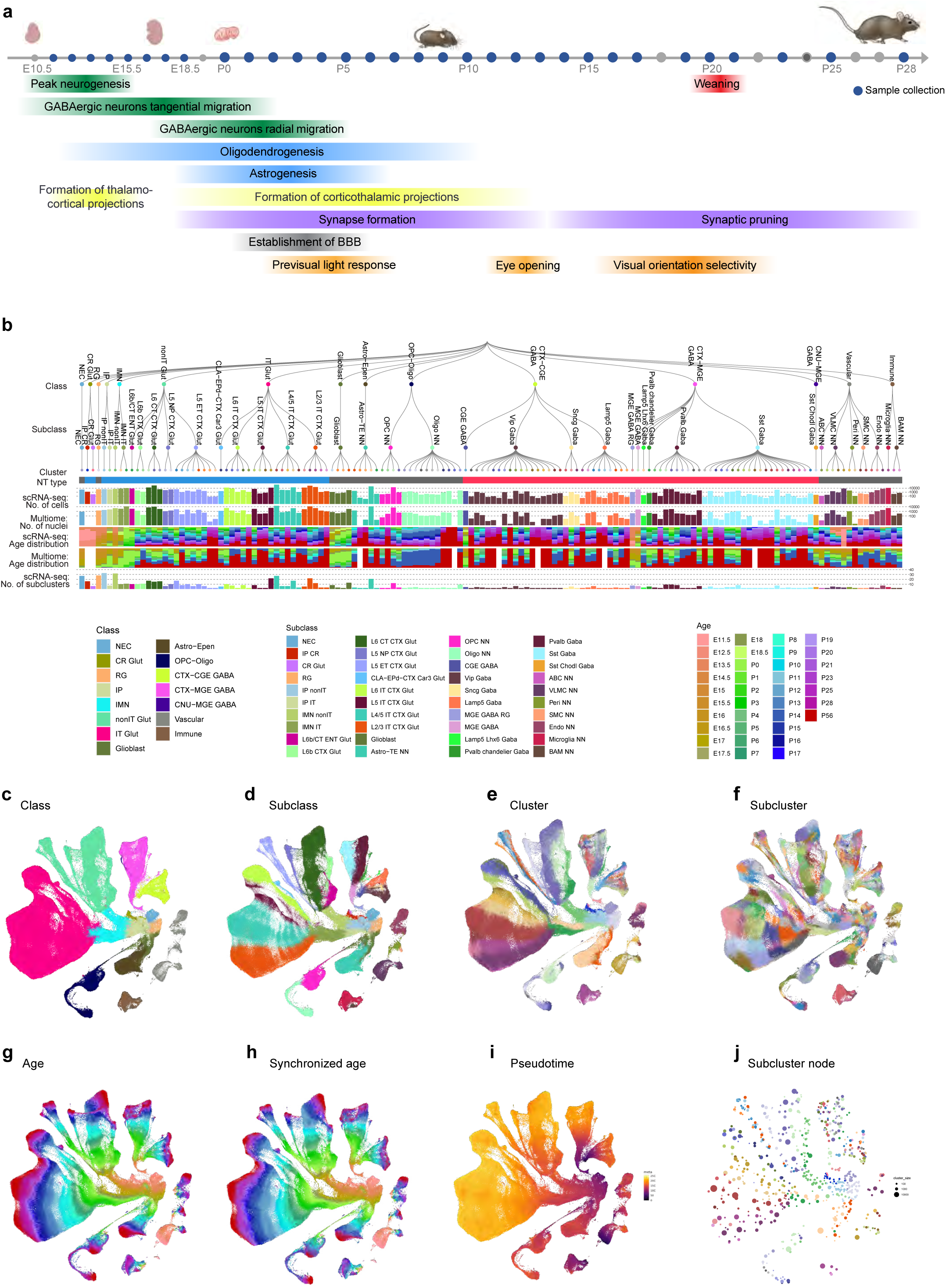
Transcriptomic developmental cell type atlas of the mouse visual cortex. **(a)** Schematic timeline of samples collected in this study along with major developmental events of the isocortex. **(b)** The transcriptomic taxonomy tree of 148 clusters organized in a dendrogram (10xv3 n = 568,674 cells; 10x multiome n = 194,545 nuclei). The classes and subclasses are marked on the taxonomy tree. Full cluster names are provided in **Supplementary Table 3**. Bar plots represent (top to bottom): major neurotransmitter type, number of scRNA-seq cells, number of multiome cells, age distribution of scRNA-seq cells, age distribution of multiome nuclei, and number of scRNA-seq subclusters for each cluster. **(c-i)** UMAP representation of all cell types colored by class (c), subclass (d), cluster (e), subcluster (f), age (g), synchronized age (h), and pseudotime (i). **(j)** Constellation plot showing the UMAP centroids of subcluster nodes colored by cluster. NEC, neuroepithelial cells. CR, Cajal–Retzius cells. RG, radial glia. IP, intermediate progenitors. IMN, immature neurons. IT, intratelencephalic. ET, extratelencephalic. L6b, layer 6b. NP, near-projecting. CGE, caudal ganglionic eminence. MGE, medial ganglionic eminence. Astro, astrocytes. Oligo, oligodendrocytes. OPC, oligodendrocyte precursor cells. GABA, GABAergic. Glut, glutamatergic. NN, non-neuronal.

In these data, the precise developmental age of each sampled cell is unknown—only the collection time from which it was sampled. Additionally, due to the challenges to determine the age of the prenatal samples, the collection time might not be accurate, especially for early developmental stages. Importantly, cells from the same collection time point may exist at different developmental stages and maturation levels. To synchronize cells in the same developmental stage, we performed global clustering of all cells and predicted developmental ages based on transcriptomes (**Methods**), such that cells with the same predicted ‘synchronized age’ have more homogeneous temporal transcriptomic profiles (**Extended Data Fig. 2a**). Developmental ages that are difficult to discriminate in transcriptomic space were further merged into synchronized age bins for downstream analysis (**Extended Data Fig. 2d**).

To build the developmental trajectory of the adult cell types, we first conducted label transfer using the adult mouse whole brain taxonomy we recently established^15^. This Allen Brain Cell – Whole Mouse Brain (ABC-WMB) Atlas served as a reference for cells at the adult stage to assign cell type identities at cluster level. Adult cell type identifies were then propagated to younger cells by performing sequential cell type label transfer from older to younger synchronized ages for all postnatal ages (**Methods, Extended Data Fig. 2a, 3, Supplementary Table 3, Fig. 1b**). Overall, for the P20_28 age bin, we label-transferred 34 glutamatergic clusters, 51 GABAergic clusters, and 12 glial clusters derived from the adult ABC-WMB Atlas, capturing majority of the cell type diversity in the adult mouse visual cortex.

For the embryonic time points, we used the La Manno et al^33^ developing mouse brain scRNA-seq dataset as reference to identify broad cell types (**Methods, Supplementary Table 3, Fig. 1b**). We focused on our global clusters which had the highest proportion of cells from the prenatal stages. Global clusters which were mapped to Radial glia were assigned as neuroepithelial cells (NEC, expressing *Hmga2*) or RG (expressing *Sox2*, *Pax6* and *Hes5*), same name at class, subclass and cluster levels in each case. Global clusters mapped to Neuroblast were assigned the IP class (expressing *Eomes*). Those IP clusters that were enriched with *Lhx9*, *Rmst*, *Nhlh1* and *Nhlh2* were annotated as the IP nonIT subclass or cluster (same name at these two levels), and those enriched with *Pou3f2* were the IP IT subclass/cluster. Global clusters which were highly enriched with neuronal markers *Ncam1*, *Dcx* and *Neurod6* and had low expression of *Eomes* were annotated as the IMN class. Among IMNs, those enriched with *Fezf2* were the IMN nonIT subclass/cluster and those enriched with *Cux1* were the IMN IT subclass. In addition, global cluster enriched with markers of the preplate Cajal-Retzius (CR) cells (*Ebf1*, *Ebf2*, *Ebf3*, *Reln*, *Calb2*, *Tbr1* and *Trp73*) were assigned as the CR Glut class, which were further divided into the IP CR (expressing *Eomes*) and the mature CR Glut (expressing *Reln* and *Calb2*) subclasses/clusters.

Within each cluster at each synchronized age bin, we performed iterative *de novo* clustering on the QC-qualified cells, resulting in an initial transcriptomic cell-type taxonomy with 1,845 subclusters across all developmental stages. We then merged subclusters within each cluster based on the differentially expressed (DE) genes between all pairs of subclusters as in our previous study^15^. To finalize the transcriptomic cell-type taxonomy and atlas, we conducted detailed annotation of all the subclasses, clusters and subclusters based on their molecular signatures. During this process (including integration with the Multiome dataset, see below, **Extended Data Fig. 1a**), we identified and removed an additional set of ‘noise’ subclusters (usually doublets) that had escaped the initial QC process, or subclusters outside cortex, resulting in a final set of 568,674 high-quality single-cell transcriptomes that form 714 subclusters.

To organize the complex molecular relationships, we present a high-resolution transcriptomic cell-type taxonomy and atlas for the adult and developing mouse visual cortex with four nested levels of classification: 15 classes, 40 subclasses, 148 clusters and 714 subclusters (**Supplementary Table 3**), which includes all known neuronal and non-neuronal cell classes of the developing neocortex from literature^27^, as well as many transitional cell types and subtypes discovered here. We provide several representations of this atlas for further analysis: a dendrogram at cluster resolution along with bar graphs displaying various metadata information (**Fig. 1b**), and uniform manifold approximations and projections (UMAPs) at single-cell resolution colored with different types of metadata information (**Fig. 1c-i**). We also generated a list of 6,724 DE genes that differentiate among all clusters and subclusters (**Supplementary Table 4**). The scRNA-seq dataset shows clearly that cells collected at different ages are well-separated in the transcriptomic space (**Fig. 1g,h**), indicating distinct transcriptomic changes occurring between different developmental stages. The transcriptomic temporal trajectory aligns closely with the ages, with cells from adjacent ages also being adjacent in the transcriptomic space.

Of the 148 clusters (**Supplementary Table 3**), 132 clusters (containing 517 subclusters) are aligned with the adult ABC-WMB Atlas^15^, representing maturing cell types. These clusters belong to 27 of the above mentioned 28 canonical cortical cell subclasses^14,15^ (without lymphoid cells), plus one that might be destined to the entorhinal cortex of the hippocampal formation (see below), under a total of 9 classes. The labels of these 28 subclasses and 132 clusters are adopted from the ABC-WMB Atlas, while some of their class labels are modified to be more consistent with the embryonic classes. The remaining 16 clusters (containing 197 subclusters) represent progenitor cells and immature neurons in embryonic and perinatal stages and belong to 12 subclasses under 8 classes.

Neuronal cell types constitute a large proportion of the developmental atlas, including 10 classes: NEC, CR Glut, RG, IP, IMN, nonIT Glut, IT Glut, CTX-CGE GABA, CTX-MGE GABA, and CNU-MGE GABA (**Fig. 1b,c**). The 10 classes are further divided into 29 subclasses, 109 clusters and 599 subclusters. The nonIT Glut class consists of four main glutamatergic subclasses – L5 ET, L5 NP, L6 CT, and L6b, plus a L6b/CT ENT subclass that is mostly present at E17-P2 and may be destined to the adult entorhinal cortex based on our mapping result (**Fig. 1b,d**). The IT Glut class contains four main subclasses – L2/3 IT, L4/5 IT, L5 IT, and L6 IT, plus a CLA-EPd-CTX Car3 subclass that consists of a distinct L6 cell type shared with the claustrum and the endopiriform nucleus^13,20^ (**Fig. 1b,d**).

It has been known that cortical GABAergic neurons originate from the embryonic ventral telencephalon, or subpallium, and migrate over long distances to populate the cortex. All of the cortical GABAergic neurons are born in three subpallial progenitor zones: the caudal ganglionic eminence (CGE), medial ganglionic eminence (MGE), and the preoptic area (POA)^25^. Neural progenitor cells in these regions transition from RG, that divide on the surface of the ventricle, to IPs with limited self-renewal capacity, before exiting the cell cycle to generate IMN^34^. The IMNs migrate from the subpallium, through distinct migratory streams, to the cortex where the IMNs mature^35,36^. Our data show that MGE-derived GABAergic progenitors differentiate into four subclasses, Sst Gaba, Pvalb Gaba, Pvalb chandelier Gaba and Lamp5 Lhx6 Gaba, in the CTX-MGE class, as well as one subclass, Sst Chodl Gaba, in the CNU-MGE class^15,37^ (**Fig. 1b,d**). Our previous study showed that the Sst Chodl Gaba subclass mainly contains striatal *Sst*+ interneurons, with 2 clusters specifically located in cortex^37^. The CGE GABA progenitors gradually differentiate into Lamp5 Gaba, Vip Gaba and Sncg Gaba subclasses.

All non-neuronal cell types are classified into 5 classes: Glioblast, OPC-Oligo, Astro-Epen, Immune and Vascular, which are further divided into 11 subclasses (**Fig. 1b-d**). Glioblast (expressing *Qk*) is the progenitor for the OPC-Oligo and Astro-Epen classes. The OPC–Oligo class contains two subclasses, OPC (expressing *Olig1*, *Olig2*, *Pdgfra*) and oligodendrocytes (Oligo; expressing *St18*, *Opalin*). The Astro-Epen class contains one subclass of telencephalic astrocytes, Astro-TE (expressing *Apoe*, *Aldh1l1* and *Slc1a3*). The Immune class consists of two subclasses: microglia (expressing *Siglech*, *Sall1* and *Ifitm10*) which are first observed at E11.5 and BAM (expressing *F13a1*, *Pf4*, *Mrc1*) which emerges at P5. The Vascular class consists of 5 subclasses: ABC (expressing *Slc47a1*), VLMC (expressing *Apod* and *Slc6a13*), pericytes (expressing *Kcnj8*), SMC (expressing *Acta2* and *Myh11*), and endothelial cells (Endo; expressing *Ly6c1* and *Slco1a4*). ABC cells are very rare across all time points. SMCs emerge at P9, while the other Vascular cell types are present since E11.5.

### Building cell type development trajectories

Trajectory analysis is an essential tool for studying the dynamic process of cellular development and differentiation. Popular computational methods such as Monocle^38^, PAGA^39^, Slingshot^40^ and RNA velocity^41^ leverage the gradients in transcriptomic space to infer cell type trajectory. However, one of the main challenges with these tools is to deconvolute the temporal gradient with other gradients associated with cell type heterogeneity. We found that these tools were not successful to derive the trajectory of the cortical development with desired cell type resolution. For example, the trajectories inferred by Monocle3 ^42^ switched back and forth between different layers and different ages for IT cells (**Extended Data Fig. 2c**), making the results extremely difficult to interpret. Although it is inherently difficult to disentangle the biological implications of different transcriptomic gradients, the age information for each sample can substantially simplify this problem, especially for postnatal development.

Given the cell type identities at the adult stage, we were able to progressively propagate cell type identities between two adjacent ages (see above), as all the cells in the later age evolve from cells in the earlier age. Since our atlas had dense temporal sampling, we found that identifying corresponding cell types in two adjacent time points that have only subtle transcriptomic differences could be readily solved using existing methods such as Seurat label transfer (**Methods**, **Extended Data Fig. 2a,b, 3**). Here, edge weights of the trajectory tree were defined based on mutual nearest neighbors (MNN) in the integrated space **(Methods, Extended Data Fig. 2b)**. For this task, instead of using actual age, we used the synchronized age as defined above, so cells that are developmentally more advanced or delayed within the same age are reassigned based on transcriptomic signatures. This strategy worked well until there was too much ambiguity in cell type assignment at earlier developmental times, and thus we switched the cell type nomenclature from adult types to developmental types such as RG, IP and IMN. During embryonic development, cells evolve more rapidly, and cells within the same age can be present in different development states. During this period, the transcriptome gradient corresponding to differentiation is dominant, while cell type diversity is much simpler compared to the later developmental stage. We found that established methods such as Monocle3 worked well in this case. Therefore, we defined the embryonic trajectory via the same MNN approach but using Monocle3 based pseudo-time instead of synchronized age.

Overall, we retained all edges between a cluster and its potential antecedents that have edge weights > 0.2 (**Supplementary Table 5**). To simplify visualization and conceptualization of the developmental process, we chose the edge with the max weight between a cluster and one antecedent to build the developmental trajectory map across the entire timeline from E11.5 to P56 (**Fig. 2a**, **3**). Of note, the total of 963 chosen edges with max weights to build all the trajectories had an average weight of 0.72 (and over 87% of them had weights > 0.5), whereas the 331 unchosen edges all had weights < 0.5 with an average of 0.29 (**Supplementary Table 5**). This result indicates a relatively unambiguous trajectory pattern. We then computed the global pseudo-time based on the entire developmental trajectory map (**Methods, Fig. 1i, 3**).

**Figure 2.**
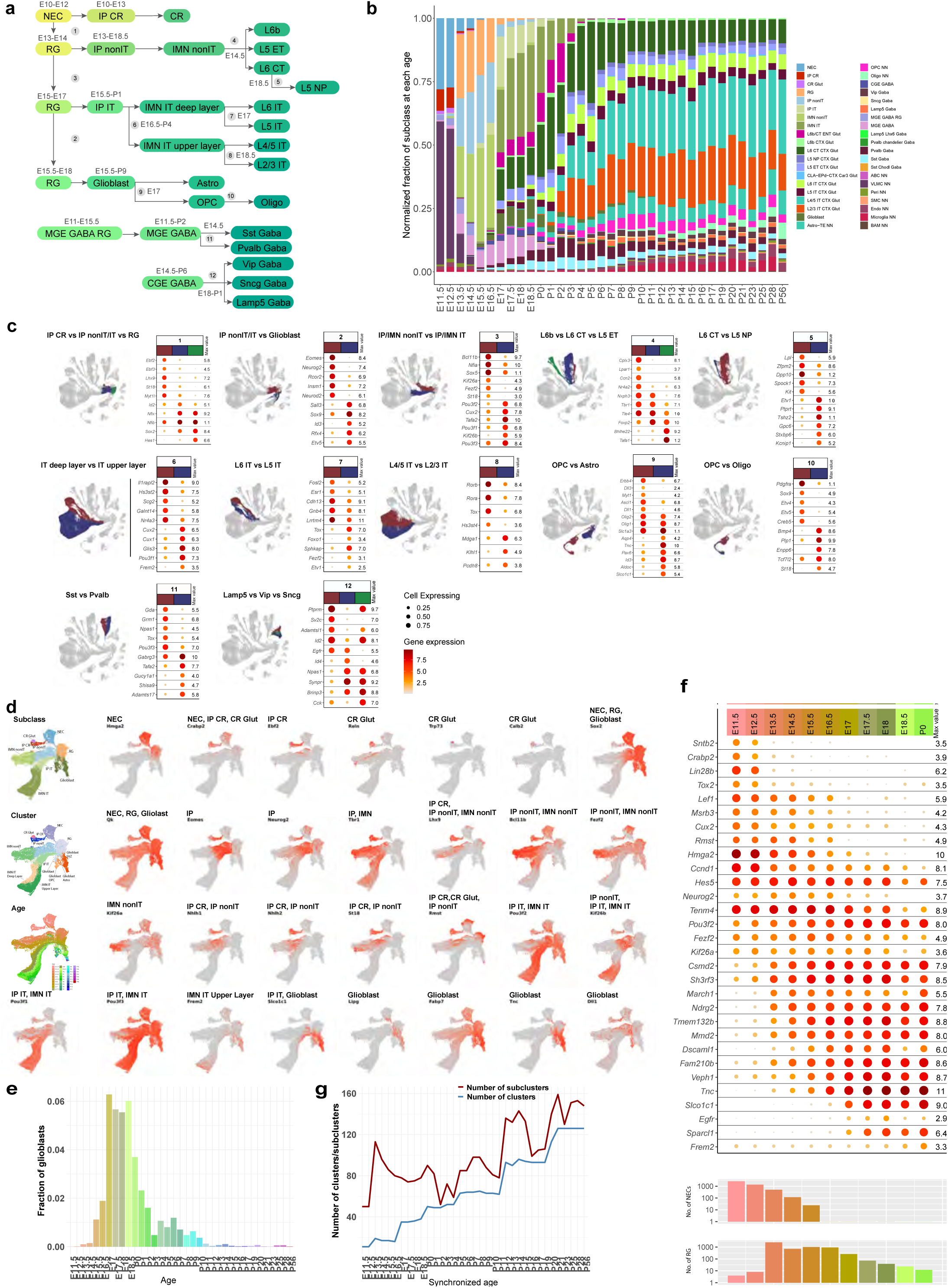
Developmental trajectories of visual cortex cell subclasses. **(a)** Transcriptomic trajectories of VIS cortical subclasses with estimated timing of onset and major branching nodes. **(b)** Relative proportions of cells corresponding to the different cell subclasses at each age. E11.5 and E12.5 are mainly composed of NEC, IP CR, CR, MGE GABA RG, VLMC, and microglia. RG, IP nonIT and IMN nonIT constitute a large proportion from E13.5 to E16.5. IP IT and IMN IT have large proportions from E17.0 to E18.5. Neuronal subclass composition starts to be stable from P6. Note that relative proportions between neuronal and non-neuronal cells do not reflect the actual situation due to the variable FACS plans employed for different scRNA-seq libraries (**Methods, Extended Data Fig 1d, Supplementary Table 1**). **(c)** UMAP representations of major branching nodes shown in (a) and dot plots showing marker gene expression in each descendant branch of each branching node. Dot size and color indicate proportion of expressing cells and average expression level of a marker gene in each subclass, respectively. **(d)** UMAP representations of early developmental cell types colored by subclass, cluster, age, and expression of key marker genes separating different trajectories. **(e)** Fraction of glioblast cells at each age. **(f)** Dot plot showing expression of DE genes across embryonic ages and P0 in NEC and RG populations. Numbers of NEC and RG cells at each age point are shown at the bottom. **(g)** Number of clusters and subclusters at each synchronized age.

**Figure 3.**
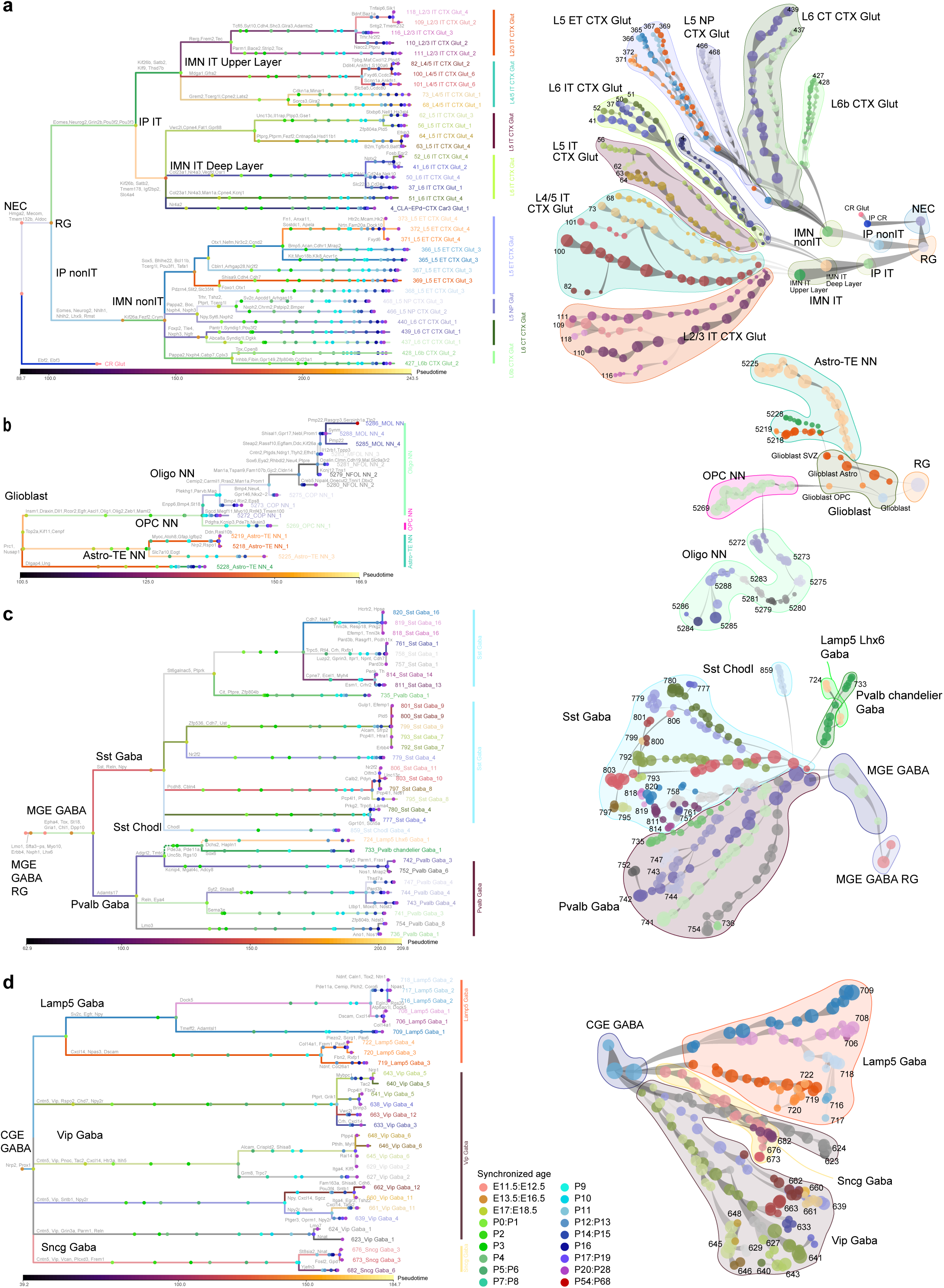
Developmental trajectories of visual cortex cell types. (a-d) Transcriptomic trajectory tree (left) and constellation plot (right) of glutamatergic (a), neuroglia (b), MGE (c), and CGE (d) clusters, which are grouped into subclasses. Each branch represents a cluster, whose name is labeled in the same color. In (a), 36 glutamatergic clusters derived from neuroectoderm. Root is NEC and tips are E14.5 terminal CR Glut cluster and P56 terminal nonIT and IT cell clusters. In (b), for neuroglia, root is RG and tips are 15 P56 terminal OPC-Oligo and Astro-TE clusters. In (c-d), MGE and CGE GABAergic neurons are derived from distinct trajectory trees. For MGE, root is MGE GABA RG and tips are 32 P56 terminal CTX-MGE and CNU-MGE clusters. For CGE, tips are 29 P56 terminal CTX-CGE clusters. Marker genes for each branch point are shown along each branch. Branch lengths represent pseudo-time, a measurement of how much progress an individual cell type has made through a process such as cell differentiation. Internal nodes on each branch represent cells from that cluster subdivided by synchronized age bins.

We constructed a branched trajectory tree for the neuronal and glial subclasses in visual cortex (**Fig. 2a**), supported by key marker genes for each branching node (**Fig. 2c,d, Extended Data Fig. 4**). The relative proportions of most subclasses change dramatically during the embryonic period and start to stabilize after P2, but exact days differ between subclasses (**Fig. 2b, Supplementary Table 3**). The trajectory tree reveals that the earliest cell type emerges from NEC is IP CR (before E11.5) which matures into CR cells, then RG emerge at E13 followed immediately by the emergence of IP nonIT cells, and both IP IT cells and glioblasts appear around E15.5 (**Fig. 2a,b**). IP nonIT cells transition into IMN nonIT cells, from which three subclasses of nonIT neurons, L5 ET, L6 CT and L6b, emerge at E14.5, while the fourth subclass, L5 NP, appears to derive from L6 CT at E18.5. IP IT cells transition into IMN IT cells, and subsequently, deep-layer IMN IT turns into L6 IT and L5 IT neurons at E17 and upper-layer IMN IT turns into L4/5 IT and L2/3 IT neurons at E18.5. In the meanwhile, glioblasts give rise to astrocytes and OPCs around E17. Separately for GABAergic neuron classes, MGE RG and MGE cells appear in cortex before E11.5, and MGE cells turn into Sst and Pvalb neurons at E14.5; CGE cells appear in cortex later around E14.5, and they turn into Vip, Sncg and Lamp5 neurons at E18-P1.

Based on the classical view of neurogenesis, neocortical progenitors begin to produce excitatory neurons as early as E10.5 in mice at which time NECs extend radial fibers and gradually transition into RG^43,44^. The earliest-born neurons migrate away from the ventricular surface, segregating from progenitors to form the preplate^45^. The preplate is then split, by later-born neurons migrating into the preplate, into the marginal zone and subplate and establishing the cortical plate between the two^46,47^. From here deeper layers (L6 and L5) are established early, between E11.5 and E13.5, while superficial layers (L4, L2/3) are established later, between E14.5 and E16.5, in a so-called “inside-out” manner^43,45^.

In our data, at the earliest stage (E11.5 to E12.5), cells originating from pallium are mainly composed of NEC (*Hmga2* and *Ccnd1*), IP CR and the early born CR cells (*Ebf1*, *Ebf2*, *Ebf3*, *Reln*, *Calb2*, *Tbr1* and *Trp73*) (**Fig. 2a-d, branching node 1, Extended Data Fig. 4**). CR cells migrate to the marginal zone, and disappear almost completely in the postnatal neocortex by programmed cell death, with a subpopulation surviving up to adulthood in hippocampus^48–50^. The IPs at this stage are antecedents of CR cells, hence are named the IP CR subclass; their expression of *Eomes* and *Neurog2* decreases as CR cells mature (**Fig. 2d**).

Beginning at E13, RG (*Sox2*, *Pax6* and *Hes1*) and IPs (*Eomes*, *Neurog2* and *Rcor2*) emerge and gradually transition into IMNs (*Dcx*, *Neurod1*, *Neurod2* and *Neurod6*). Our datasets suggest that most IPs generated between E13.5 and E16.5 are IP nonIT and transition into IMN nonIT neurons, whereas most IPs between E17 and P0 are IP IT and transition into IMN IT neurons (**Fig. 2b**). The nonIT neurons are located at deep layers, while most IT neurons are located at upper layers. This observation partially supports the traditional view of inside-out neurogenesis in which deep-layer neurons are generated before upper-layer neurons. However, the distinction should be more precisely categorized as nonIT vs IT. Deep-layer IT neurons are generated by IP cells born at later time than nonIT neurons, even though they co-localize in deep layers.

We observe clear molecular signatures that distinguish nonIT and IT lineages at IP and IMN stages. The nonIT IPs and IMNs express *Rmst*, *Lhx9*, *St18*, *Nhlh1*, *Nhlh2* and *Fezf2,* and the IT IPs and IMNs express high levels of *Pou3f1*, *Pou3f2, Pou3f3* and *Prdm16* (**Fig. 2a,c,d, node 3, Extended Data Fig. 4**). Also interestingly, while *Kif26a* is expressed in RG and *Kif26b* in IP cells, their expressions are transiently turned off before turned on again with specific expression of *Kif26a* in nonIT IMNs and Kif26b in IT IMNs (**Fig. 2d**), and both genes are downregulated again in adulthood. These two closely related paralogs in the kinesin family have mirrored temporal progression in distinct lineages, and *Kif26b* is known to play an important role in regulating adhesion of the embryonic kidney mesenchyme^51^.

Canonical deep-layer neuron markers *Fezf2*, *Bcl11b*, *Foxp2* and *Tle4* are all specific to nonIT lineage, but exhibit varying temporal dynamics (**Fig. 2a,c,d, Extended Data Fig. 4**). *Bcl11b* and *Fezf2* show enrichment in nonIT lineage at the IP stage, while *Foxp2* and *Tle4* only show nonIT enrichment at the late IMN stage. Well-known upper-layer regulators^52^ *Satb2* and *Cux2* show modest enrichment in IT lineage at IP stage, and the enrichment grows stronger in IMN stage (**Extended Data Fig. 4**). *Cux2* expression is further restricted to upper layer IT neurons in early postnatal stage, while the expression difference of *Satb2* between IT and nonIT lineage gradually decreases in late postnatal stages.

The transcriptomic difference of IT versus nonIT lineages is not only present in IPs, but already in RG at different ages (**Fig. 2f**). For example, RG at E13.5 and E14.5 express higher level of *Rmst*, which is enriched in nonIT lineage at IP stage; in contrast, RG at later ages show higher expression of *Pou3f2*, which is enriched in IT lineage at IP stage (**Fig. 2d,f**). *Rmst* is a long noncoding RNA, previously reported to interact with *Sox2* to regulate neurogenesis^53^. Our results suggest that it might do so in a time and state dependent manner, and likely involved in nonIT fate specification.

Recent studies have suggested that the transcriptional profile of cortical RG changes as they generate nonIT neurons, IT neurons, and glial cells^43,54^. However, it is still unclear if cell fate is driven by pre-specified progenitor populations, progressive fate-restriction as the cells develop, or a combination of both^44,54,55^. In our data, the divergence of progenitors for glutamatergic neurons (nonIT Glut and IT Glut) and glia (OPC-Oligo and Astro) may start as early as E15.5 (**Fig. 2a-e, node 2**), and the RG subclass shows a continuum of cells among different states (**Fig. 2f**). First, earlier-stage RG cells are enriched for *Neurog2* and *Tenm4*^56,57^, which may represent a committed neurogenic state, while expression of *Tnc* is seen in later-stage RG cells, which may represent a committed gliogenic state (**Fig. 2f**). Second, glioblasts in the non-neuronal branch emerge at E15.5 (**Fig. 2b,e**). These glioblasts express higher levels of *Fabp7*, *Lipg*, *Slco1c1*, *Tnc*, *Qk* and *Slc1a3* than RG, indicating their transition toward the glial cell lineage (**Fig. 2d,f**).

Our data suggests that RG already show complex temporal gene expression changes, and they exit the RG states at different ages carrying these temporal signatures to become IPs or glioblasts that are committed to differentiate into distinct neuronal (nonIT or IT) or glial lineages. These results are consistent with and could explain the observed heterogeneity in previous lineage tracing and transcriptomic profiling studies^54,55,58,59^.

### Developmental trajectories of glutamatergic neuron types

Our analysis indicates that the postmitotic immature neurons, IMN nonIT and IMN IT, progressively diversify into more distinct cell subclasses and types (**Fig. 1c-f, 2a, 3a**). In the nonIT lineage, IMNs (*Fezf2*, *Bcl11b* and *Neurod2*) emerge at E13.5, with increasing expression of *Foxp2*, *Tle4* and *Crym* at late IMN stage. This lineage splits around E17 into L6 CT, L5 ET, and L6b (**Fig. 2a-d, node 4, Extended Data Fig. 4**). The gene expression profile of the late IMN nonIT cells closely resembles that of L6 CT, the most prevalent subclass in the nonIT lineage and appearing the earliest. In L5 ET subclass, *Foxp2* and *Tle4* are downregulated, and *Pou3f1* and *Bhlhe22* are upregulated.

L6b subclass is believed to be derived from subplate with shared markers *Cplx3*, *Lpar1*, *Nr4a2*, and *Ccn2* (**Fig. 2c, Extended Data Fig. 4**). *Nxph4* and *Pappa2* are specific L6b markers postnatally, but they are also expressed at IMN and earlier stages. Subplate cells largely die out by P3, and their remnants become L6b cells^60^. There is a distinct population of L6b like cells with shared expression of subplate markers *Cplx3*, *Lpar1*, *Nr4a2*, but not *Ccn2*, *Nxph4* and *Pappa2*. This population is more abundant than L6b at E17-P3 (**Fig. 2b**), with specific expression of *Cyp26b1* and *Cobll1*, and mapped to adult L6b/CT ENT subclass. Based on Allen Developing Brain Atlas^61^ (developingmouse.brain-map.org), *Cyp26b1* is expressed specifically in the entorhinal and piriform cortical regions at E18.5, which further suggests that these neurons are likely located outside the visual cortex. L5 NP subclass emerges later than the above three nonIT subclasses, around E18.5 (*Ptprt*, *Tshz2*; **Fig. 2a-c, node 5**, **Extended Data Fig. 4**). It appears to derive from early L6 CT cells, but how it emerges remains unclear, with very few transition cells connecting to the closest antecedent type. Unlike most other subclasses of cortical glutamatergic neurons, L5 NP cells do not have long-range projections, and their functions remain elusive^11,62^.

The IP IT subclass gives rise to the IMN IT subclass, which is further divided into deep-layer and upper-layer IMN populations (**Fig. 2a,c, node 6**). *Frem2* is enriched in the upper layer IMN population, with this enrichment persisting until P10 and then gradually fading after eye opening (**Fig. 2d**). More markers emerge that split deep-layer IT and upper-layer IT populations after IMN stage, including *Il1rapl2* and *Hs3st2* enriched in L5 IT and L6 IT subclasses and *Cux1* and *Cux2* in L2/3 IT and L4/5 IT subclasses (**Fig. 2c, node 6, Extended Data Fig. 4**). The IMN IT Deep Layer cluster continues to differentiate into L5 IT and L6 IT subclasses around E17 with enrichment of *Fosl2* in L6 IT and *Fezf2* in L5 IT (**Fig. 2c, node 7, Extended Data Fig. 4)**. *Nfia* and *Sox5*, which show strong enrichment in the nonIT lineage, are also enriched in L6 IT (**Extended Data Fig. 4, node 3).** Within upper-layer IT population, we observe separation of L2/3 IT and L4/5 IT subclasses around E18.5, with *Rorb*, *Rora* and *Tox* enriched in L4/5 IT and *Mdag1* and *Klhl1* in L2/3 IT (**Fig. 2c, node 8, Extended Data Fig. 4)**.

Within each glutamatergic subclass, cells continue to differentiate and diversify, giving rise to new cell types/clusters. We derived a cluster trajectory tree of all cell types and conducted DE gene analysis at each branching point (**Fig. 3a, Extended Data Fig. 5, 6**).

For the L5 ET subclass, clusters 371-373 (*Chrna6*) represent the most distinct subset^11,13,19^, emerging at P3 with specific expression of transcription factors *Pou6f2* and *Otx1* (**Extended Data Fig. 5**). Expression of marker gene *Chrna6* begins relatively late, around P9, and peaks in adulthood. Clusters 372 and 373 diverge from 371 after P21, with 373 specifically expressing *Hk2*. While *Fxyd6* is widely expressed in nonIT cells, it is downregulated in specific cell types, including clusters 372 and 373, after P21. Based on our trajectory analysis, *Chrna6*+ clusters 371-373 share a common lineage with clusters 365 and 366, with shared expression of *Kctd8* (**Extended Data Fig. 5**). We have identified multiple transcription factors potentially involved in regulation of different L5 ET clusters, including *Foxo1*, *Bmp5*, *Lhx2*, *Zfp804b* and *Erg*. There is no apparent spatial segregation of different L5 ET clusters in visual cortex.

The L5 NP subclass contains two clusters, 466 and 468, which are diverged around P3, with *Sv2c* and *Nxph2* enriched in each cluster respectively (**Extended Data Fig. 5**). *Nxph2*+ cluster 466 appears to be slightly deeper than cluster 468. The L6 CT subclass has three major clusters, 440, 439 and 437, diverging at E17 (**Extended Data Fig. 5**). Interestingly, *Nxph2*+ L6 CT cluster 440 is very distinct from the other L6 CT clusters but more related to L5 NP subclass based on trajectory analysis, with shared expression of transcription factor *Pou3f2* with L5 ET and L5 NP. The separation between L6 CT clusters 437 and 439 (the dominant L6 CT cluster, emerging as early as E13.5) is quite subtle transcriptomically, marked by enrichment of *Pantr1* and *Htr4*, respectively, but very distinct spatially: cluster 437 is clearly deeper than 439 and is co-localized with L6b cells (**Extended Data Fig. 5f**). *Pantr1*, a noncoding RNA gene adjacent to transcription factor *Pou3f3*, is absent in the deep L6 CT cluster 437 and L6b but present in all other more superficial nonIT clusters. In L6b subclass, two major clusters 427 and 428 diverge around P1, with transcription factors *Foxp2*, *Nr4a2* and *Id4* enriched in 427 and *Tox* enriched in 428 (**Extended Data Fig. 5**). There is no apparent difference in spatial distribution of these two clusters, but 427 is more closely related to L6 CT subclass transcriptomically. Overall, most of the nonIT clusters begin to diverge by P3 except for a few L5 ET clusters (372-373) that emerge at the onset of critical period (**Fig. 3a, Extended Data Fig. 5h**).

In the IT lineage, many clusters that split off early have distinct layer distribution (**Fig. 3a, Extended Data Fig. 6f,h**). For instance, in the L5 IT subclass, clusters 64 and 56 diverge around E17, and 64 is more superficial than 56. In the L4/5 IT subclass, clusters 100 and 73 diverge at P0, with 100 being more superficial than 73. In the L2/3 IT subclass, clusters 110 and 111 separate around P3, with 110 located more superficially than 111. Many genes show distinct layer distribution at early stages of IT cell type divergence (**Extended Data Fig. 6g**). This result indicates the cortex has more continuous and refined sublayer gradient than the classical 6-layer model, and these sublayers are specified by early postnatal age.

More clusters arise in later stage of development after eye opening, and these newer clusters usually have less distinct spatial distribution from sibling clusters. For example, L2/3 IT cluster 109 diverges from 110 at around P11 with increased expression of *Bdnf* and decreased expression of *Adamts2*, while cluster 118 further diverges from cluster 109 at P21 with increased expression of *Baz1a and Tnfaip6* (**Extended Data Fig. 6**). Spatially within L2/3, clusters 118 and 110 are located somewhat more superficially than 109 (**Extended Data Fig. 6f**). Recently, functions of related L2/3 IT cell types in the somatosensory cortex have been characterized, and *Baz1a*+ L2/3 neurons show strong functional connections with Sst inhibitory neurons and orchestrate local network activity pattern^63^. Furthermore, in the visual cortex, it has been found that vision selectively drives the specification of L2/3 glutamatergic neuron types, and dark rearing reduces the diversity of these L2/3 types^64^. We also observe late divergence of L4/5 IT clusters 101 and 82 from cluster 100 at P14 and P20 respectively, which display subtle difference in spatial distribution, with cluster 82 located more superficially than 100 while 101 deeper than 100 (**Extended Data Fig. 6**). There are also new cell types emerging for L5 IT and L6 IT subclasses, with L6 IT clusters 41 and 50 emerging from 37 around P12, cluster 52 emerging from 41 around P20, and L5 IT clusters 62 and 63 emerging from 56 and 64, respectively, around P19. Overall, the late-onset IT clusters emerge in mainly two waves - first after eye opening, and then at critical period (**Fig. 3a, Extended Data Fig. 6h**). Many activity-dependent genes are upregulated during this process, which is shown in greater detail in a later section.

Some genes that contribute to adult cell type specificity have interesting temporal dynamics during development (**Extended Data Fig. 6g**). For example, *Cd24a* is widely expressed in embryonic stages and turned off gradually since P9 in IT cell types. It is turned off completely first in L4/5 subclass by P14, then in most other cell types later around P17, but stays on till adulthood in L6 IT cluster 51. Similarly, *Lefty2* is turned on in L2-L5 IT cells around P2 but stays on in L5 IT cluster 68 much later than other cell types.

Overall, most nonIT clusters already exist before eye opening, except for a few *Chrna6*+ L5 ET clusters; in contrast, IT clusters continue to emerge from P11, around time of eye opening, to as late as P21, at the onset of critical period (**Fig. 3a**). This suggests that IT cells become molecularly distinct at embryonic stage and continue to diversify throughout the postnatal period.

### Developmental trajectories of glial cell types

Radial glia transition into gliogenesis starting at E15.5, as indicated by the increasing expression of *Tnc* (**Fig. 2, node 2**). *Slco1c1* and *Sparcl1* are turned on in RG at E17, with further activation in glioblasts (**Fig. 2d,f**). Glioblasts emerge at E15.5 and contain two clusters initially: Glioblast and a special population we refer to as Glioblast SVZ (**Fig. 3b**).

The Glioblast SVZ cluster shares expression of *Veph1*, *Tspan18*, *Tfap2c* and *Adamts18* with RG and shares expression of *Slco1c1* and *Tnc* with astrocytes (**Extended Data Fig. 7**). *Gja1* is turned on in this population later at ∼P0, and *Thbs4* is turned on at ∼P9. This cluster is mapped to the adult astrocyte cell types located in the subventricular zone (SVZ) bordering rostral dorsal striatum, part of the rostral migration stream (RMS)^15^. These SVZ-RMS astrocytes create a migration permissive environment by providing soluble and non-soluble cues to the newly formed, migrating olfactory bulb neurons. The Glioblast SVZ cluster is likely the precursor of SVZ-RMS astrocytes.

The Glioblast cluster is labeled by both oligodendrocyte markers *Olig1* and *Olig2,* and astrocyte markers *Tnc*, *Slco1c1* and *Egfr*, and gives rise to both astrocytes and OPCs/oligodendrocytes (**Fig. 2a,c, node 9, Fig. 3b**). This cluster quickly splits into clusters Glioblast Astro and Glioblast OPC, with enrichment of *Slco1c1*, *Aldoc*, *Id3* and *Pax6* in the astrocyte lineage and enrichment of *Dll1*, *Dll3*, *Ascl1* and *Erbb4* in the OPC/oligodendrocyte lineage (**Fig. 2c, node 9, Extended Data Fig. 4, 7**). Notch ligands *Dll1*, *Dll3* and *Ascl1* are expressed transiently and downregulated as the cells transition from glioblasts to OPCs, while *Erbb4* maintains its expression. It has recently been shown that Notch signaling plays a dual role in both promoting and inhibiting oligodendrogenesis to fine-tune regulation of oligodendrocyte generation^65^. In our dataset, *Dll1* and *Dll3* activation coincides with downregulation of astrocyte markers *Tnc* and *Slco1c1* precisely, suggesting Notch signaling can also be involved repressing astrocyte fate. *Sox9*, strongly expressed in RG and glioblasts, is downregulated in OPC and turned off completely after cells exiting OPC; in contrast, *Sox10* is activated at the end of glioblast stage and remains active throughout the developmental process of oligodendrocytes (**Extended Data Fig. 7g**). The OPC cluster 5271 shows strong expression of cell cycle genes *Mki67* and *Top2a*, indicating that these cells are still rapidly proliferating, and this cluster largely disappears by eye opening. Telencephalon spatial patterning transcription factors *Foxg1* and *Lhx2* are strongly expressed in RG, downregulated but maintained in astrocytes, down regulated more dramatically in OPC, and disappear in mature oligodendrocytes. This pattern can explain why spatial identity is maintained in astrocytes, but not in oligodendrocytes, as previously reported^15,33^.

During postnatal development, OPCs are predominant, but after P11 their proportion gradually decreases (**Extended Data Fig. 7h**). Committed oligodendrocyte precursors (COP) start to appear around P2, marked by downregulation of *Creb5*, *Etv5*, *Etv4*, *Sox9* and *Pdgfra* and upregulation of *St18*, *Bmp4*, *Enpp6* and *Plp1* (**Fig. 2c, node 10, Extended Data Fig. 4, 7**). Newly formed oligodendrocytes (NFOL) emerge around P11, coinciding with eye opening, while mature oligodendrocytes (MOL) appear around P12 and continue to increase their proportion and diversity until adulthood (**Extended Data Fig. 7h**). These results align with previous studies showing that neuronal activity influences OPC and oligodendrocyte proliferation, differentiation, and myelin remodeling^66^.

We identified four astrocyte clusters in this dataset (**Extended Data Fig. 7**). Cluster 5225 is the most dominant astrocyte cell type within the visual cortex. Clusters 5218 and 5219, with enriched expression of *Gfap*, *Myoc* and *Atoh8*, are interlaminar astrocytes (ILA) localized at the pia of cortex^15^. Cluster 5228 is a rare astrocyte cell type that is enriched in lateral cortex and cortical subplate (CTXsp) marked by *Thbs4* expression and is related to the SVZ-RMS astrocytes based on their transcriptomic profiles^15^. Our trajectory analysis indicates that cluster 5228 likely originates from the Glioblast SVZ cluster.

### Developmental trajectories of GABAergic neuron types

The earliest GABAergic cell populations emerge in visual cortex at E11.5 and express transcription factors *Dlx1*, *Dlx2*, *Ascl1* and *Gsx2*, which are required for specification of all GABAergic neurons in the subpallium^25,67,68^. Analyzing the trajectories of cortical GABAergic cells at the cluster level is challenging due to their long-distance migration patterns, particularly during early development, and their inherent heterogeneity. The transcriptional signals that differentiate these cell types are complex and continuous across multiple dimensions. These cells migrate from the ganglionic eminence along tangential paths, with some cell types being pre-specified before reaching the cortex. Since our postnatal data collection only includes GABAergic cells in the cortex, it is likely that some antecedent nodes in the trajectory paths are outside the cortex, thus missing in our dataset, and our trajectory analysis assuming all ancestral nodes are present in the current dataset could be misleading. Nonetheless, we still successfully inferred multiple trajectory paths with good confidence (**Fig. 2a, 3c,d, Extended Data Fig. 8, 9, Supplementary Table 5**).

At E11.5, we observed in the cortex the initial emergence of MGE GABAergic progenitors, the MGE GABA RG and MGE GABA subclasses, which progress to MGE GABA immature neurons at E14.5 (**Fig. 2b**). *Ascl1* and *Tead2* are strongly enriched in progenitor stage, followed by activation of *Lhx6*, *Nkx2-1* and *Lhx8*, which are key regulators of development of MGE-derived GABAergic neurons^67,69,70^ (**Extended Data Fig. 8**). *Nkx2-1* and *Lhx8* are transiently expressed, while *Lhx6* is expressed even in adulthood. We also observed expression of *Nfib* and *Sp9* in early stages of MGE cells, which slowly decrease and maintain low level expression in some adult cell types. While previous studies suggested that Sst and Pvalb cells may originate from different domains in MGE^71^, we do not see segregation of these two populations in the MGE RG and MGE subclasses, though subtle differential gene signatures might exist. Starting at E14.5, MGE cells gradually differentiate into two subclasses, Sst Gaba and Pvalb Gaba, with *Shisa6*, *Pou3f3*, *Npas1* and *Tox* enriched in the Sst subclass, and *Adamts17*, *Shisa9*, *Tafa2* and *Zfp804b* enriched in the Pvalb subclass (**Fig. 2a,c, node 11, Extended Data Fig. 4**). While *Sst* is expressed early in embryonic stages, *Pvalb* is not expressed until after eye opening (**Extended Data Fig. 8g**).

We identified three additional highly distinct MGE subclasses, the Sst Chodl and Pvalb chandelier subclasses emerging around P1, and the Lamp5 Lhx6 subclass emerging around P5 (**Extended Data Fig. 8h**). These subclasses probably have diverged from other MGE cell types before reaching the cortex. *Nfib*, *Sp9* and *Nkx2-1* are enriched in Pvalb chandelier and Lamp5 Lhx6 subclasses even in adulthood, while they are significantly downregulated during development in most other MGE cell types (**Extended Data Fig. 8g**).

Within the Pvalb and Sst subclasses, we identified five primary developmental trajectories for each (**Fig. 3c, Extended Data Fig. 8e**). Grouping of these GABAergic cell types by trajectories matches the definition of Morpho-electric and transcriptomic (MET) types previously categorized in mouse visual cortex using Patch-seq^16^, as shown in Extended Data Fig. 6 of our recent study^37^. Clusters within each trajectory group often split at late postnatal ages, especially during eye opening (at ∼P11) or critical period (at ∼P21), suggesting continued diversification.

Within the Pvalb subclass (**Fig. 3c, Extended Data Fig. 8**), out of the five Pvalb MET types, four are fast-spiking basket cells (or cells with related morphologies) located in different layers, and one (Pvalb MET 5) is the chandelier cells^16^. Our developmental group 1 with clusters 736 and 754 marked by *Gpr149*, and group 2 with cluster 741 marked by *Reln* and *Pdlim3*, both correspond to the Pvalb MET 3 type (in L5). Cluster 736 emerges from 754 at P19. Group 3 marked by *Tpbg* and *Calb1* contains clusters 742 and 752, with 742 corresponding to Pvalb MET 4 (in L2/3), and 752 diverging from 742 at P17 and corresponding to Sst MET 2. Group 4 with clusters 743, 744 and 747 (split at P11), enriched with *Sema3e*, *St6galnac5* and *Ptprk*, corresponds to Pvalb MET 2 (in L6). The *Th*+ Pvalb cluster 735 corresponds to Pvalb MET 1 (in L6). However, the developmental trajectory of cluster 735 (emerging at P1) appears highly ambiguous, with its closest antecedent being Sst cluster 758 (**Fig. 3c, Extended Data Fig. 8a-c**). This is consistent with our previous finding that the L6 *Th*+ Pvalb cells may be a transition type between Pvalb and Sst subclasses^13^. In addition, the Pvalb chandelier cluster 733 corresponds to Pvalb MET 5 (in L2/3).

Within the Sst subclass (**Fig. 3c, Extended Data Fig. 8**), there are 13 Sst MET types^16^. Besides the Sst Chodl subclass (cluster 859, corresponding to Sst MET 1, long-range projecting neurons), the five developmental groups also correspond to specific MET types^37^. All Sst clusters exhibit highly restrictive layer distribution. In group 1, clusters 757, 758 and 761 correspond to Sst MET 9-10, and clusters 811, 814, 818, 819 and 820 correspond to Sst MET 12-13. Sst MET 9-13 types are all L5/6 non-Martinotti cells. Group 1 is characterized by the presence of *Crh* and *Crhr2*, with significant enrichment of *St6galnac5* and *Ptprk*. In group 2, clusters 795 and 797 correspond to Sst MET 2 (L2/3 fast-spiking-like cells), and clusters 803 and 806 correspond to Sst MET 3-5 types (L2/3 and L5 fanning Martinotti cells). Group 2 is marked by *Cbln4* and *Calb2*, with enrichment of *Tox*. In group 3 marked by *Hpse*, clusters 792 and 793 correspond to Sst MET 8 (L4-targeting Martinotti cells), and clusters 799, 800 and 801 correspond to Sst MET 7 (L5 T-shaped Martinotti cells). In groups 4 and 5, clusters 779, 777 and 780 correspond to Sst MET 6 (also L5 T-shaped Martinotti cells). Groups 4 and 5 display similar transcriptomic profiles, sharing markers *Nr2f2* and *Myh8*, with *Pdyn* enriched in group 4 and *Kit* enriched in group 5. Interestingly, we previously found that the Sst MET 2 type (L2/3 fast-spiking-like cells) may be another transition type between Pvalb and Sst subclasses^13,16^, and Sst clusters 795 and 797 and Pvalb cluster 752 are all mapped to Sst MET 2, consistent with their relatedness in the transcriptomic space (Extended Data Fig. 8a-c).

Many Sst clusters emerge relatively late within each group, with late activation of key genes (**Fig. 3c, Extended Data Fig. 8**). For example, *Crh* is activated around P5 and *Crhr2* around P10. Trajectory analysis suggests that *Crhr2*+ clusters 811, 814, 818, 819 and 820 diverge from *Crh*+ clusters 757 and 758 around P7, with further divergence occurring after P19. In group 1, 757, 758 and 761 split at P20, 811 and 814 split at P12, and 818, 819 and 820 split at P21. In group 2, 795 is born around P14, while 797 and 806 diverge from 803 around P19. In group 3, all 5 clusters diverge from 792 at P19-21. In group 5, 777 splits from 780 at P20.

CGE-derived neurons emerge in cortex around E14.5, and they gradually split into Vip, Sncg and Lamp5 subclasses during E18-P1 (**Fig. 2a,b, 3d**). *Id4* is enriched in the Vip subclass, *Npas1* and *Synpr* are enriched in both the Vip and Sncg subclasses, *Bcl11b* is enriched in the Lamp5 subclass, and *Ptprm* and *Id2* are enriched in both the Sncg and Lamp5 subclasses (**Fig. 2a,c, node 12, Extended Data Fig. 4**).

In the Vip subclass, we identified five main developmental trajectories, with clusters within each group often split at late postnatal ages, suggesting continued diversification (**Fig. 3d, Extended Data Fig. 9**). Group 1, marked by *Crispld2* and *Mybpc1*, contains clusters 645, 646, 648 and 629 that are split from 645 at P21. Group 2, marked by *Rspo2* and *Rspo4*, contains cluster 627. Group 3, marked by *Chat* and *Npy2r*, contains the root cluster 641, plus 643 and 663 emerging at P11, 633 at P19, 638 at P23, and 640 at P56. Group 4, marked by *Sntb1*, contains clusters 662, 661, 660 and 639, with 639 emerging at P2, 662 at P9, 661 at P15, and 660 at P21. Group 5, marked by *Grin3a* and *Igfbp6*, contains two clusters, with 623 split from 624 at P23. Most Vip clusters are present in the L2/3 layer, except for cluster 639 enriched in the deep layers. Vip MET 1-5 types represent L2/3-5 bipolar or bitufted cells^16^. Most clusters in groups 1-3 are mapped to Vip MET 4 and 5, cluster 641 corresponds to Vip MET 2, and cluster 663 and group 4 clusters 660-662 all correspond to Vip MET 1 type^37^. Group 5 is a highly distinct Vip type with almost no *Vip* expression (**Extended Data Fig. 9e,g**) and no matching MET type^37^, suggesting that these cells were not sampled in the Patch-seq study.

The Sncg subclass has one main trajectory marked by *Plcxd3*, *Frem1*, *Egln3* and *Sncg*, with *Sncg* expressed the latest (**Fig. 3d, Extended Data Fig. 9**). Among the 3 Sncg clusters, cluster 676 gives rise to 682 and 673 at P11 and P20, respectively. These clusters all correspond to Sncg MET 2 type which is the main type for CCK+ basket cells^13,16^.

In the Lamp5 subclass, we identified four main developmental trajectories (**Fig. 3d, Extended Data Fig. 9**). Group 1, marked by *Egln3*, *Col14a1* and *Fbn2*, contains clusters 719 (emerging at P2), 720 (P12) and 722 (P21). Group 2 (clusters 716, 717 and 718, split from 718 at P25) and group 3 (clusters 706 and 708, split at P28) are very similar, marked by shared expression of *Dock5* and *Ndnf*, with 708 as the root cluster and 718 split from 708 at P11. Group 4, containing cluster 709 and enriched in *Lsp1* and *Cemip*, shares expression of *Tox2* and *Sv2c* with groups 2 and 3 and emerge at P1 along with cluster 708. Groups 2-4 clusters 706, 708, 709 and 718 all correspond to Lam5 MET 1 type which represents the L1-5 neurogliaform cells^13,16^. No MET type matches group 1 clusters, suggesting that these cells were not sampled in the Patch-seq study. The Lamp5 cells are predominantly found in L1, whereas cluster 709 also includes neurons located in the deeper layers.

Taken together, the above results reveal high degree of correspondence between transcriptomic trajectories and morpho-electrical properties of highly specific Sst and Pvalb GABAergic neuronal types, as well as major Vip, Sncg and Lamp5 GABAergic types. Most of these GABAergic MET types correspond to distinct trajectory paths with early developmental origins, with late arising clusters (T-types) contributing to diversification within each MET type. A prominent exception of this is the many Sst Martinotti cell and non-Martinetti cell MET types, each of which corresponds to a specific set of Sst clusters emerging in late postnatal development stages, resulting in several MET types with different axon-targeting specificity contained within a trajectory group, suggesting that the extensive transcriptomic cell type diversification of Sst neurons is associated with the formation and refinement of the intricate local circuits between Sst types and other inhibitory and excitatory neuron types^18,72^.

### Gene co-expression modules

Gene modules can provide a more integrated description of complex biological processes such as cell type diversification than the expression pattern of individual genes alone. We attempted to identify gene modules across ages and cell types. We first identified the key sources of transcriptomic variation across time points within each class. Next, we identified DE genes that are linked to cell types at class level within a time point. The DE genes across time points and cell classes were then clustered, resulting in 96 modules that consist of genes that co-express across different cell classes and ages, ranging from 6 to 67 genes in each module (**Figure 4, Supplementary Table 6**). Using gene ontology (GO) term enrichment analysis, we assigned biological processes to most modules. The roles of these modules cover several key aspects of brain development including cell fate determination, cell division, synapse function, immune function and myelination. Not surprisingly, modules related to vascular or immune activity are enriched in endothelial and microglial clusters, respectively. Similarly, some modules linked to oligodendrocyte and astroglial function are specific to those cell types.

**Figure 4.**
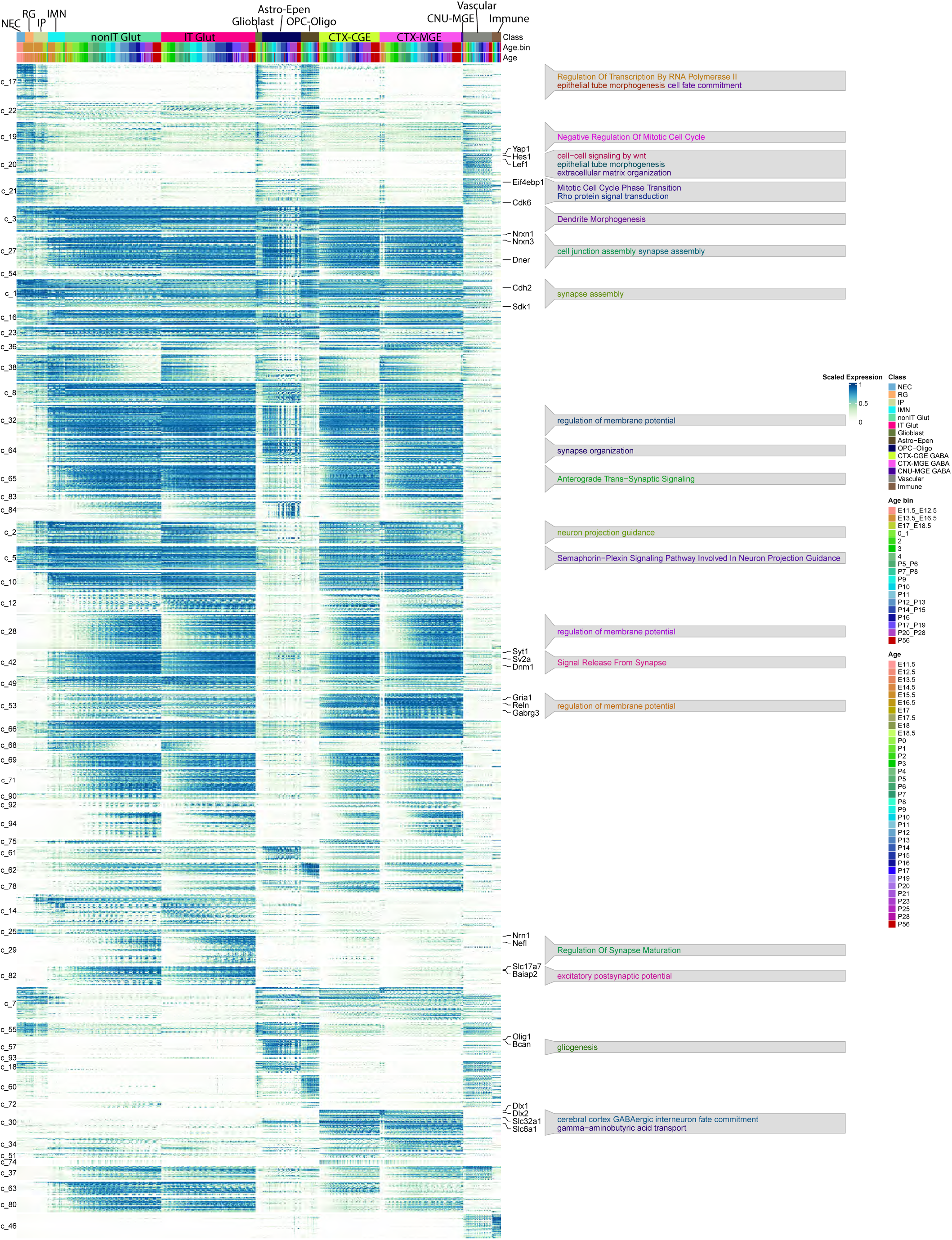
Gene co-expression modules across cell types and ages during development. Module gene expression heatmap of each class in the developing taxonomy. Clusters are organized by gene co-expression modules shown as color bars on the right side of the heat map and by age bin and class on the top of the heatmap. Module score is the mean expression of genes in the module within each cluster. Significant GO enrichment terms of gene modules are highlighted.

While gene modules that are enriched in progenitors are linked to broad developmental processes such as multicellular organism growth, developmental induction, neurogenesis and neuronal migration. Distinct lineage-specific gene modules start to emerge at E13.5 (**Figure 4**).

Class modules c_77, c_81 and c_89 are enriched in NECs and RGs, with a high proportion of cell proliferation-related genes (**Supplementary Table 6**). Proliferation genes in c_ 89 are expressed in S phase (associated with DNA replication), while those in c_77 and c_81 are mostly expressed in G2/M phase (associated with mitosis). Module c_4 is highly expressed in IPs, and genes in this module are related to cell fate commitment, neuron differentiation and the Notch signaling pathway.

Module c_6 is predominantly active in NECs, RGs, IPs and IMNs and is linked to cell migration and presynaptic assembly (**Supplementary Table 6**). Gene products of *Mdga1*, *Efnb1* and *Gpc4* are involved in cell-cell interaction thereby mediating the assembly of presynaptic terminals. For example, *Mdga1* regulates the interaction of *Nlgn2* with neurexins, which are presynaptic adhesion molecules and *Efnb1* mediates EphB-dependent presynaptic development via PDZ-binding domain-dependent interaction with syntenin-1^73–75^.

The idea that the temporal dynamics of module activity in progenitor cells during development can inform analyses of cortical cell type specification is particularly exemplified by module 18 (**Figure 4, Supplementary Table 6**). This module is active in radial glia throughout development and gradually becomes highly restricted to glial subtypes. GO analysis of module c_18 indicates these genes play a role in glial cell differentiation and negative regulation of Wnt signaling. Wnt is a key regulator of neuronal differentiation in the nervous system, controlling the development of neuronal circuits. Consistent with a role for c_18 in gliogenesis, genes in this module – *Notch1*, *Metrn*, *Ntnt1* – have reported roles in both glial cell differentiation and axonal network formation during neurogenesis^76,77^.

As differentiation proceeded, there is a shift towards neuron projection, synapse function, ion transportation, and myelination, reflecting developmental maturation.

We examined the ability of gene modules to represent neuronal cell type identities. Our analyses revealed three class modules (c_14, c_82 and c_29) that are active in glutamatergic neurons during development (**Figure 4, Supplementary Table 6**). Genes in c_14 are enriched in IPs, nonIT, and IT neurons, and are linked to pan-glutamatergic cell type development. Genes in this module are associated with axonogenesis, neural precursor cell proliferation, neuron migration, neurotransmitter uptake, neuropeptide signaling pathway, and neuron projection fasciculation. Class modules c_82 (associated with dendritic spine, distal axon and synaptic signaling) and c_29 (associated with fear response and synaptic regulation) are enriched in the IT and nonIT neurons. All three modules increase in activity over the developmental time course, with c_29 emerging later than the other two, at P11.

To further examine how well our module analysis can inform cortical cell fate specification, we focus on subclass modules. Within each of the classes, subclasses share specific gene modules that are linked to the general specification of the cell class, and each subclass has its unique temporal gene modules (**Extended Data Fig. 10, 11, Supplementary Table 6**). For example, the shared subclass modules within the IT class at early ages are enriched for genes involved in neuronal cell death (necroptosis, apoptosis), growth factor signaling, and axon guidance, whereas at later ages the shared gene modules in the IT class are enriched for genes involved in metal ion transport (zinc, sodium, potassium) and intracellular protein transport. Similarly, in the nonIT class the shared gene modules are enriched for genes involved in mitosis, cell migration, negative regulation of projection development at the early stages, and cell-cell adhesion and synaptic vesicle exocytosis at late stages.

Within the nonIT class, modules for the L6b subclass (snonIT_14 and 19) contain genes including *Kcnj5, Gng4, Lpar1 and Drd1* which are involved in the activation of G-protein gated potassium channels^78^. Modules in the L5 NP subclass (snonIT_25 and 29) contain genes like *Chrm2*, *Camk2s*, *Grm4* and *Trpc3* which are linked to GPCR signaling at the synapse. The module in the L5 ET subclass (snonIT_34) contains genes like *Epha6, Reln, Itgav* and *Slit2* which are linked to axon guidance^79,80^.

### Dynamic changes during eye opening

The above trajectory analysis reveals increased cell type diversity in the visual cortex from early to late developmental stages, as well as extensive transcriptional heterogeneity within a cell type, as shown in the single cluster of NEC and that of RG (**Fig. 2f**). As a means to quantify diversity and heterogeneity, we plotted the total number of clusters and the total number of subclusters within each cluster across synchronized ages (**Fig. 2g**). We find that the number of clusters continues to increase with time, with jumps at P11-13 and P19-21. In contrast, at subcluster level, there are several bouts of increased subcluster numbers, indicating heightened heterogeneity of transitional cell subtypes or cell states at different time periods that are associated with specific developmental events, such as neurogenesis (E13.5-P1), axon growth and synapse formation (P5-P9), eye opening (P12-P15), and critical period of experience-dependent plasticity (P20-P28). The emergence of new cell types following eye opening (**Fig. 3, Extended Data Figures 5-9**) spurred our exploration into the molecular characteristics preceding and following this event. Previous studies showed that vision is required for the development of cortical circuitry during the critical period for ocular dominance plasticity (P21–P38)^32^.

We first compared the overall transcriptional profiles and conducted DE gene analysis of cell types before and after eye opening within each subclass or cluster, combining scRNA-seq data during P7-P10 for before eye-opening period and during P11-P15 for after eye-opening period (**Fig. 5a-d, Supplementary Table 7**). Genes with |log_2_(FC)| > 1 and FDR < 0.05 are considered having significant expression changes. Remarkably, all neuronal and non-neuronal subclasses have diverse transcriptional changes and there are genes turned on or off for each subclass and cluster (**Fig. 5e,f, Supplementary Table 7**). On average, glutamatergic subclasses, including both IT (∼1,600-2,000 DE genes for each subclass) and nonIT (∼1,200-1,800 DE genes for each subclass), have more DE genes than GABAergic subclasses, except for Pvalb. While most neuronal subclasses have more up-regulated genes than down-regulated genes, all non-neuronal subclasses have more down-regulated genes. For example, microglia have 1,797 down-regulated genes but only 80 up-regulated genes (**Fig. 5e**).

**Figure 5.**
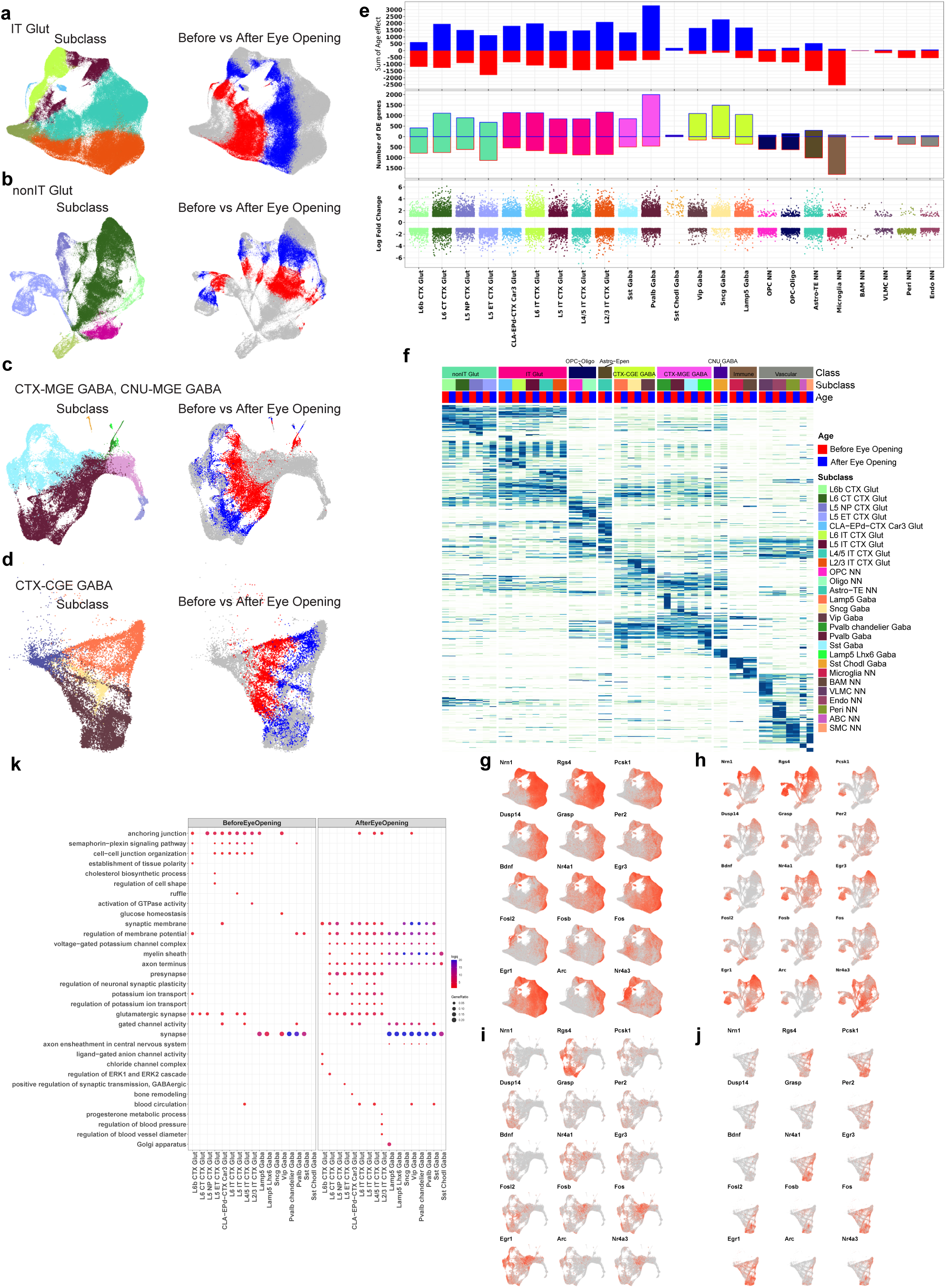
Dynamic gene expression changes before and after eye opening. (a)-(d) UMAP representation of the IT Glut and IMN IT cell types (a), nonIT Glut and IMN nonIT cell types (b), CTX-MGE GABA and CNU-MGE GABA cell types (c) and CTX-CGE GABA (d) colored by subclass and age (before eye opening: P7-10; after eye opening: P11-15). **(e)** DE genes between before and after eye-opening age points for all cell subclasses. Bottom, log2 fold change of each DE gene. Middle, number of DE genes up or down regulated during eye opening. Top, sum of log2 fold changes of all DE genes up or down regulated during eye opening. **(f)** Heat map showing the expression of specific DE genes in each subclass before and after eye opening. **(g-j)** Expression changes of IEGs on IT (g), nonIT (h), CTX-MGE and CNU-MGE (i) and CTX-CGE (j) UMAPs. **(k)** GO enrichment dot plot showing example significant top GO terms before or after eye opening in each neuronal subclass. Dot size and color indicate gene ratio (the percentage of genes that are present in a GO term compared to the total number of genes in that category) and significance (-log adjP value), respectively. Max gene ratio was set to 0.2 and max significance was set to 20.

These DE genes (**Supplementary Table 7**) include many immediate-early genes (IEGs), such as *Fos*, *Fosb*, *Fosl2*, *Egr1*, *Arc*, *Bdnf*, and *Nr4a3* (**Fig. 5g-j**), consistent with previous findings^81^. These IEGs often have different temporal patterns among different subclasses, suggesting that different IEGs may have different effects in cortical microcircuits.

We also observed cluster-level transcriptional changes with eye opening. For example, *Pdlim1*, which encodes a protein involved in AMPA receptor trafficking and regulates synaptic plasticity, shows significant enrichment following eye opening in some L5 IT, L5 ET and L6 IT clusters (**Supplementary Table 7**). In the Sst (758 and 811) and Vip (624 and 645) clusters we observed increased expression of *Crh* after eye opening (**Supplementary Table 7**). *Crh* encodes the stress hormone corticotropin-releasing hormone and signals through its receptor *Crhr1* which is enriched in both IT Glut (L6 IT 37 and 51) and nonIT Glut (L6 CT 437, 439 and 440; L5 ET 368) clusters, suggesting that Crh from Sst and Vip interneurons might modulate the excitability of glutamatergic neurons. Additionally, in Sst (792, 803, 811 and 859) and Pvalb (742 and 754) clusters, there is an enrichment of *Crhbp*, a gene encoding Crh-binding protein, a secreted factor that negatively regulates Crh signaling^82^.

From GO term enrichment analysis (**Fig. 5k**), we observe strong enrichment in the semaphorin-plexin signaling pathway (*Flna*, *Plxna3*, *Sema6c*, *Sema4g*, *Met*) and anchoring junction (*Traf4*, *Gjc1*, *Wtip*, *Tgfbr1*, *Pard6g*, *Pard3*) in downregulated genes in glutamatergic neurons. Conversely, genes associated with presynapse (*Mt3*, *Kcnab2*, *Kcna1*, *Cntnap1*), synaptic membrane (*Eno1*, *Kcna1*, *Cntnap1*, *Mpp2*, *Kcna2*), potassium ion transport (*Tmem38a*, *Kcna2*, *Kcnk1*, *Amigo1*), regulation of membrane potential (*Scn2b*, *Got1*, *Glrx*), and regulation of neuronal synaptic plasticity (*Vgf*, *Synpo*, *Neurl1a*, *Arc*, *Egr1*) are significantly upregulated in glutamatergic neurons.

We also observe enrichment of specific GO terms in specific subclasses. For example, after eye opening, L2/3 IT neurons show enrichment in regulation of blood pressure and regulation of blood vessel diameter (*Atp1a1*, *Adrb1*, *Pparg*, *Cd34*), while L4/5 IT and L6 IT neurons, as well as Sst and Vip interneurons, show enrichment in blood circulation (*Tmem38a*, *Slit2*, *Sema3a*, *Rgs4*)^83^. Interestingly, we observed that even when the same cell type is involved in the same biological process at different time points, the specific genes involved can vary. For example, L4/5 IT also shows enrichment in blood circulation before eye opening but with a different set of genes enriched during this period (*Vegfb*, *Ptpro*, *Ptger3*, *Gjc1*).

After eye opening, genes associated with myelin sheath are broadly enriched across different neuronal subclasses, in all IT subclasses except for L4/5 IT (*Atp1a2*, *Cntnap1*, *Kcnj11*, *Ldhb*, *Omg*), in L6 CT (*Nefl*, *Tppp*, *Cntnap1*, *Nefm*), and in all GABAergic subclasses (*Tppp*, *Thy1*, *Pgam1*, *Nsf*). In OPC-Oligo, *Rpl* and *Rps* genes which are associated with translation are enriched before eye-opening, and genes associated with regulation of neuronal synaptic plasticity (*Kcnj10*, <u>Ncdn</u>, *S100b*, *Camk2a*) and myelination are enriched after eye opening^84^ (**Supplementary Table 7**).

### Epigenomic chromatin accessibility landscape across developmental trajectories

In addition to the scRNA-seq dataset, we also collected the Multiome dataset that provides both chromatin accessibility profile and transcriptomic profile for each single nucleus. The dataset contains a total of 378,541 nuclei from 41 libraries collected at embryonic time points E13.5, E15.0, E15.5, E16.0, E17.0, E17.5, E18.0, and postnatal time points P0, P2, P4, P5, P8, P9, P11, P14, P56 (**Supplementary Table 1**). We integrated the Multiome snRNA-seq dataset with the scRNA-seq dataset using scVI^85^ (**Methods**) and obtained the transferred cell class, subclass and cluster labels from the reference scRNA-seq atlas for each nucleus (**Supplementary Table 8**). Multiome nuclei with poor mapping probabilities were removed from downstream analysis, which could be attributed to either lower-quality transcriptomes or presence outside the visual cortex, with a total of 194,545 Multiome nuclei remaining after filtering. Due to sparser postnatal time points for Multiome compared to scRNA-seq, we combined the time points into age groups E11.5_E12.5, E13_E16.5, E17_E18.5, P0_P3, P4_P6, P7_P10, P11_P15, P20_P28, and P54_P68, which are consistent with the synchronized age bins. Note that age groups E11.5_E12.5 and P20_P28 do not contain any Multiome samples. The UMAPs based on the integrated scVI latent space show great intermixing of the scRNA-seq and snRNA-seq data, and clear delineation of subclasses and age groups that are consistent between the two datasets (**Fig. 6a-c**).

**Figure 6.**
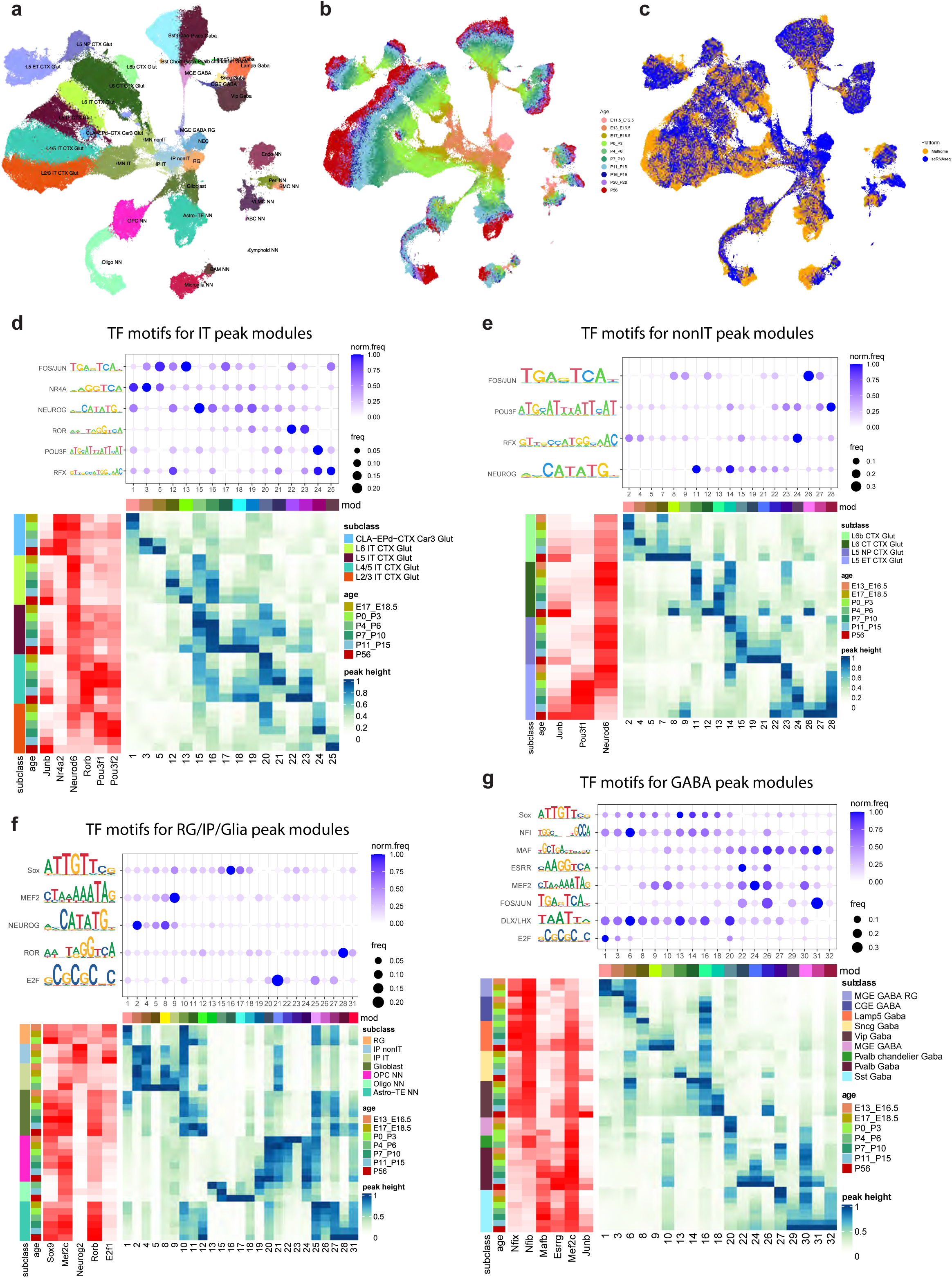
Integration of scRNA-seq and Multiome data and identification of transcription factor regulators for cell-type specific epigenomic dynamics. (a-c) UMAP representation of scRNA-seq and Multiome cells in the integrated space, colored by subclass (a), age group (b), and modality (c). The scRNA-seq cells shown in the UMAP are the subsampled ones (up to 200 cells per cluster) used for scVI integration. **(d-g)** Transcription factor motif enrichment for chromatin accessibility peak modules with different cell type and temporal specificities in IT (d), nonIT (e), RG/IP/Glia (f), and GABA (g). Within each panel, the dot plot at the top shows the average motif frequency for each peak module, dot size indicates the frequency, and color corresponds to the frequency normalized for each motif with maximum of 1. The large heatmap at the bottom shows the average accessibility for each peak module (in columns) across each subclass-by-age group (in rows). Accessibility values are normalized per peak module with 1 indicating the maximum value, and 0 indicating no accessibility. The heatmap at the left shows the average expression of specific transcription factors belonging to the motif families across each subclass-by-age group. The values are normalized per gene with 1 indicating the maximum value, and 0 indicating no expression.

We called chromatin accessibility peaks (total 958,146 peaks) using ArchR^86^ based on pseudo-bulk sets composed of mapped subclasses, clusters, and categories defined by both subclass and age group (**Methods**). We kept only the subclass-by-age-group categories with more than 50 cells (nuclei). We then performed pairwise differentially accessible (DA) peak analysis between all subclasses, and between all subclass-by-age-group categories using Chi-squared test (**Methods**). To study the peaks involved in regulation of cell types and their temporal dynamics, we selected DA peaks within each age group across different subclasses and pooled all DA peaks across all the age groups. We then identified peak modules with similar subclass specificity and temporal patterns among the DA peaks based on their average accessibility across subclass-by-age-group categories (**Methods**). We also computed DE genes across subclass-by-age-group categories. To associate peak accessibility with gene expression, we identified all the DA peak and DE gene pairs such that the DE gene is within a 5-Mb window centered at the DA peak and the corresponding gene expression and peak accessibility has a correlation greater than 0.5. For visualization purpose, for each peak module, we selected the top 500 peak-gene pairs with the strongest correlations (**Supplementary Table 9**).

We first applied this approach to study the subclass specificity within the IT Glut and nonIT Glut classes separately, starting from E17 (**Extended Data Fig. 12**). The reason to separate glutamatergic cells into these two populations is that many genes are re-used to specify different cell types in these two populations, introducing additional complexity for interpretation. For the IT Glut class, we identified early and late peak-gene pairs for each subclass (**Extended Data Fig. 12a**). For example, *Nr4a2* is a transcription factor highly specific to CLA-EPd-CTX Car3 Glut subclass that turns on early at E17_18.5 and stays on in adulthood. We identified peaks associated with *Nr4a2* that have the same subclass specificity but become weaker over time. *Nr2f2* and *Car3* are two other genes specific to this subclass, but these two genes have much stronger expression in late developmental stages, especially *Car3*, whose expression peaks at adult stage and corresponding peaks show similar temporal patterns. For L6 IT, we found *Nr4a3* gene expression and peak accessibility to turn on relatively early around P0 and peak around P10, and *Sema3e* peak accessibility and gene expression are strongest in adult stage. *Fezf2* has the strongest expression and peak accessibility in L5 IT, and its expression and peak accessibility turn on early and decrease in adulthood, while L5 IT marker *Deptor* turns on late after P10, and *Etv1* turns on later than *Fezf2* but earlier than *Deptor*. For L4 IT subclass, *Whrn* turns on earlier than *Pamr1*. *Pou3f2* and *Pou3f1* are widely expressed in IT cells, but have much stronger expression in upper layer neurons, and both of their expression decrease after P11. *Stard8* turns on relatively late in L2/3 IT cells specifically. There is no prominent peak module specific for L2/3 IT at early stages. We also observed a distinct peak module that is shared by L2/3, L4/5 and L5 IT neurons that peak at P7_P10, with *S100a10* as an exemplary gene. There is a similar gene module (exemplified by *St6galnac5*) shared by L5 and L6 IT subclasses that also peak at P7_P10, but the corresponding accessibility profiles have less specific temporal pattern. Overall, we find that cell type specific transcription factors tend to turn on early, while other functional genes turn on late.

We conducted similar analysis for the nonIT Glut class (**Extended Data Fig. 12b**). For L6b subclass, we identified *Hs3st3b1* as an early marker, and *Moxd1* and *Cplx3* turn on late, while *Nxph4* expression remains relatively stable in development. Similarly, *Syt6* and *Arhgap25* are the early and late L6 CT markers, respectively, *Tshz2* and *Vw2cl* as the early and late L5 NP marker respectively, and *Pou3f1* and *Lratd2* as early and late L5 ET markers. These marker genes all have matching chromatin accessibility profiles with similar subclass and temporal specificities. There are also other peak modules that are specific, but shared by multiple subclasses, e.g., module 8 is shared between L6b and L6 CT, module 9 is shared between L6b and L5 ET, module 22 is shared by L5 NP and L5 ET, and module 26 is shared by L6 CT and L5 ET. As in the IT population, we also observed multiple peak modules (13, 15, 24) with strongest activities in P7_P10. It is interesting that these peak modules are significantly more distinct between different subclasses than those for the IT Glut class, which are usually shared by multiple subclasses.

The GABAergic and glia populations show similar results (**Extended Data Fig. 13**). For each subclass, we identified early and late gene markers and corresponding accessibility peaks. Many well-recognized GABAergic subclass markers such as *Lamp5*, *Sncg*, *Vip* and *Pvalb*, and glia markers such as *Mbp* and *Apq4*, along with their associated accessibility peaks, are all turned on relatively late, except for *Sst* and its associated peaks and transcription factors which are turned on in embryonic stages.

### Identifying transcription factor regulators for cell-type specific epigenomic dynamics

To identify the potential transcription factor (TF) regulators for each peak module with different cell type specificity, we performed differential TF motif analysis between peak modules using all pairwise comparison (**Methods**). For the TF motifs that appear as significant in any pairwise comparison, we plot the motif presence frequencies across all the peak modules together with each module’s average subclass and temporal accessibility pattern (**Fig. 6d-g**). The motifs are typically described at the level of TF family instead of specific family members as their DNA binding motifs tend to be highly similar to each other. On the other hand, we can usually identify specific TF members within the family that have consistent gene expression patterns to narrow down the potential regulators.

For the IT Glut class, we identified NR4A motif to be enriched in both early and late Car3 specific peak modules, consistent with the specific *Nr4a2* expression in the Car3 subclass (**Fig. 6d, Extended Data Fig. 12a**). Additionally, the bHLH neurogenic motif NEUROG, shared by TFs *Neurog1*, *Neurog2*, *Neurod1*, *Neurod2*, *Neurod4*, and *Neurod6*, was found to be enriched in peaks specific to deep layer IT subclasses (**Fig. 6d**). While all these TFs are highly expressed in the IP populations, *Neurog1*, *Neurog2* and *Neurod4* expression diminish after the IP stage, while *Neurod1*, *Neurod2*, and *Neurod6* persist in adult neurons, albeit at weaker levels ((**Fig. 2c,d, 6d,f, Extended Data Fig. 4**). Postnatally, *Neurod1* expression is confined to the upper layer, while *Neurod6* exhibits greater expression in the deep layer neurons (**Fig. 6d**). These results suggest a potential role for *Neurod6* in regulating deep layer IT cell types. We also identified enrichment of the ROR motif in peak modules specific to the L4/5 IT subclass, which aligns with the specific expression of *Rorb* in this cell type (**Fig. 6d**). We also found enrichment of the POU3F motif in the L2/3 IT subclass, consistent with the stronger expression of *Pou3f1* and *Pou3f2* in the upper layer neurons, and their documented regulatory roles^87^ (**Fig. 6d, Extended Data Fig. 12a**). Furthermore, we detected enrichment of RFX motif in the L2/3 IT subclass, although we didn’t identify RFX members that have similar cell type specificity (**Fig. 6d**). It is plausible that they may serve as co-factors by recruiting other transcription factors to activate target sites. Finally, we observed enrichment of FOS/JUN AP1 motif in all the peak modules in late developmental and adult stages across all IT subclasses, indicating their roles in neuron maturation and activity^88^ (**Fig. 6d**).

For the nonIT Glut class, we found significant enrichment of the POU3F motif in the L5 ET subclass, particularly in the peak module that is activated early and persists till adulthood (**Fig. 6e**). This finding is consistent with the specific expression of *Pou3f1* in L5 ET, suggesting *Pou3f1* as a key regulator of L5 ET. We also found enrichment of RFX motif in L5 ET (**Fig. 6e**). However, similar to the L2/3 IT subclass, the specific member of the family involved remains unclear. We found enrichment of the NEUROG motif in the L6 CT and L5 NP subclasses (**Fig. 6e**). Notably, *Neurod6* expression is much higher in L6 CT and L5 NP compared to L5 ET and L6b, indicating its potential regulatory role for these cell types. Similar to IT cells, we also found enrichment of FOS/JUN motif in peak modules that are activated late, especially in the L5 ET subclass (**Fig. 6e**).

For RG, IP, and glia populations, we found enrichment of SOX motif in the peak modules specific to the Oligo subclass, which is the most different from the OPC subclass (**Fig. 6f**). *Sox10* is a transcription factor specific to the oligodendrocyte lineage, but it is uniformly expressed in OPC and Oligo subclasses (**Extended Data Fig. 7, 13b**). *Sox8* has similar expression as *Sox10* in oligodendrocytes but is also expressed in RG, gliobast and astrocytes. *Sox9*, is widely expressed in RG, glioblast, astrocytes and OPC, but is turned off in Oligo as the oligodendrocytes mature (**Fig. 6f, Extended Data Fig. 4, 7**). *Sox8*, *Sox9* and *Sox10* are all members of the SOXE group TFs. They play critical roles in development across diverse biological processes including gliogenesis, chondrogenesis, sex determination, as well as pancreatic, skin and kidney development^89^. They often function as dimers, and individual SOXE mutants often have less severe phenotypes than double or triple SOXE mutants^90,91^. *Sox9* is known to function either as an activator or a repressor depending on the partner factors and subsequent recruitment of either co-activators or repressors^92^. *Sox9* could potentially function in similar manners by first promoting glial fate^93^, then working with *Sox8* and/or *Sox10* to specify OPC/oligodendrocyte lineage, and finally repressing oligodendrocyte maturation till further developmental cues. Interestingly, *Sox10* is reported to repress *Sox9* expression by upregulating miR335 and miR338, which in turn downregulate *Sox9* protein level^94^. We also found enrichment of MEF2 and NEUROG motifs in IP populations, consistent with their roles in neurogenesis (**Fig. 6f**). ROR motif is enriched in astrocyte specific peak modules, in particular the mature astrocyte peak module, consistent with the increased expression of *Rora* and *Rorb* in astrocyte development (**Fig. 6f, Extended Data Fig. 4**). For example, *Rorb* has been reported to promote astrocyte maturation in culture^95^. Finally, we observed enrichment of E2F motif in peak modules that are enriched in OPC, IP, RG, glioblasts and astrocytes, in complementary patterns (**Fig. 6f**). E2F TFs are major regulators of cell cycle and cell proliferation, consistent with the fact that the enriched populations are nearly all proliferating cell types.

For GABAergic neurons, we observed enrichment of NFI (Nuclear Factor I) motif in peak modules specific to the CGE class, consistent with the specific expression of *Nfib*, *Nfix* and *Nfia* in this population, implying regulatory roles of NFI TFs in specifying the CGE lineage (**Fig. 6g, Extended Data Fig. 4**). Similarly, we identified enrichment of MAF motif in peak modules specific to the MGE class (**Fig. 6g**). *Mafb* has very specific expression in this population and is known to regulate MGE interneuron fate and function^96^. Interestingly, there is enrichment of ESRR motif in peak modules specific to the Pvalb subclass, especially at late developmental and adult stages (**Fig. 6g**). *Esrrb* has weak but specific expression for Pvalb neurons, especially in late developmental stages, while *Esrrg* has stronger expression in Pvalb neurons but with less specificity. It is plausible that estrogen-related receptor signaling pathway is involved in regulating Pvalb neuron maturation. We also identified depletion of MEF2 motif in peak modules specific to early MGE GABA RG subclass, consistent with *Mef2c* expression pattern, and its role in neuronal differentiation in general^97^ (**Fig. 6g**). In contrast, there is enrichment of E2F motif associated with cell proliferation in peak modules specific to the early MGE GABA RG subclass (**Fig. 6g**). Finally, like the IT and nonIT Glut cells, there is enrichment of FOS/JUN motif in peak modules that are activated in Pvalb and Sst subclasses in late developmental and adult stages (**Fig. 6g**).

### Genes regulated by multiple peaks with distinct temporal and cell-type specificity

When we compared peak accessibility with corresponding gene expression, we observed greater cell type and subclass specificity in the epigenomic data than in the transcriptomic data in many cases. While this could be because modules are defined in epigenomic space first, but based on a few examples we studied, it can also be attributed to the fact that expression of the same gene is controlled by multiple chromatin accessibility peaks with different cell type and temporal specificities.

For example, we were surprised that we did not find strongly correlated peak and gene pairs for *Cux2*, one of the most well studied transcription factors regulating the cortical cell type development. *Cux2* has complex cell-type and temporal expression pattern during cortical development (**Fig. 7a, Extended Data Fig. 4**). In glutamatergic cells, it is expressed in IP cells first, then more restricted to IMN IT cells, then further restricted to L2/3 IT, L4/5 IT and Car3 populations. It is also expressed in GABAergic (mostly MGE) cells and weakly in OPCs. To understand the overall epigenomic landscape of the *Cux2* gene, we extracted all the peaks located within the *Cux2* gene body (193 Kb) and 50 Kb upstream of *Cux2*’s main transcription start site. We then focused on the peaks that show differential accessibility between different subclasses within each age group. We observed strikingly complex accessibility patterns of different *Cux2* peaks (**Fig. 7a**), with distinct peak modules that are specific to IP (modules 1-3), IMN IT and upper layer IT cells (modules 4-6), Car3 cells (module 7), shared by L2-4 and Car3 cells (modules 8-9), specific to early L2/3 (module 10), shared by OPC and MGE (module 11), shared by IP, OPC and MGE (module 12), or specific to MGE (modules 13-16). We labeled specific peaks with distinct patterns and highlight them both in the heatmap (**Fig. 7a**) and in the cell type genomic tracks (**Fig. 7b**). Most of the peaks present in early-stage RG and IP populations disappear in adulthood, except those that are present near the promoter, or widely accessible. The accessibility of peaks in the promoter area overall shows strong consistency with RNA expression across all the cell types under study, while the peaks in more distal areas show accessibility in a highly cell-type and temporal specific manner. To study the subtler temporal progression, we examined the expression of *Cux2* gene and accessibility of specific peaks at the single cell level (**Fig. 7b,c**). Peak 1 is specific to IP, Peak 2 to IMN IT and L2-4, Peak 8 to MGE (decreasing over time), Peak 5 to L2-4 (increasing over time), Peak 10 mainly to IMN IT, and Peak 15 specific to Car3 and surprisingly in Microglia (although we don’t see expression of *Cux2* gene in Microglia).

**Figure 7.**
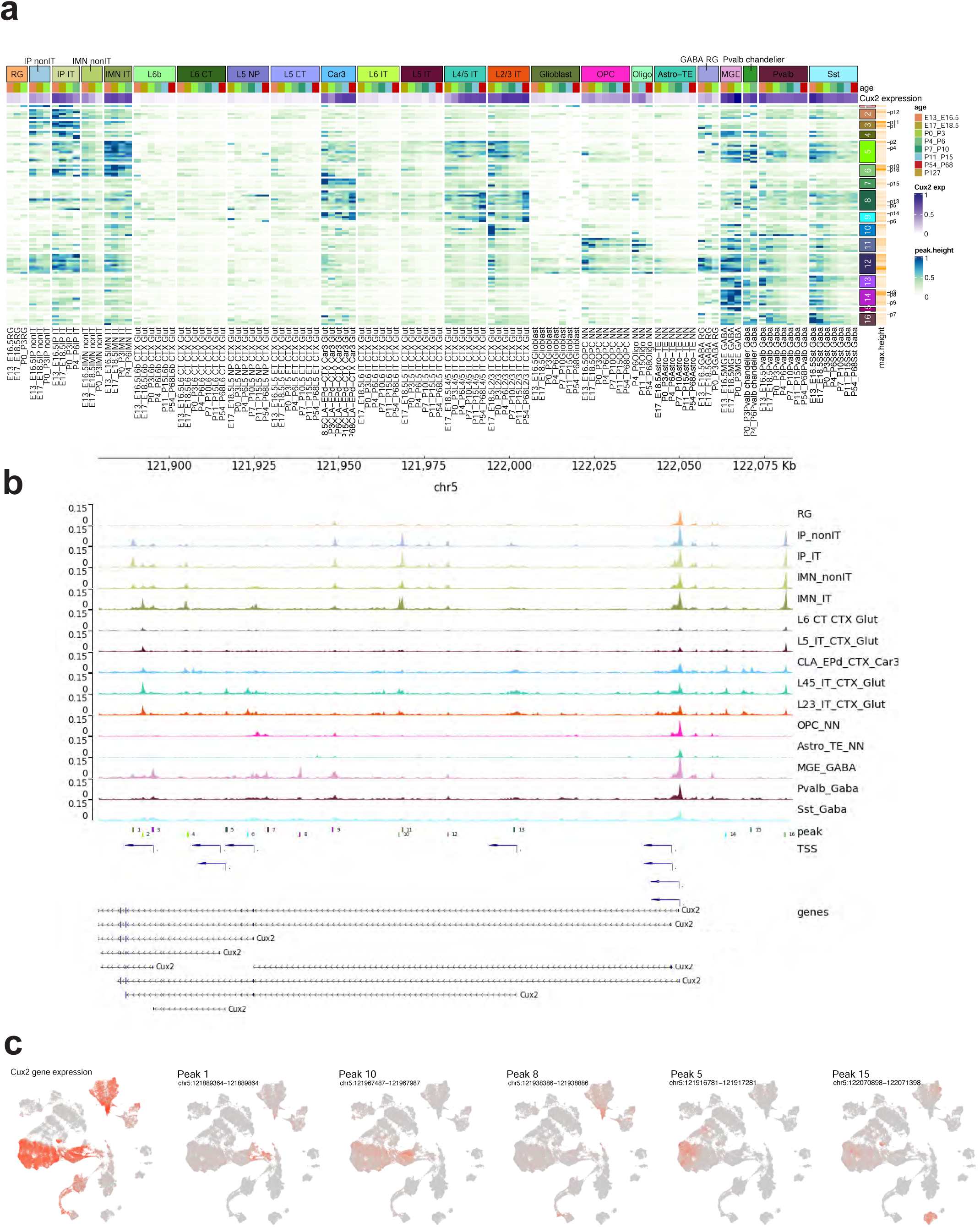
Differential accessibility peaks associated with the *Cux2* gene in different cell types or different developmental ages. **(a)** Heatmap representation of accessibility of differentially accessible peaks located in *Cux2* gene body and 50 Kb upstream. Each row corresponds to a peak, ordered by peak module, and each column corresponds to a cell category defined by subclass and age group. The *Cux2* gene expression level is shown in purple at the top. The heatmap color represents the average peak accessibility (height) in each subclass-by-age-group category, normalized with 1 indicating the maximum value for each peak and 0 indicating no accessibility. The peak module and maximum peak height are shown for each peak to the right. Specific peaks are numbered and labeled. **(b)** The accessibility tracks per subclass surrounding the *Cux2* gene, along with the genomic locations of labeled peaks in (a). TSS, transcription start site. **(c)** UMAP representation of Multiome cells, colored by *Cux2* expression and accessibility of a subset of peaks labeled in (a).

We also examined the epigenomic landscape of several other genes that are expressed in multiple cell types in different lineages. There seems to be a general trend for genes with long gene body to be regulated by distinct peaks at different developmental stages and in different cell types, as shown by another example, the ion channel receptor gene *Grik1* (**Extended Data Fig. 14).** *Grik1* is activated postnatally in L4/5 IT, L5 NP, OPC, MGE and CGE, and its 394 Kb gene body is associated with highly distinct peaks in each case. This mechanism allows optimization of regulatory pathway for each cell type independently with minimal interference from other cell types during evolution, providing a gene with greater flexibility to contribute to diverse cellular functions in various contexts. In contrast, transcription factor *Fezf2* has only 6 Kb gene body, and most of the regulatory elements are packed within a 10 Kb window around the gene body (**Extended Data Fig. 15)**. While we can still identify differential peaks between early-stage RG and IP cell types, L5 IT cell types and nonIT cell types, the distinctions are a lot more subtle. *Fezf2* is crucial for development of the central nervous system and is highly conserved across species due to strong evolutionary constraints, which may leave little room for evolving completely independent regulatory sites in each cell types.

### Epigenomic changes before and after eye opening

Motivated by the transcriptomic differences observed before and after eye opening (**Fig. 5**), we investigated the epigenomic differences during this developmental stage. For each subclass, we computed DA peaks between P7_P10 and P11_P15. A total of 32,865 DA peaks are identified, sorted by the subclass with the highest accessibility and the peak age group (**Fig. 8a**). More DA peaks are detected in IT subclasses, but significant differences are evident across all subclasses (**Fig. 8a**). Notably, among glutamatergic subclasses, more increasing peaks than decreasing peaks are seen after eye opening, particularly within L5 IT and L6 CT subclasses, with the least amount of difference in L5 NP subclass (**Fig. 8b**). Considerable overlap in DA peaks is found among glutamatergic subclasses, especially among the IT subclasses (**Fig. 8b**). Subsequently, we computed the correlation of epigenetic changes before and after eye opening across all subclasses (**Fig. 8c**). Strong correlations are evident between L2/3 IT, L4/5 IT and L5 IT subclasses. L6 IT and L6 CT exhibit weaker correlations with other IT subclasses, and L5 ET display even weaker correlations with IT and CT subclasses. L5 NP, non-neuronal and GABAergic subclasses show minimal correlated epigenetic changes with any other subclasses. The abundance of DA peaks within IT subclasses was partially attributed to their greater prevalence, providing enhanced statistical power for differential analysis. To mitigate this bias stemming from cell numbers, we also computed separately the sum of positive and negative changes in the common set of 32,865 DA peaks for each subclass (**Fig. 8d**). The metric solely assesses the absolute change without factoring statistical significance, rendering it less reliant on sample size. The overall amount of positive and negative changes shows the same trend as the number of DA peaks across different subclasses (**Fig. 8b**), albeit with smaller variation, e.g., the amount of change for L6 IT subclass, which has few DA peaks presumably due to smaller cell number, now is a lot more comparable with other IT and L6 CT subclasses (**Fig. 8d**).

**Figure 8.**
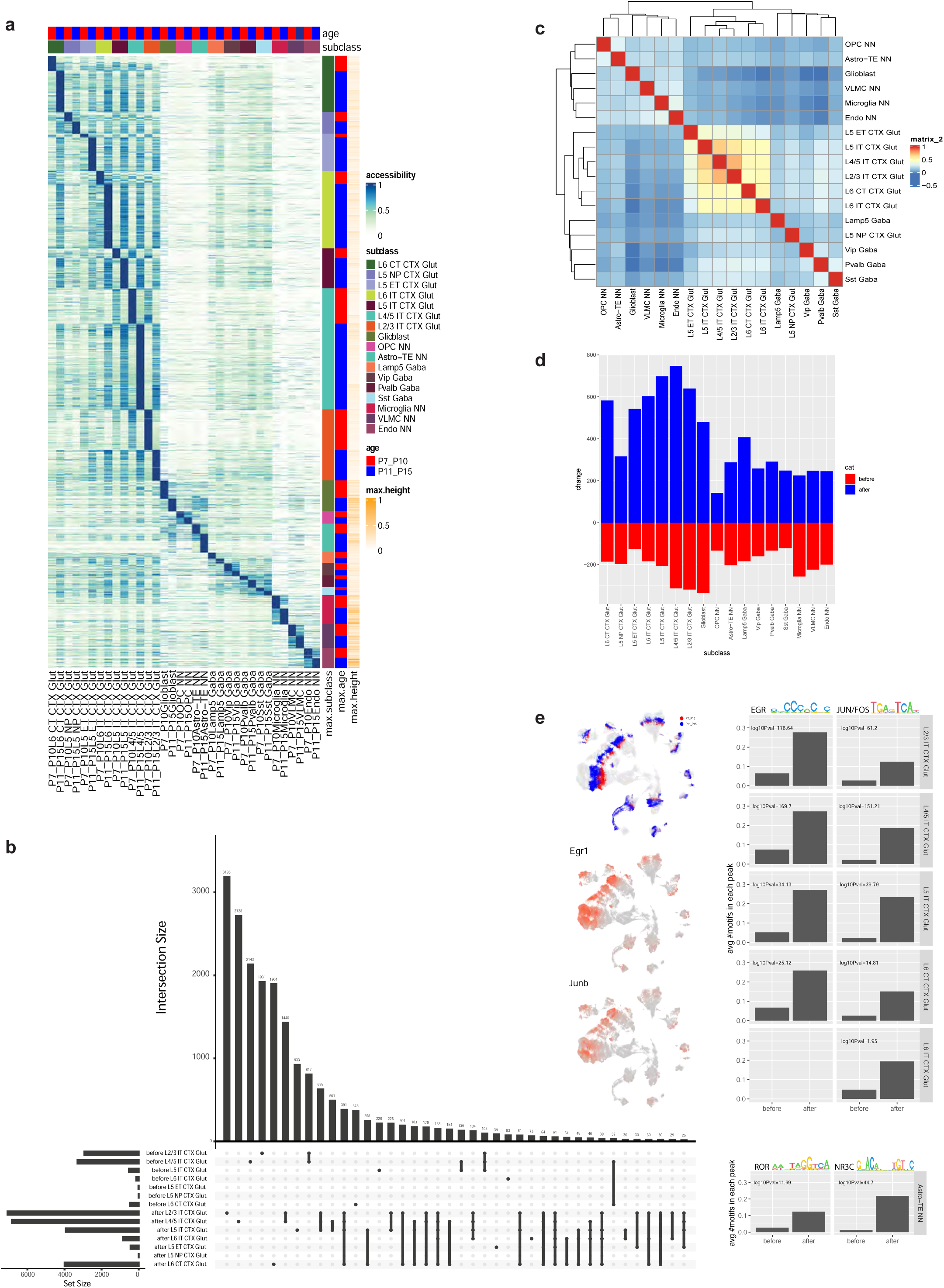
Cell-type specific chromatin accessibility changes before and after eye opening. **(a)** Heatmap representation of accessibility of DA peaks before and after eye opening. Each row corresponds to a peak, ordered by the subclass and age group with maximum accessibility. **(b)** Number of DA peaks before and after eye opening shared among different glutamatergic subclasses. Each column corresponds to a combination of different subclasses, and the bar height represents the number of peaks shared by the given combination of subclasses. The bar graph to the left of the subclass labels shows the total number of DA peaks for each subclass before or after eye opening. **(c)** Correlation of the chromatin accessibility changes before and after eye opening among all subclasses. The chromatin accessibility change is measured as the difference of average peak height between the two age groups for the given subclass, based on all the DA peaks defined in (a). **(d)** Cumulative positive and negative changes for each subclass before and after eye opening based on all the DA peaks defined in (a). **(e)** The differential motifs between increased and decreased DA peaks identified in each subclass. The average number of motif occurrences per peak is shown on the Y axis, the -log_10_(adjusted P value) for significance is labeled for each comparison. The expression values of putative transcription factor regulators are shown in the UMAP.

Finally, we tried to uncover the gene regulatory mechanisms driving the epigenomic changes associated with eye opening. We performed differential TF motif analysis between the increasing and decreasing peaks for the top six subclasses with the most DA peaks. Interestingly, we found enrichment of two neuronal activity-dependent motifs, EGR and JUN/FOS, in the increasing peaks across nearly every tested glutamatergic subclass, except for the lack of the EGR motif in L6 IT, likely due to insufficient statistical power (**Fig. 8e**). Correspondingly, there is a significant increase of *Egr1* and *Junb* gene expression after eye opening, consistent with our finding of increased expression of many immediate early genes (IEGs) after eye opening described above (**Fig. 5g-j**). This finding suggests that activity-dependent IEGs such as *Egr1* and *Junb*, induced following eye opening, can play profound roles in regulating extensive downstream epigenomic and transcriptomic changes that contribute to neuronal maturation. For astrocytes, we identified enrichment of nuclear receptor motifs, ROR and NR3C (**Fig. 8e**), while the ROR motif and *Rorb* and *Rora* genes were already identified as potential regulators of astrocyte differentiation in earlier analysis (**Fig. 6f**). *Rora*, *Rorb*, *Nr3c1* and *Nr3c2* all have increasing expression in astrocytes till adulthood and may contribute to astrocyte maturation (**Extended Data Fig. 7**). Recent study demonstrated that *Nr3c1* knockout in prefrontal cortex astrocytes disrupted memory recall^98^.

## Discussion

In this study, we created a comprehensive transcriptomic and epigenomic cell type atlas and trajectory map of the developing mouse visual cortex that densely covers the embryonic and postnatal developmental stages. We systematically identified the precise timing of the onset of all excitatory, inhibitory, and non-neuronal cell subclasses and types/clusters within the visual cortex, and we discover a pattern of continuous cell type diversification. We also systematically categorized large numbers of differentially expressed (DE) gene modules and differentially accessible (DA) chromatin peak modules that are concurrently associated with specific cell types and developmental ages, which serve as molecular signatures of cell type diversification and the emergence of new cellular and circuit properties (**Fig. 1a**).

Key new insights include the following. We find transcriptional heterogeneity within each of the embryonic cell populations, i.e., radial glia, glioblasts, intermediate progenitors, and immature neurons, suggesting early specification of cell fates that become increasingly pronounced and distinct with time (**Fig. 1-3**). Post-neurogenesis, we find that both excitatory and inhibitory neurons exhibit gradually increased complexity, with new subclasses and types emerging along the developmental timeline, including a burst of new cell types after eye opening and at critical period, especially for the IT and ET excitatory neurons and the Sst inhibitory neurons (**Fig. 2, 3, 5, Extended Data Fig. 5-9**). Throughout development, we find cooperative dynamic changes in gene expression and chromatin accessibility in specific cell types, identifying both chromatin peaks potentially regulating the expression of specific genes and transcription factors potentially regulating specific peaks (**Fig. 6, 8, Extended Data Fig. 12, 13**). In several prominent examples, we find that a single gene can be regulated by multiple peaks associated with different cell types and/or different developmental stages (**Fig. 7, Extended Data Fig. 14, 15**).

To determine the association of a distant peak with a target gene, most computational methods rely on correlating peak accessibility with promoter accessibility or target gene expression. Our findings suggest that while this approach may be effective in populations with limited cell type diversity, it may not apply to genes with complex regulatory landscapes. For such genes, where multiple peaks contribute to gene expression in specific cell types but not others, this assumption may fail. Our analysis, along with previous studies of whole-brain cell type atlases, reveals that many genes are often reused in different cell types with varying temporal dynamics, indicating that the impact may be more significant than previously understood, and we need more sophisticated analysis paradigm to address these cases.

In this study, we introduced a novel analysis paradigm that enables trajectory inference at a fine cell-type resolution by directly utilizing the temporal information embedded in the data (**Fig. 2, 3, Extended Data Fig. 5-9**). This method decouples the temporal gradient from the cell type gradients, resulting in trajectories that are easy to interpret and align with prior knowledge. The current method still has several limitations. Firstly, preserving rare transitional cell types in trajectory analysis remains challenging. Secondly, a more robust approach is needed to model the uncertainty in label transfer from adult to developmental stages. Additionally, managing cases where cell types outside the targeted areas appear in some but not all developmental stages is difficult, requiring intensive manual curation and is prone to errors. Lastly, we employed different methods for embryonic and postnatal trajectory analysis, which ideally should be unified. Nonetheless, this approach can be enhanced and applied to the whole brain in the future, allowing for the tracing of adult cell types back to early developmental stages and constructing developmental cell type atlases that can be directly linked to existing adult whole-brain atlases.

A widely accepted concept in early cortical development is a sequential inside-out model, namely, neural precursors (i.e., radial glia) generate deep-layer excitatory neurons first, then upper-layer excitatory neurons, and finally glial cells (astrocytes and oligodendrocytes). While our data is largely consistent with such a general timeline, with nonIT IP cells appearing the earliest at E13, we observe the emergence of both IT IP cells and glioblasts at E15.5, and the appearance of astrocytes and OPCs at E17 is also closely following that of the deep-layer IT neurons (**Fig. 2a,b**). Furthermore, within the nonIT and IT neuron classes, the emergence of subclasses is not entirely following the inside-out order. For the nonIT class, L6 CT, L5 ET and L6b (subplate) cells all appear at ∼E14.5, whereas the L5 NP cells emerge later, at E18.5. For the IT class, L6 IT and L5 IT emerge similarly at E17, whereas L4/5 IT and L2/3 IT cells emerge similarly at E18.5. Therefore, our transcriptomically defined trajectories suggest a more nuanced view, i.e., a staggered parallel differentiation process for the excitatory neuronal and glial cell types that are derived from the common pool of radial glia. Our observation of transcriptomic heterogeneity within the RG population, with nonIT and IT neuronal markers as well as glial markers appearing in a staggered overlapping manner further supports this view (**Fig. 2f**). Our findings are compatible with the body of recent studies that reveal extensive heterogeneity in the repertoire of cortical cell types each RG progenitor generates, which may be due to a series of probabilistic decisions in individual RG progenitors leading to varied lineage progression^30,55^.

We must note that our trajectories do not equal lineages, and it is not possible to derive lineage relationships from single-cell transcriptomic data only, as it provides a one-time snapshot of a cell without knowledge of the cell’s history. The transcriptomically defined trajectories depict the cell type or cell state transitions with time at population level. It is possible that some cells go through all the transitions while others skip certain steps. For example, an RG cell may generate nonIT cells first, then IT cells, and then turn its fate towards the generation of glial cells; whereas another RG cell may transition from neurogenic to gliogenic sooner and skip the generation of IT cells; and yet another RG cell may stay in the neurogenic state and never transition to gliogenic. These different scenarios, plus the asynchronous timing of the state transitions, could account for the heterogeneity observed in the RG population (**Fig. 2f**). A related issue is direct neurogenesis, i.e., an RG cell generates postmitotic nonIT or IT neurons directly, versus indirect neurogenesis, i.e., an RG cell generates an intermediate progenitor (IP) cell first and that IP cell then generates postmitotic nonIT or IT neurons^44,99^. Our transcriptomic data is unable to distinguish between the direct or indirect neurogenic history of any individual nonIT or IT neuron. However, our transcriptomic trajectories clearly define the IP cell types, which express the canonical IP marker gene *Eomes* and are separated into CR, nonIT and IT types, are a major intermediate step between RG and immature and mature neurons at the population level (**Fig. 2**). Therefore, despite the lack of detailed lineage information, the transcriptomic trajectory map provides a comprehensive overview of cell type and cell state transitions and the relationships among cell types across time that are inscribed in gene expression profiles.

Perhaps an even more remarkable finding is the extensive diversification of cell types after birth, with the total number of neuronal clusters increasing from 40 at P0, to 51 at P8, 68 at P16 and 95 at P25 (**Fig. 2g**). While nearly all cell subclasses are generated prenatally, vast majority of cell clusters emerge postnatally. This diversification coincides with the maturation of neurons, formation of synaptic connections, myelination, activity-dependent plasticity, etc. In particular, during the eye-opening stage (P11 to P14) and around the onset of critical period (P21), many new clusters emerge, especially for the IT excitatory neurons and the Sst and Vip inhibitory neurons (**Fig. 3, Extended Data Fig. 5-9**). Of the 19 IT neuron clusters, nearly half (9) emerge after P11, compared to 2 (both *Chrna6*+ L5 ET types) out of 15 nonIT neuron clusters. The 9 late-emerging IT clusters come from all subclasses – L2/3 IT, L4/5 IT, L5 IT and L6 IT. GABAergic subclasses also have substantial numbers of clusters emerging after P11, i.e., 13 of 20 for Sst, 4 of 9 for Pvalb, 11 of 17 for Vip, 2 of 3 for Sncg, and 6 of 9 for Lamp5. Using Patch-seq multimodal MET types, we were able to relate the developmental transcriptomic clusters defined here to the conventionally defined GABAergic neuron types based on axon-projection patterns^16,18,37^. We find that the Sst Martinotti and non-Martinotti cells with extensive axon-projecting diversity correspond to several specific and distinct groups of Sst clusters diverging from synaptogenesis and on (after P7), linking temporally precise transcriptional specificity with connectional specificity. *Cbln4*, a gene known to play critical roles in the synaptic targeting of excitatory neuron dendrites by SST interneurons^72^, has the highest expression in the Sst clusters that are L2/3-5 fanning Martinotti cells, L4-targeting Martinotti cells, or L2/3 fast-spiking-like cells.

We find that eye opening is also associated with broad-ranging, cell-type specific gene expression changes and activated chromatin accessibility peaks (**Fig. 5, 8**), extending beyond previous studies^64,81^. In these changes, the activation of immediate early genes and their transcriptional regulatory motifs in both excitatory and inhibitory neuron types are highly significant, presenting a mechanism for broad gene expression regulations to refine cell-type specific functions.

A remaining issue in defining cell types is the distinction between cell type and cell state. We recognize that this issue is particularly challenging to resolve during development, when a cell is constantly in transitioning states until it crosses a putative threshold to become a new type, given the extraordinary multidimensional transcriptional gradients across time, space and cell identity, as exemplified by the many gene modules we identified (**Fig. 4, Extended Data Fig. 10, 11**). Tentatively, a rough proximation is to assume that clusters may represent cell types while subclusters are more likely to reflect different states within a cell type. These can be refined as we gain better understanding of the developmental processes. Furthermore, given the highly complex and dynamic expression patterns of individual genes, it makes more sense to track different cell types and states via transcriptomic clusters and subclusters rather than individual marker genes. The transcriptomic and epigenomic developmental cell type atlas will help to delineate the relationship between clusters and marker genes in a precise way and facilitate the study of gene function in cell type and circuit development.

## Methods

### Mouse breeding and husbandry

All experimental procedures related to the use of mice were approved by the Institutional Animal Care and Use Committee of the Allen Institute for Brain Science, in accordance with NIH guidelines. Mice were housed in rooms with temperature (21–22 °C) and humidity (40–51%) control at no more than five adult animals of the same sex per cage. Mice were provided food and water ad libitum and were maintained on a regular 14:10 h light/dark cycle. Mice were maintained on the C57BL/6 J background. We excluded any mice with anophthalmia or microphthalmia.

The presence of vaginal plugs was monitored at 12-hour intervals (6 am and 6 pm). To harvest embryos with accuracy to 0.5 days, only dams with visible plugs were used to obtain embryonic time points. For postnatal time points, births were recorded at 12-hour intervals (6 am and 6 pm). Animal handling was reduced as much as possible until weaning at P21. At weaning, animals were separated from their mothers and opposite-sex siblings. Weaned mice were group-housed, kept separate from the opposite sex and maintained under normal housing conditions until dissection.

All donor animals used for data generation are listed in **Supplementary Table 1**. No statistical methods were used to predetermine sample size. In total we used 53 donors to collect scRNA-seq data from 919,547 cells across 35 timepoints between E11.5 and adulthood. We collected samples daily between E11.5 and P21, with the addition of E17.0 and E18.0 time points. After P21, we collected samples at P23, P25, P28, and adult samples between P54 and P68 (collectively simplified as P56). Brain dissections for all groups took place in the morning. From ages E11.5 and E12.5 we collected whole brain tissue, from ages E13.5 and E14.5 we collected cerebrum and brain stem (CH-BS), and from other ages we dissected visual cortex (VIS). For multiome data generation, we collected samples starting at E13.5 to adulthood. In total we collected multiome data from 378,541 nuclei from 37 donors across 15 time points. At embryonic time points we dissected CH-BS and at postnatal timepoints we collected either VIS or isocortex.

In some cases, transgenic mice were used for fluorescence-positive cell isolation by fluorescence-activated cell sorting (FACS). To enrich for neurons profiled by scRNA-seq, cells were isolated from the pan-neuronal Snap25-IRES2-cre line (RRID: IMSR_JAX:023525) crossed to the Ai14-tdTomato reporter (RRID: IMSR_JAX:007914) (31 out of 53 donors, **Supplementary Table 1**).

### Single-cell isolation

Single cells were isolated following a cell-isolation protocol developed at AIBS^100^. The brain was dissected, submerged in artificial cerebrospinal fluid (ACSF), embedded in 2% agarose, and sliced into 350-μm coronal sections on a compresstome (Precisionary Instruments). Block-face images were captured during slicing. ROIs were then microdissected from the slices and dissociated into single cells.

Dissected tissue pieces were digested with 30 U/ml papain (Worthington PAP2) in ACSF for 30 min at 30 °C. Due to the short incubation period in a dry oven, we set the oven temperature to 35 °C to compensate for the indirect heat exchange, with a target solution temperature of 30 °C. Enzymatic digestion was quenched by exchanging the papain solution three times with quenching buffer (ACSF with 1% FBS and 0.2% BSA). Samples were incubated on ice for 5 min before trituration. The tissue pieces in the quenching buffer were triturated through a fire-polished pipette with 600-µm diameter opening approximately 20 times. The tissue pieces were allowed to settle and the supernatant, which now contained suspended single cells, was transferred to a new tube. Fresh quenching buffer was added to the settled tissue pieces, and trituration and supernatant transfer were repeated using 300-µm and 150-µm fire-polished pipettes. The single-cell suspension was passed through a 70-µm filter into a 15-ml conical tube with 500 µl of high-BSA buffer (ACSF with 1% FBS and 1% BSA) at the bottom to help cushion the cells during centrifugation at 100g in a swinging-bucket centrifuge for 10 min. The supernatant was discarded, and the cell pellet was resuspended in the quenching buffer. We collected 483,755 cells without performing FACS. The concentration of the resuspended cells was quantified, and cells were immediately loaded onto the 10x Genomics Chromium controller.

To enrich for neurons or live cells in some samples, cells were collected by FACS (BD Aria II running FACSdiva v8) using a 130-μm nozzle, following a FACS protocol developed at AIBS^101^. Cells were prepared for sorting by passing the suspension through a 70-µm filter and adding Hoechst or DAPI (to a final concentration of 2 ng/ml). The sorting strategy with example images has been described previously^101^. We collected 17,865 Calcein- and Hoechst-positive cells, 17,974 Hoechst-positive cells, 13,912 RFP-positive, and 283,089 RFP- and Hoechst-positive cells (**Extended Data Fig. 1d, Supplementary Table 1**). Around 30,000 cells were sorted within 10 min into a tube containing 500 µl of quenching buffer. Each aliquot of sorted 30,000 cells was gently layered on top of 200 µl of high-BSA buffer and immediately centrifuged at 230g for 10 min in a centrifuge with a swinging-bucket rotor (the high-BSA buffer at the bottom of the tube slows down the cells as they reach the bottom, minimizing cell death). No pellet could be seen with this small number of cells, so we removed the supernatant and left behind 35 µl of buffer, in which we resuspended the cells. Immediate centrifugation and resuspension allowed the cells to be temporarily stored in a high-BSA buffer with minimal ACSF dilution. The resuspended cells were stored at 4 °C until all samples were collected, usually within 30 min. Samples from the same ROI were pooled, cell concentration quantified, and immediately loaded onto the 10x Genomics Chromium controller.

### Single-nucleus isolation

Mice were anaesthetized with 2.5–3% isoflurane and transcardially perfused with cold, pH 7.4 HEPES buffer containing 110 mM NaCl, 10 mM HEPES, 25 mM glucose, 75 mM sucrose, 7.5 mM MgCl2, and 2.5 mM KCl to remove blood from brain^102^. Following perfusion, the brain was dissected quickly, frozen for 2 min in liquid nitrogen vapor and then moved to −80 °C for long term storage following a freezing protocol developed at AIBS^103^.

For VIS dissections, frozen mouse brains were sectioned using a cryostat with the cryochamber temperature set at −20 °C and the object temperature set at −22 °C. Brains were securely mounted by the cerebellum or by the olfactory region onto cryostat chucks using OCT (Sakura FineTek 4583). Tissue was trimmed at a thickness of 20–50 µm and once at the desired location slices with thickness of 300 µm were generated to dissect out ROI(s) following reference atlas. Images were taken while leaving the dissection in the cutout section. Nuclei were isolated using the RAISINs^104^ method with a few modifications as described in a nuclei isolation protocol developed at AIBS^105^. In short, excised tissue dissectates were transferred to a 12-well plate containing CST extraction buffer. Mechanical dissociation was performed by chopping the dissectate using spring scissors in ice-cold CST buffer for 10 min. The entire volume of the well was then transferred to a 50-ml conical tube while passing through a 100-µm filter and the walls of the tube were washed using ST buffer. Next the suspension was gently transferred to a 15-ml conical tube and centrifuged in a swinging-bucket centrifuge for 5 min at 500 rcf and 4 °C. Following centrifugation, the majority of supernatant was discarded, pellets were resuspended in 100 µl 0.1× lysis buffer and incubated for 2 min on ice. Following addition of 1 ml wash buffer, samples were gently filtered using a 20-µm filter and centrifuged as before. After centrifugation most of the supernatant was discarded, pellets were resuspended in 10 µl chilled nuclei buffer and nuclei were counted to determine the concentration. Nuclei were diluted to a concentration targeting 5,000 nuclei per µl.

### cDNA amplification and library construction

For 10x scRNA-seq, the cell suspensions were processed using the Chromium Single Cell 3′ Reagent Kit v3 (1000075, 10x Genomics)^106^. We followed the manufacturer’s instructions for cell capture, barcoding, reverse transcription, cDNA amplification and library construction. We loaded 8,876 ± 2,980 cells per port. We targeted a sequencing depth of 120,000 reads per cell; the actual average achieved was 64,723 ± 60,061 reads per cell across 92 libraries (**Supplementary Table 1**).

For 10x Multiome processing, we used the Chromium Next GEM Single Cell Multiome ATAC + Gene Expression Reagent Bundle (1000283, 10x Genomics). We followed the manufacturer’s instructions for transposition, nucleus capture, barcoding, reverse transcription, cDNA amplification and library construction^107^. For the snMultiome libraries, we loaded 10,108 ± 4,334 nuclei per port. For snRNA-seq we targeted a sequencing depth of 120,000 reads per nucleus. The actual average achieved, for the nuclei included in this study, was 105,701 ± 52,241 reads per nucleus across 41 libraries (**Supplementary Table 1**). For snATAC-seq we targeted a sequencing depth of 85,000 reads per nucleus. The actual average achieved, for the nuclei included in this study, was 124,023 ± 67,263 reads per nucleus across 41 libraries.

### Sequencing data processing and QC

To remove low-quality cells, we developed a stringent QC process. Cells were first classified into broad cell classes after mapping to our established Allen Brain Cell Atlas for the whole mouse brain (ABC-WMB Atlas)^15^, and cell quality was assessed based on gene detection, QC score, and doublet score. The QC score was calculated by summing the log-transformed expression of a set of genes whose expression level is decreased significantly in poor quality cells. Doublets were identified using a modified version of the DoubletFinder algorithm (available in scrattch.hicat, https://github.com/AllenInstitute/scrattch.hicat, v1.0.9) and removed when doublet score was > 0.3. In prenatal time points, neuronal precursors of non-cortical origin were excluded by low expression of Foxg1, Emx1 or Emx2. Using different QC scores and gene-count thresholds among different cell classes (**Supplementary Table 2**), we filtered out 158,230 cells and kept 761,419 cells for 10xv3 scRNA-seq data (**Extended Data Fig. 1a,b**).

We adopted a similar strategy to filter low-quality nuclei for the 10xMulti snRNA-seq dataset. Nuclei were first classified into broad cell classes after mapping to the existing ABC-WMB Atlas, and cell quality was assessed based on gene detection, QC score, and doublet score. For 10xMulti snRNA-seq dataset, although the overall gene counts were lower compared to 10xv3 scRNA-seq dataset, they showed stronger bimodal distribution of QC metrics, so we could afford to keep the high cutoffs. The different QC scores and gene-count thresholds among different cell classes are shown in **Supplementary Table 2**. For 10xMulti snATAC-seq data, we used the default criteria implemented in ArchR^86^: number of unique nuclear fragments (nFrags > 1000) and signal-to-background ratio (TSS > 3). For 10xMulti dataset, only nuclei having passed both snRNA-seq and snATAC-seq QC criteria (total 304,645 nuclei) were included in the downstream analysis (**Extended Data Fig. 1a,c**).

### Inferring synchronized developmental age

To estimate the synchronized developmental age for each single cell, we trained K-Nearest Neighbors (KNN) models (**Extended Data Fig. 2a**). We first performed global *de novo* clustering for 10xv3 single cell datasets across all time points using R package scrattch.bigcat^15^ (https://github.com/AllenInstitute/scrattch.bigcat). The automatic iterative clustering method, iter_clust_big, was used with stringent differential gene expression criteria as previous study^13^: q1.th = 0.5, q.diff.th = 0.7, de.score.th = 150, min.cells = 50. We then performed principal component analysis (PCA) based on the gene expression matrix of 5,824 marker genes derived from this *de novo* clustering. We down sampled up to 200 cells per cluster so that PCA could proceed without computing memory issues. The principal components (PC) based on sampled cells were then projected to the whole datasets. We selected the top 100 PCs and removed one PC with more than 0.7 correlation with the technical bias vector, defined as log_2_(gene count) for each cell. The KNN algorithm identified 10 nearest neighbors to each of the single cells in the input data based on their distances computed using the selected 99 PCs. The inference of synchronized developmental age using KNN algorithm was run iteratively: in the first iteration, each cell was assigned a predicted age based on the most common age among its 10 neighbors. In the following iterations, the predicted age of each cell was assigned based on the most common predicted age from the previous iteration of its 10 neighbors. Ten iterations were run until convergence into the final synchronized ages (**Fig. 1h, Extended Data Fig. 2a**).

### Label transfer and clustering

Label transfer and clustering was conducted in synchronized age (**Extended Data Fig. 2a**). For all adult cells (P56), we assigned cell type identities by mapping them to ABC-WMB Atlas^15^ using R package scrattch.mapping (v0.55, https://github.com/AllenInstitute/scrattch.mapping)^108^. After mapping, we conducted DEG analysis between the transferred cluster labels and merged across them using the DE genes to get the final cell type identities at cluster level. For cells from younger age bins from P0 to P28, we assigned cell type identities using reciprocal PCA (RPCA) implemented in Seurat (**Extended Data Fig. 2a, 3**). For example, cells from P20_P28 were mapped to P56, cells from P17_P19 were mapped to P20_P28, etc. If a cluster has fewer than 10 cells within a specific age bin, the cells are reassigned to the nearest cluster based on the 10 nearest neighbors within the same age bin. After assigning the cell types, iterative clustering was performed for each synchronized age bin (**Extended Data Fig. 2d**) within each cluster to identify subclusters at each synchronized age bin.

For global clusters which are dominantly from embryonic stage (E11.5 to E18.5), we used scrattch.mapping to assign cell types based on La Manno et al^33^ mouse development study covering E9 to E18, using a gene list of 2947 markers, derived from the study’s cluster-specific marker genes. Global clusters which are mapped to Radial glia are assigned NEC subclass (dominant by cells from E11.5 and E12.5, expressing *Hmga2*) and RG (dominant by cells from E13.5-E16.5). Global clusters mapped to Neuroblast are identified as the IP class (characterized by *Eomes* expression). Neurons born early at E11.5 and E12.5, characterized by enrichment in *Reln*, *Trp73* and *Calb2*, are classified as CR Glut subclass. According to trajectory analysis, IP clusters at E11.5 and E12.5 that are enriched in *Crabp2* and *Ebf2*, and which give rise to CR Glut, are categorized as IP CR subclass. Similarly, informed by the trajectory analysis, IP clusters enriched in *Lhx9*, *Rmst*, *Nhlh1*, and *Nhlh2* are classified as the IP nonIT subclass, whereas those with higher levels of *Pou3f2* are classified as the IP IT subclass. Embryonic global clusters that are highly enriched in *Ncam1*, *Dcx* and *Neurod6*, with low expression of *Eomes,* are annotated as the IMN class. Within the IMN class, clusters enriched in *Fezf2* are labeled as the IMN nonIT subclass, while those enriched in *Pou3f2* are labeled as the IMN IT subclass. Cells within each embryonic subclass are merged into one cluster, followed by iterative clustering within each cluster and each synchronized age bin to identify subclusters. Finally, we merged the subclusters within each cluster that do not pass the DEG criteria: q1.th = 0.4, q.diff.th = 0.7, de.score.th = 150, min.cells = 10.

The final developmental cell-type taxonomy with annotations at class, subclass, cluster and subcluster levels is summarized in **Supplementary Table 3**. All DE genes are shown in **Supplementary Table 4**.

### Reconstruction of the developmental trajectory

In postnatal stage, to connect each cluster observed in a synchronized age bin with its most probable antecedent cluster from the previous synchronized age bin, we used the mutual nearest neighbors (MNN) approach (**Extended Data Fig. 2b**). First, we merged all cells from the two consecutive synchronized age bins using Seurat. Integration using reciprocal PCA (RPCA) and batch correction were performed among libraries from these two age bins. After integration, we performed PCA, from which we calculated Euclidean distances between individual cells from the earlier and later age bins. We then determined edge weights between clusters of the successive age bins using a bootstrapping strategy. For cells of each cluster in the later age bin, we identified their 50 closest neighbor cells from the earlier age bin and then calculated the proportion of these neighbors derived from each potential antecedent cluster. We repeated these steps 100 times with 90% subsampling from the same embedding. We then took the median proportions as the set of weights for edges between a cluster and its potential antecedents. Edge weights > 0.2 from the PCA embedding were retained and shown in supplementary table, and we chose the edge with max weight for the resulting trajectory (**Supplementary Table 5**).

In embryonic stage, cells are changing dramatically within the same age. We adopted the above strategy in pseudo-time that was computed by Monocle3^42^ (**Extended Data Fig. 2c**). For cells in each cluster, we identified their 50 closest neighbor cells from clusters that have earlier median pseudo-time than itself in a bootstrapping strategy. Same as the postnatal stage, edge weights < 0.2 were removed. The developmental trajectory across the entire timeline from E11.5 to P56 is summarized in **Supplementary Table 5** (**Fig. 2a**, **3**).

### Pseudo-time

We computed the overall pseudo-time (**Fig. 1i**) based on the entire developmental trajectory generated in the above section. The pseudo-time was computed separately for the three independent trajectories: the trajectory of all excitatory neurons and glia that are derived from NEC, that of MGE, and that of CGE. One random cell in the starting cluster of each trajectory was set to be at pseudo-time = 0. Along each trajectory, we computed pseudo-time of each cell in each cluster-by-age-bin cell group by comparing its distance with the median of its most probable antecedent cluster-by-age-bin cell group in the global PCA embedding generated in the “Inferring synchronized developmental age” section.

### Identification of gene modules

Synchronized age associated co-regulated genes specific to each subclass or class were determined using an unsupervised clustering approach. First, we computed pairwise DE genes between subclasses or classes using scrattch.bigcat at each synchronized age bin. Then within each subclass or class, we detected DE genes that were significant across synchronized age bins. After that, we computed the average expression of each DE genes among cells in each subcluster. Finally, we performed Louvain clustering (k = 5, resolution = 2) on the average expression of each gene within the subclass or class to identify gene co-expression modules. All the gene modules are summarized in **Supplementary Table 6**.

### Gene ontology enrichment analysis

We were interested in relating various gene modules to known biological processes. For this task we performed gene set enrichment analyses using the R package clusterProfiler 4.0^109^ and gprofiler^110^. The function gconvert from gprofiler2 was used to convert gene IDs to their Ensmbl IDs. The functions enrichGO and simplify from clusterProfiler were then used to enrich gene ontology terms from all three GO databases (molecular function, biological process, and cellular component). An adjusted p-value cutoff of 0.01 was used to determine significant GO terms.

### Integration of Multiome and scRNA-seq datasets and label transfer

For assigning identities of nuclei from the Multiome modality, we mapped the Multiome snRNA-seq transcriptomes to the scRNA-seq based developmental cell-type taxonomy described above. Briefly, we first performed global *de novo* clustering of the snRNA-seq data and derived DE genes among all clusters. We then integrated scRNA-seq data (subsampled up to 200 cells/cluster) and Multiome snRNA-seq data (all nuclei) via scVI^85^ using DE genes from the scRNA-seq developmental cell-type taxonomy and from global clustering of the Multiome snRNA-seq data (**Supplementary Table 4**). In the integrated latent space, we applied a Random Forest classifier to predict each nucleus’ most probable cell type identity, using the scRNA-seq developmental cell-type taxonomy as reference. Clusters with a mean mapping probability of less than 0.2 were excluded. Lastly, we performed further annotation and QC of each predicted cluster and filtered out a small set of clusters deemed to be low quality or outside of cortex. The final Multiome snRNA-seq to scRNA-seq developmental cell-type taxonomy mapping result is shown in **Supplementary Table 8** (**Fig. 1b**, **6a-c**).

### Multiome peak calling

To call chromatin accessibility peaks in the snATAC-seq data, we first categorized Multiome cells (nuclei) according to both subclass and age group. To accumulate enough samples with sufficient statistical power for comparative analysis, we combined consecutive ages into the following groups: E13_E16.5, E17_E18.5, P0_P3, P4_P6, P7_P10, P11_P15, and P54_P68. We kept only the subclass-by-age-group category with more than 50 cells. We generated pseudo-bulk replicates using the ArchR^86^ function addGroupCoverages. We created a reproducible merged peak set using function addReproduciblePeakSet. Finally, we built the peak by cell matrix, which contains insertion counts within the merged peak set using function “addPeakMatrix”.

### Identification of differentially accessible peaks

To identify differentially accessible (DA) peaks, as the peak presence in each cell is mostly binary, we chose the chi-squared test to evaluate the statistical significance of DA peaks across all 958,146 peaks identified above between every pair of subclass-by-age-group category. In addition to the log foldchange (log_2_FC) and adjusted P-value (adj.P) based on the chi-square test, we also computed the fraction of cells in each category with non-zero counts for each peak. To choose statistically significant DA peaks, we required log_2_FC > 1, adj.P < 0.05 and fraction of cells with non-zero value in the foreground category to be > 0.05. This method was implemented in “de_all_pairs” function in the scrattch.bigcat package with extensive parallelization for efficiency. Because of the extensive diversity in cell types overall, we opted for conducting pairwise comparisons instead of one-versus-all comparisons. This decision was made because the cell types in the background group exhibit high heterogeneity in one-versus-all comparison scenarios, which poses challenges in detecting subtle differences. The all-pairwise approach offers enhanced accuracy in identifying DA peaks across both similar and dissimilar pairs of cell categories.

### Identification of peak modules with similar cell type and temporal specificity

To identify peaks regulating different cell types at different developmental stages, we first extracted the DA peaks for each age group across different subclasses. We then pooled all the DA peaks identified between different subclasses across all age groups and clustered them to identify peak modules. To do that, we first computed the peak-by-category matrix as the average number of reads in each peak per subclass-by-age-group category, divided by the total number of reads across all peaks per subclass-by-age-group category, then multiplied by 30,000. The clustering was performed on peak-by-category matrix, subset to the DA peaks, using Jaccard-Leiden clustering algorithm. We first computed for each peak the K-nearest neighbors (k = 10) using “Cosine” similarity metrics, then computed the Jaccard similarity graph based on the number of shared nearest neighbors between every pair of peaks, and finally performed Leiden clustering algorithm based on the Jaccard graph. In most cases, we used resolution index = 2. In cases we observed more heterogeneity within the peak module, we increased the resolution index accordingly. This method is robust, efficient, and scalable, and generates peak modules with great cell type and temporal specificity.

### Differential motif analysis

We first scanned all the peak sequences using motif database using ArchR “addMotifAnnotation” function, which produced a matrix including the number of motif occurrences in each peak. To reduce the amount of redundancy of highly similar motifs, we used motifSet = “vierstra”, collection = “archetype”. This motif database includes “motif archetypes”, which represent clustered motifs that have been essentially deduplicated based on similarity^111^. To perform differential motif analysis on peaks in different modules, we again used chi-square test between all pairs of modules using “de_all_pairs” function, using cutoff log_2_FC > 2, adj.P < 0.05, and fraction of peaks with non-zero motif occurrences in the foreground greater than 0.1. Once more, we conducted pairwise comparison across all peak modules, as we did not have sufficient prior knowledge of which peak modules might share common or distinct motifs. This strategy allowed us to identify enriched motifs in different combination of peak modules. We simplified the motif name by using the corresponding transcription factor (TF) family name and tried to identify members of the TF family with similar gene expression cell type specificity as potential regulators of the peak modules.

### Identification of peak/gene pairs with matching accessibility and gene expression

We first extracted all the peak and gene pairs such that the gene is located within the 5 Mb window centered at the peak. Then we computed the correlation between the average peak accessibility and average gene expression based on the Multiome dataset across subclass-by-age-group categories. Given that a gene can be regulated by different peaks in different cell types and/or different developmental stages, the correlation is computed only within different subsets in different contexts, e.g. within IT subclasses only. We chose a minimal correlation of 0.5 to select such peak/gene pairs. Furthermore, we computed the average accessibility profile across subclass-by-age-group categories for all the peaks within each peak module. Subsequently, we calculated the correlation between the average expression within each subclass-by-age-group category of each gene and the peak module average profiles described above. We then filtered and retained only those peak/gene pairs if the gene has the strongest correlation with the peak module corresponding to the respective peak. To accommodate space constraints, for each peak module, only the top 500 selected peak/gene pairs with the strongest peak/gene correlations were included for visualization (**Supplementary Table 9**).

## ACKNOWLEDGEMENTS

We are grateful to the Transgenic Colony Management, Lab Animal Services, Molecular Biology, Histology, and Data and Technology teams at the Allen Institute for technical support. The research was funded by the U19MH114830 grant from National Institute of Mental Health to H.Z., under the BRAIN Initiative of National Institutes of Health (NIH). The content is solely the responsibility of the authors and does not necessarily represent the official views of NIH and its subsidiary institutes. This work was also supported by the Allen Institute for Brain Science. The authors thank the Allen Institute founder, Paul G. Allen, for his vision, encouragement, and support.

## Author Contributions

Conceptualization: H.Z. Data analysis lead and coordination: C.T.J.vV. Data generation (scRNA-seq and Multiome): C.T.J.vV., E.D.T., D.B., D.C., T.C., M.C., M.J.D., R.F., J. Gloe, N.G., J. Guzman, C.R.H., D.H., W.H., K.J., R.M., E.M., N.P., T.P., N.V.S., J.S., A. Torkelson, A. Tran, H.T., K.R., B.L., N.D., K.A.S., Z.Y., H.Z. Data processing and analysis: Y.G., C.T.J.vV., C.L., A.B.C., R.C., J. Goldy, B.N., J.W., M.J.H., K.A.S., Z.Y., H.Z. Project management: C.P., K.A.S. Management and supervision: C.T.J.vV., K.R., B.L., M.J.H., N.D., K.A.S., B.T., Z.Y., H.Z. Manuscript writing and figure generation: Y.G., C.T.J.vV., Z.Y., H.Z. Manuscript review and editing: Y.G., C.T.J.vV., Z.Y., H.Z.

## Competing Interests

H.Z. is on the scientific advisory board of MapLight Therapeutics, Inc. The other authors declare no competing interests.

## Additional Information

Correspondence and requests for materials should be addressed to: H.Z. or Z.Y.

## Data Availability

Primary data are being made available through BRAIN Initiative Cell Atlas Network (BICAN), www.portal.brain-bican.org, and Neuroscience Multi-omic Data Archive (NeMO), https://nemoarchive.org/.

## Code Availability

Data analysis code used in the manuscript is available via github https://github.com/AllenInstitute/scrattch.bigcat and https://github.com/AllenInstitute/MouseDev/tree/main/DevVIS.

## Figure Legends and Extended Data Figure Legends

(see below with figures and extended data figures)

## Supplementary Tables

**Supplementary Table 1:** scRNA-seq and Multiome libraries, with metadata.

**Supplementary Table 2:** QC criteria by class for both scRNA-seq and Multiome data.

**Supplementary Table 3:** Transcriptomic cell type taxonomy and atlas of the developing mouse visual cortex, including subclass composition at each age. The adult clusters are also mapped to Tasic et al 2018 (ref 11) cortical cell type taxonomy.

**Supplementary Table 4:** DE genes for the transcriptomic cell type atlas of the developing mouse visual cortex, and DE genes from global clustering of the scRNA-seq or Multiome snRNA-seq data.

**Supplementary Table 5:** Cell type trajectory trees of the developing mouse visual cortex.

**Supplementary Table 6:** Gene co-expression modules at class and subclass levels.

**Supplementary Table 7:** DE genes before and after eye opening.

**Supplementary Table 8:** Multiome snRNA-seq developmental taxonomy.

**Supplementary Table 9:** Differential chromatin accessibility peak modules, with peak/gene pairs (top 500 per peak module).

---

**Extended Data Figure 1.**
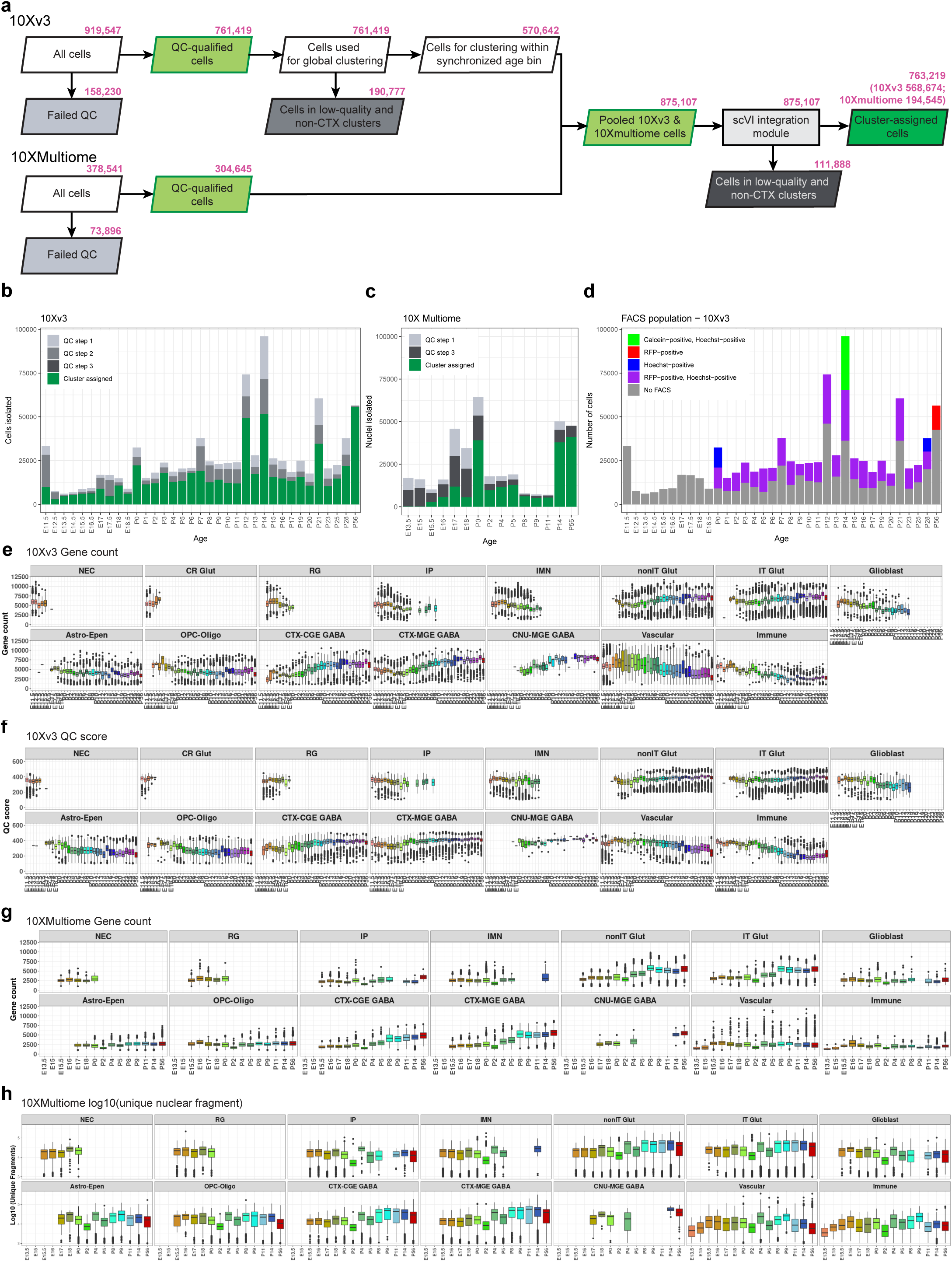
scRNA-seq and Multiome data processing and analysis workflow and quality control. **(a)** Number of cells at each step in the scRNA-seq and Multiome data processing and analysis pipeline. The identification of doublets and low-quality cells and clusters is described in detail in Methods. The 10xv3 and 10x Multiome data were first QC-ed and analyzed separately. After initial clustering the datasets were combined and QC-ed again before and after joint clustering. **(b-c)** Number of cells after each QC step in scRNA-seq (b) and Multiome data (c). The color codes of QC steps correspond to the colored QC boxes in (a). **(d)** Number of cells from each FACS population in scRNA-seq data. **(e-h)** Box plots of gene detection (e) and QC score (f) for 10xv3, and gene detection (g) and number of unique fragments (h) for 10x Multiome, per cell across different cell classes and ages.

**Extended Data Figure 2.**
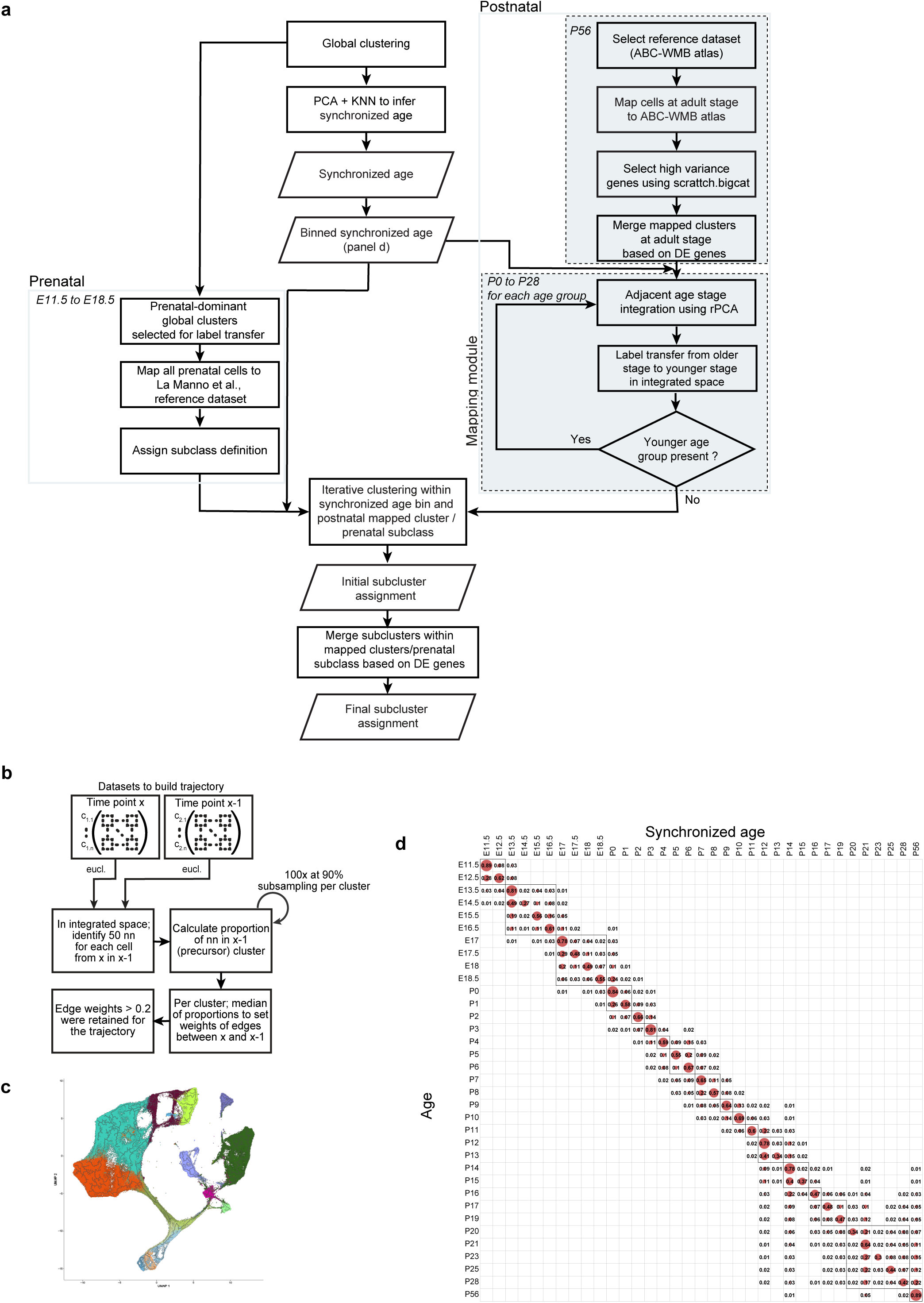
Detailed scRNA-seq and Multiome data analysis workflow. **(a)** Adjacent cell type mapping and clustering pipeline. **(b)** Mutual nearest neighbor (MNN) algorithm implementation for building trajectories. **(c)** Trajectory of glutamatergic cells built from Monocle3, showing that the embryonic part of the trajectory looks reasonable, but the postnatal part of the trajectory appears erratic. **(d)** Confusion matrix of the fraction of shared cells between each actual age and synchronized age. Boxes denote synchronized age bins.

**Extended Data Figure 3.**
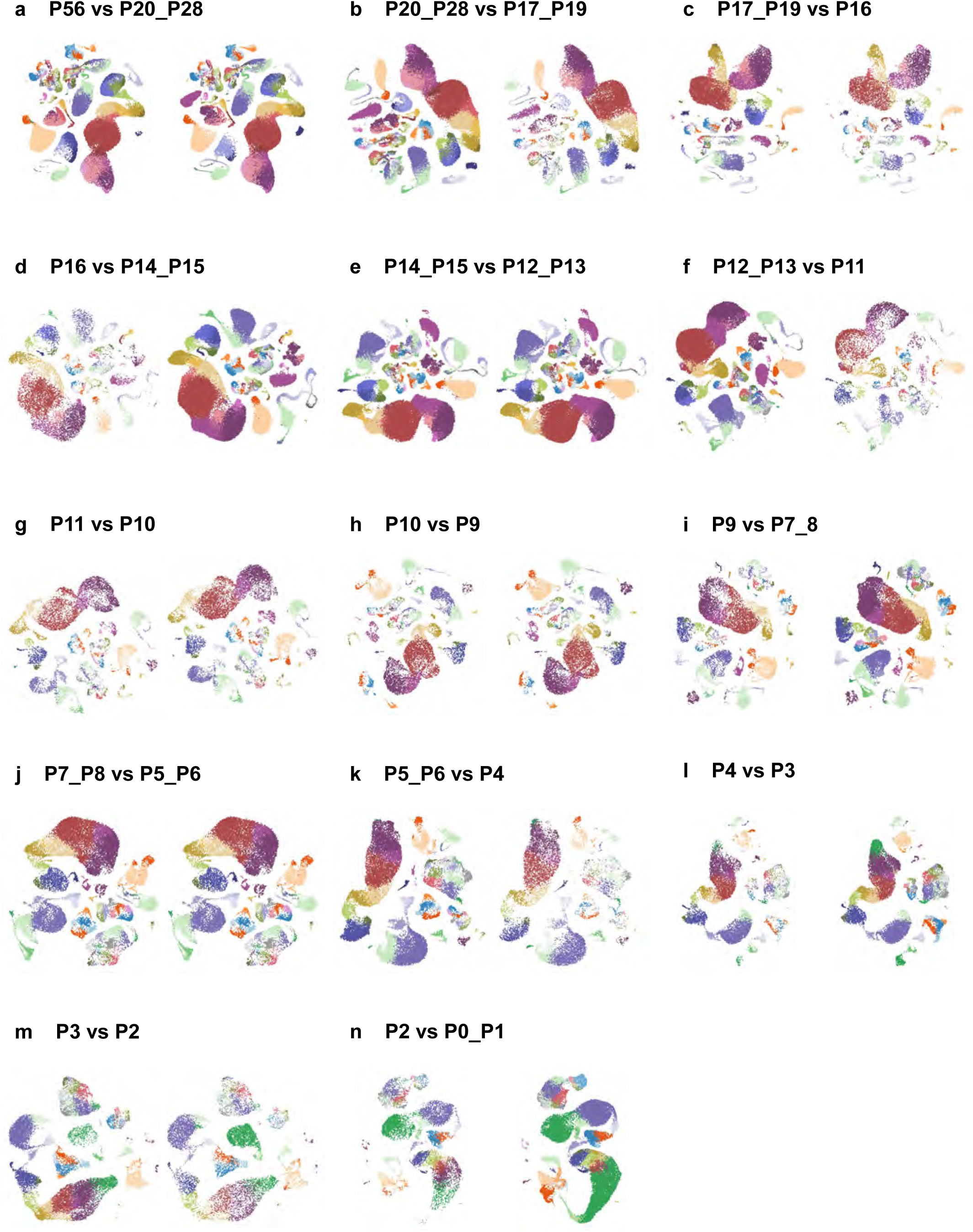
Integration between adjacent age bins for label transfer. (a-n) UMAP comparison of each synchronized age bin with its adjacent younger age bin after integration and label transfer, showing common clusters.

**Extended Data Figure 4.**
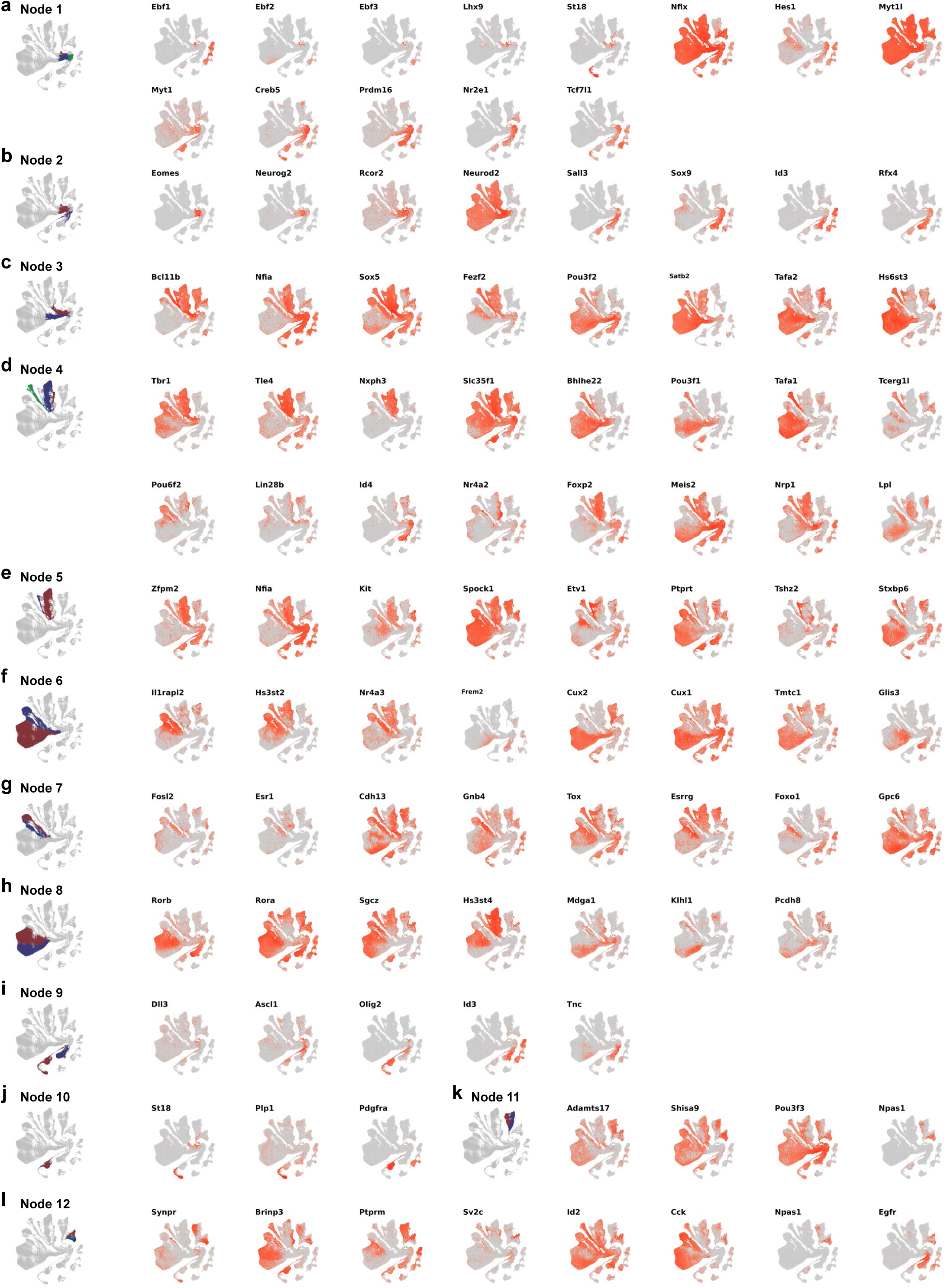
Expression of branching marker genes on UMAP. (a-l) Expression of marker genes at each branching node corresponding to Figure 2a.

**Extended Data Figure 5.**
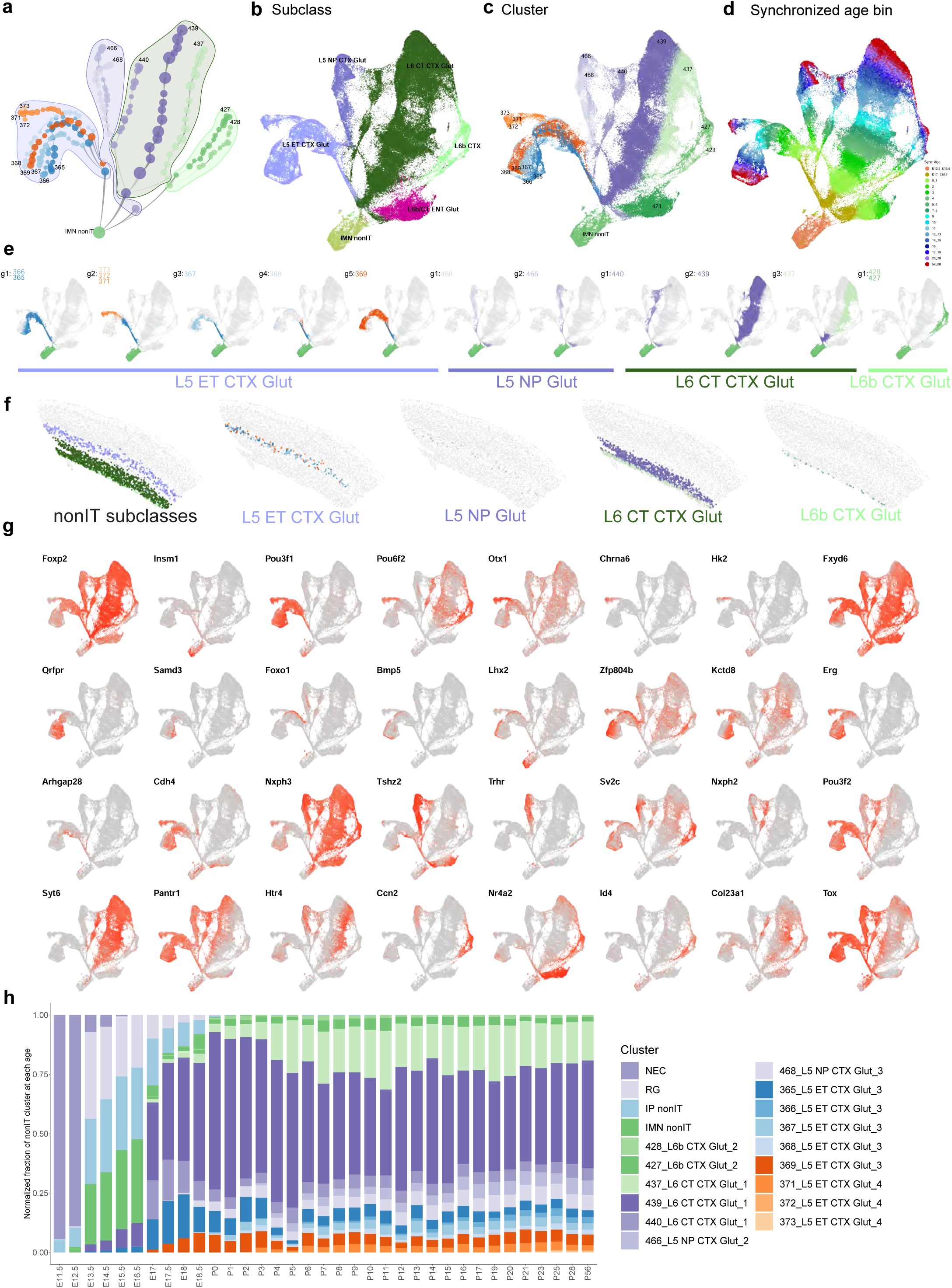
Developmental trajectories of visual cortex nonIT Glut cell types. **(a)** Transcriptomic trajectory tree for nonIT clusters starting from the common IMN nonIT antecedent. Nodes are clusters subdivided by synchronized age bins, and edges represent antecedent-descendent relationship between adjacent nodes, with thinner end at the antecedent node, and thicker end at the descendent node. Nodes are grouped by subclass, and adult clusters are labeled. Nodes from L6b/CT ENT subclass are not included. **(b-d)** UMAP for nonIT cells colored by subclass (b), cluster (c) and synchronized age bin (d). **(e)** Clusters are grouped together based on similar trajectories. Within each cluster group, all cells along their trajectories, including all antecedent nodes, are shown and are colored by cluster membership. **(f)** Spatial distribution of nonIT subclasses and clusters within each subclass at adult stage, based on the ABC-WMB Atlas^15^. **(g)** Marker genes illustrating cell type diversification along trajectories. **(h)** Cluster composition of all nonIT cells at each age.

**Extended Data Figure 6.**
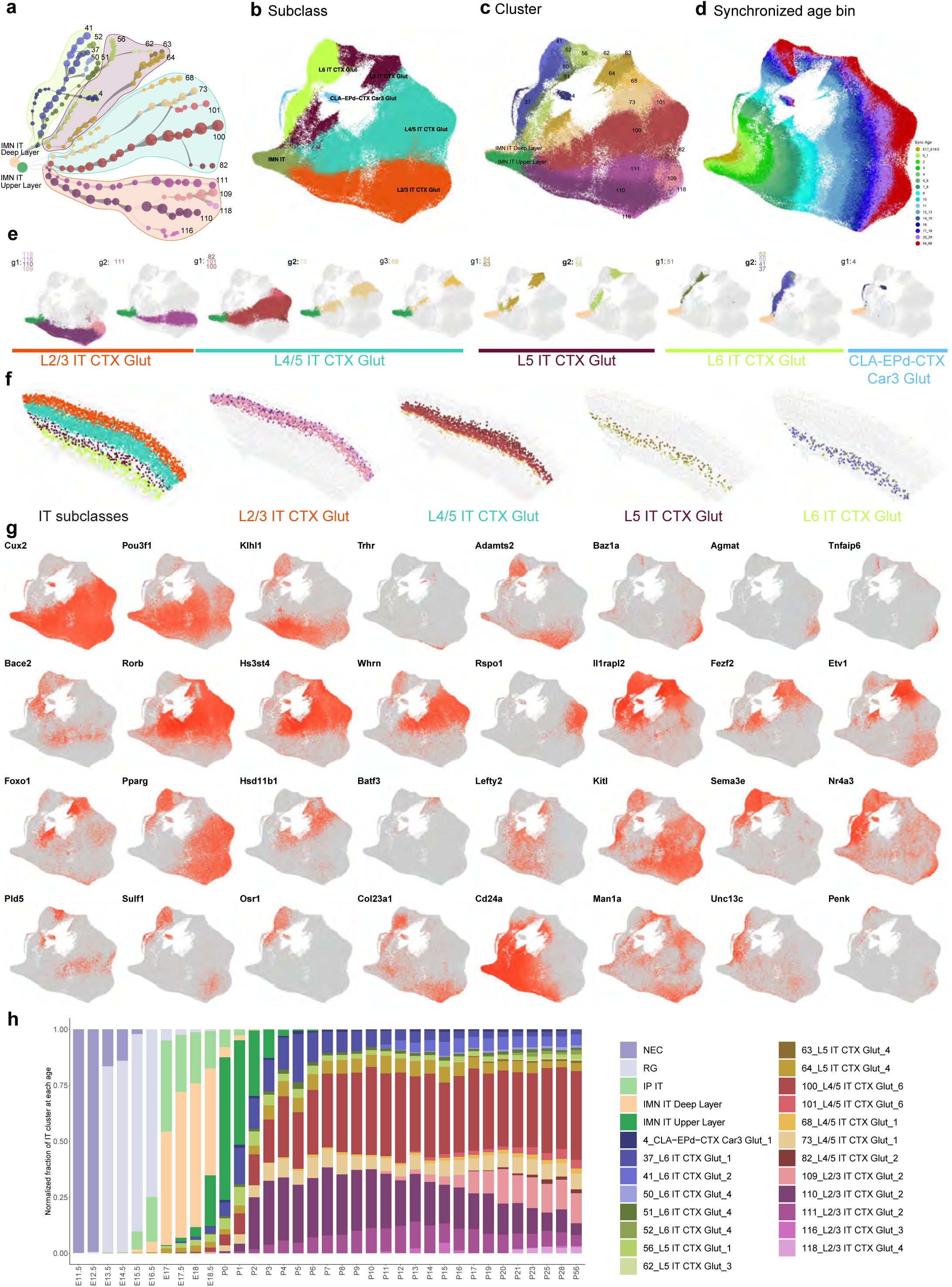
Developmental trajectories of visual cortex IT Glut cell types. **(a)** Transcriptomic trajectory tree for IT clusters starting from the common IMN IT antecedents. Nodes are clusters subdivided by synchronized age bins, and edges represent antecedent-descendent relationship between adjacent nodes, with thinner end at the antecedent node, and thicker end at the descendent node. Nodes are grouped by subclass, and adult clusters are labeled. **(b-d)** UMAP for nonIT cells colored by subclass (b), cluster (c) and synchronized age bin (d). **(e)** Clusters are grouped together based on similar trajectories. Within each cluster group, all cells along their trajectories, including all antecedent nodes, are shown and are colored by cluster membership. **(f)** Spatial distribution of IT subclasses and clusters within each subclass at adult stage, based on the ABC-WMB Atlas^15^. **(g)** Marker genes illustrating cell type diversification along trajectories. **(h)** Cluster composition of all IT cells at each age.

**Extended Data Figure 7.**
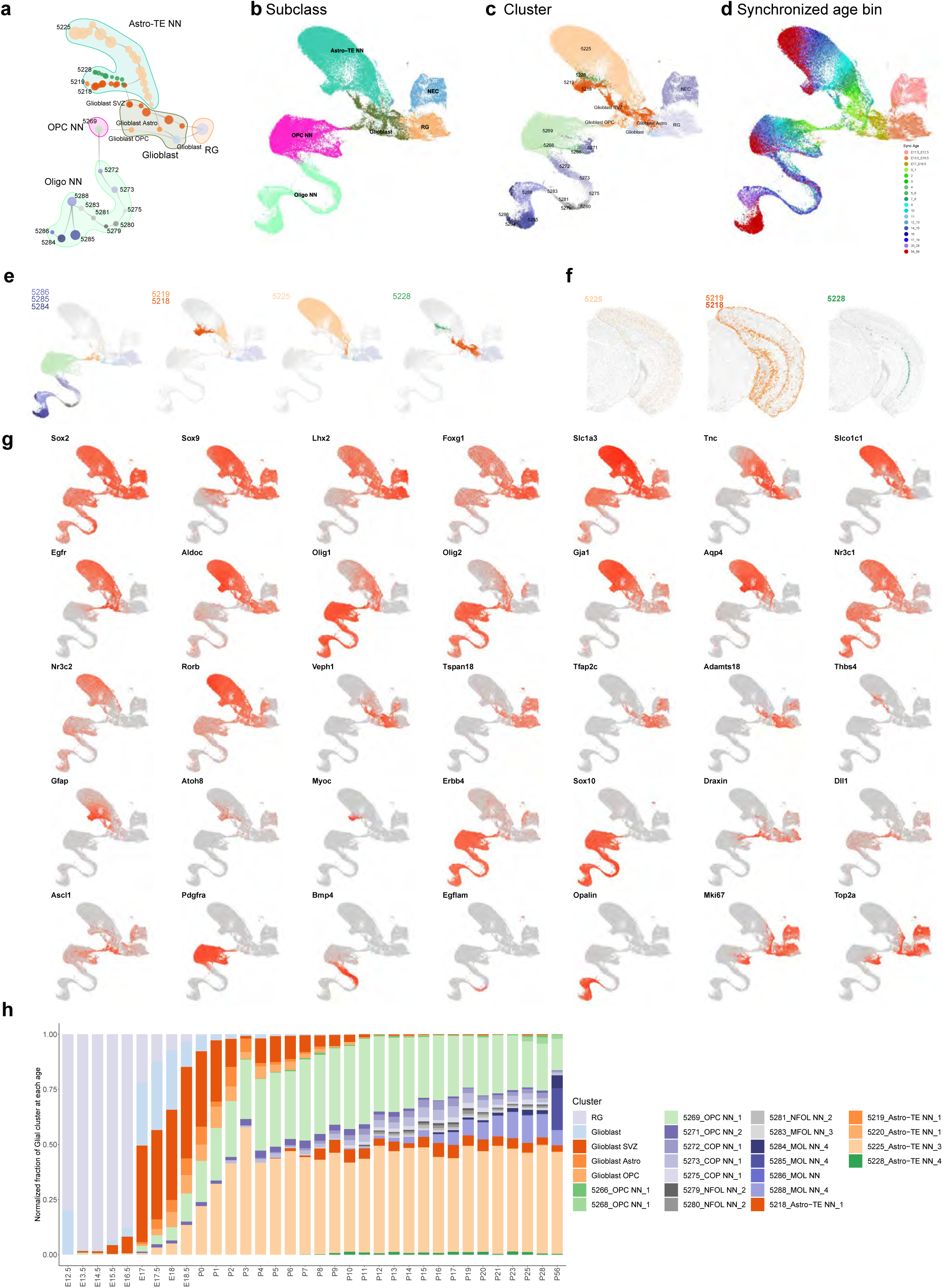
Developmental trajectories of visual cortex Glia cell types. **(a)** Transcriptomic trajectory tree for glia clusters starting from the common RG antecedent. Nodes are clusters subdivided by synchronized age bins, and edges represent antecedent-descendent relationship between adjacent nodes, with thinner end at the antecedent node, and thicker end at the descendent node. Nodes are grouped by subclass, and adult clusters are labeled. **(b-d)** UMAP for glial cells colored by subclass (b), cluster (c) and synchronized age bin (d). **(e)** Clusters are grouped together based on similar trajectories. Within each cluster group, all cells along their trajectories, including all antecedent nodes, are shown and are colored by cluster membership. **(f)** Spatial distribution of astrocyte clusters at adult stage, based on the ABC-WMB Atlas^15^. **(g)** Marker genes illustrating cell type diversification along trajectories. **(h)** Cluster composition of all glial cells at each age.

**Extended Data Figure 8.**
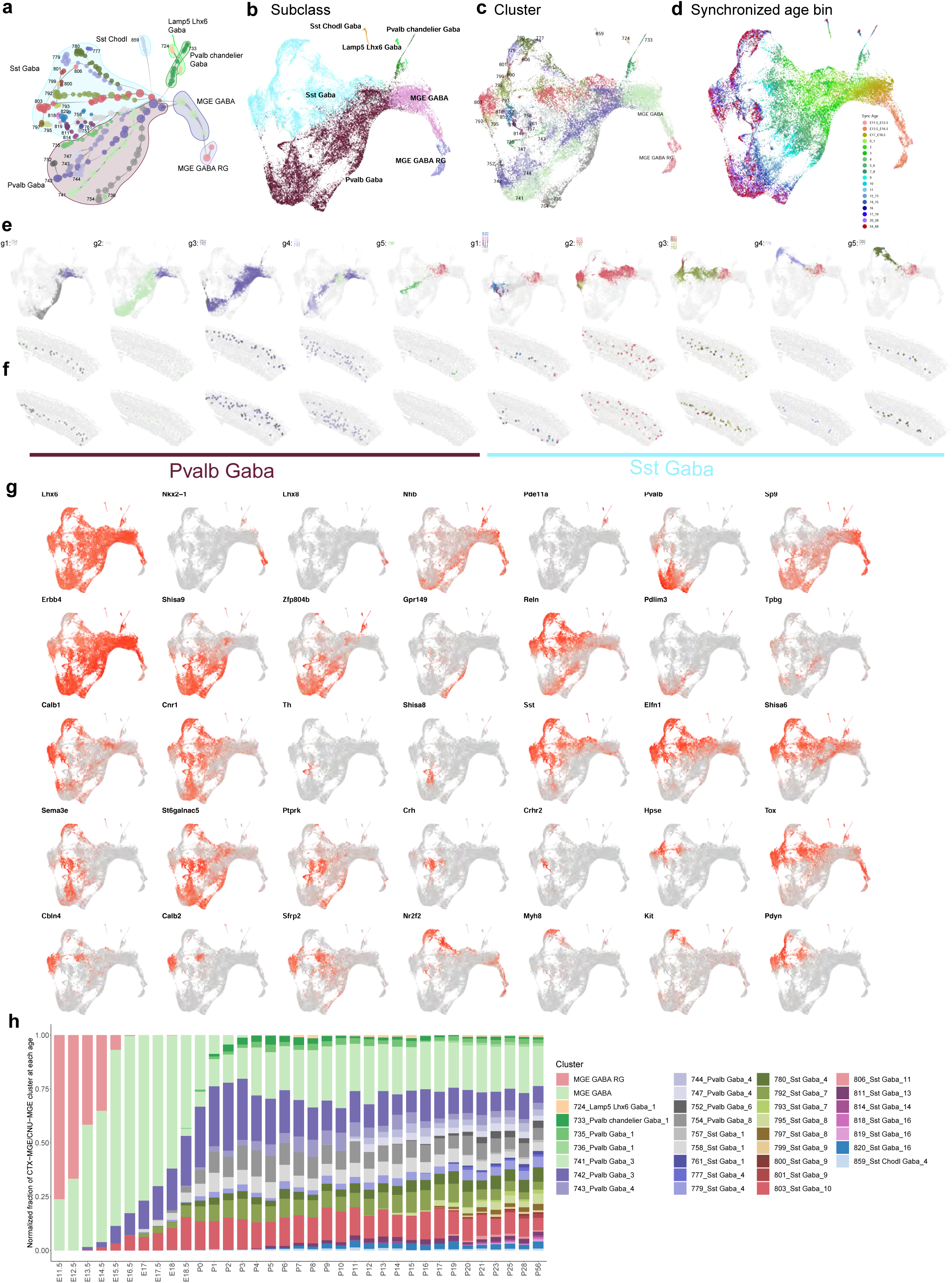
Developmental trajectories of visual cortex MGE GABA cell types. **(a)** Transcriptomic trajectory tree for MGE clusters starting from the common MGE GABA RG antecedent. Nodes are clusters subdivided by synchronized age bins, and edges represent antecedent-descendent relationship between adjacent nodes, with thinner end at the antecedent node, and thicker end at the descendent node. Nodes are grouped by subclass, and adult clusters are labeled. **(b-d)** UMAP for MGE cells colored by subclass (b), cluster (c) and synchronized age bin (d). **(e)** Clusters are grouped together based on similar trajectories. Within each cluster group, all cells along their trajectories, including all antecedent nodes, are shown and are colored by cluster membership. **(f)** Spatial distribution of MGE subclasses and clusters within each subclass at adult stage, based on the ABC-WMB Atlas^15^. **(g)** Marker genes illustrating cell type diversification along trajectories. **(h)** Cluster composition of all MGE cells at each age

**Extended Data Figure 9.**
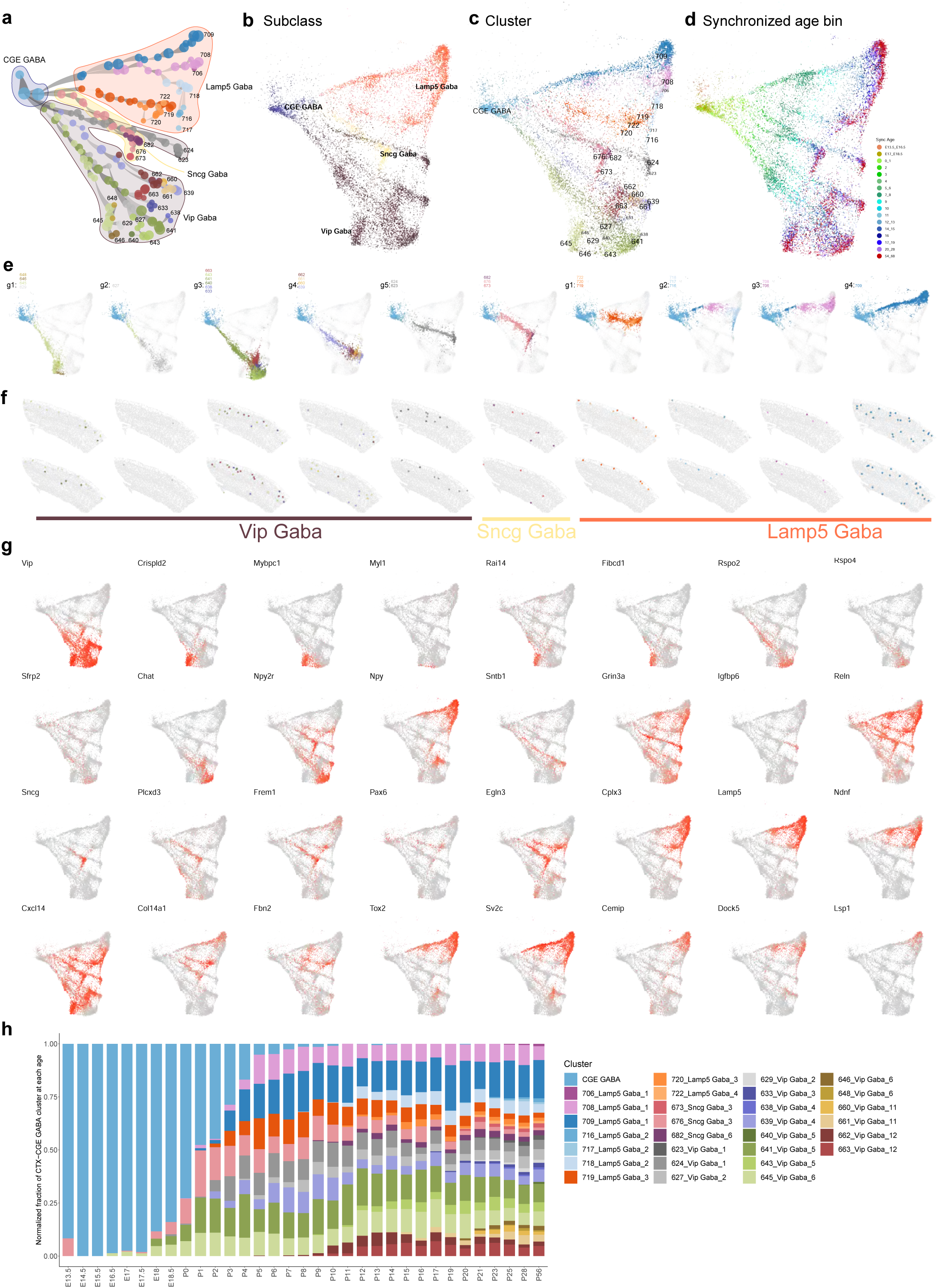
Developmental trajectories of visual cortex CGE GABA cell types. **(a)** Transcriptomic trajectory tree for MGE clusters starting from the common CGE GABA antecedent. Nodes are clusters subdivided by synchronized age bins, and edges represent antecedent-descendent relationship between adjacent nodes, with thinner end at the antecedent node, and thicker end at the descendent node. Nodes are grouped by subclass, and adult clusters are labeled. **(b-d)** UMAP for CGE cells colored by subclass (b), cluster (c) and synchronized age bin (d). **(e)** Clusters are grouped together based on similar trajectories. Within each cluster group, all cells along their trajectories, including all antecedent nodes, are shown and are colored by cluster membership. **(f)** Spatial distribution of CGE subclasses and clusters within each subclass at adult stage, based on the ABC-WMB Atlas^15^. **(g)** Marker genes illustrating cell type diversification along trajectories. **(h)** Cluster composition of all CGE cells at each age.

**Extended Data Figure 10.**
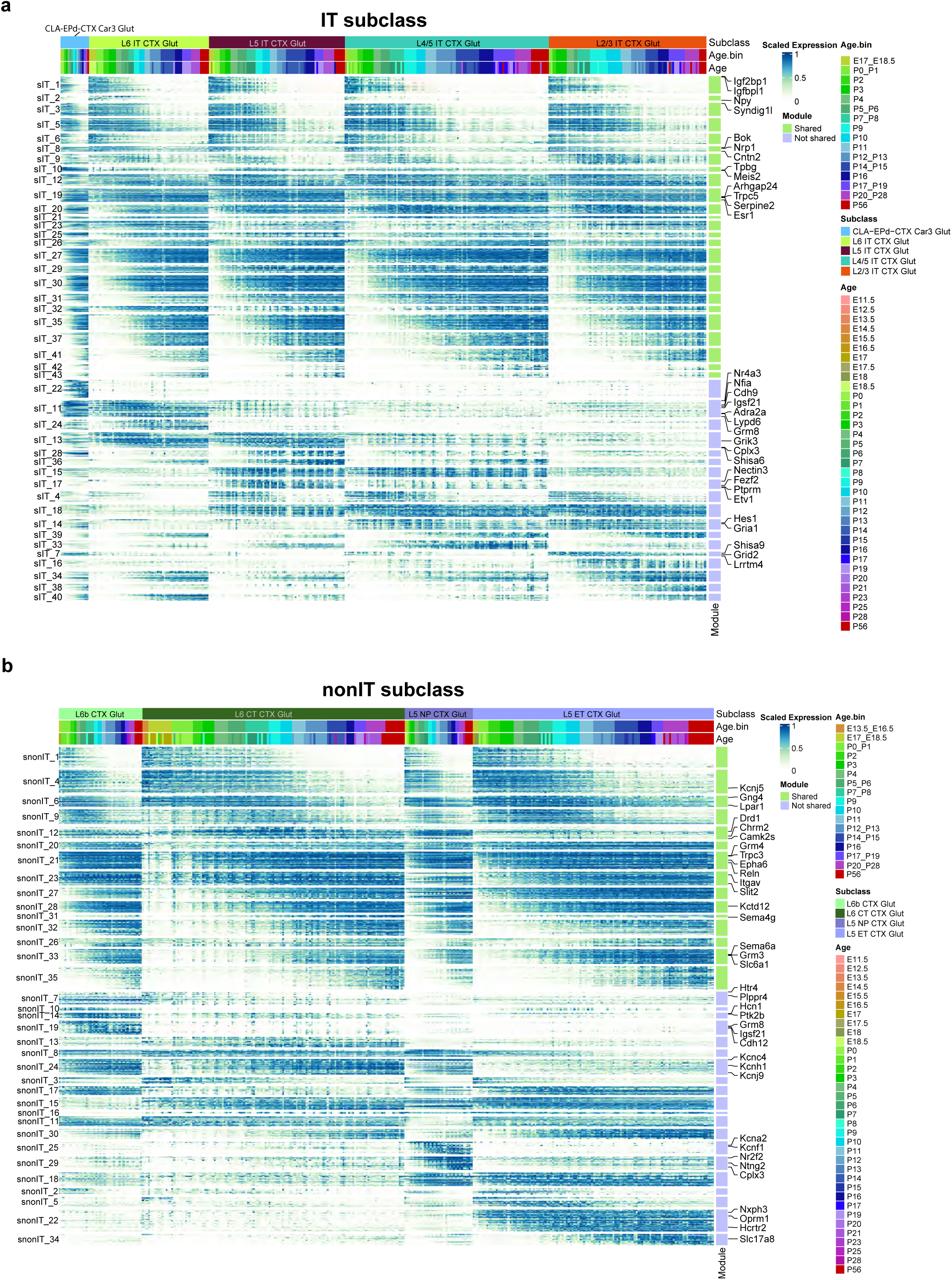
Gene modules across ages of glutamatergic subclasses. (a-b) Expression of DE genes for each subclass of IT (a) and nonIT (b) neurons, organized in gene co-expression modules shown as colored bars on the right of the heat map. Green and blue bars denote shared and subclass-specific modules, respectively. Module IDs are shown on the left, exemplary DE genes are shown on the right.

**Extended Data Figure 11.**
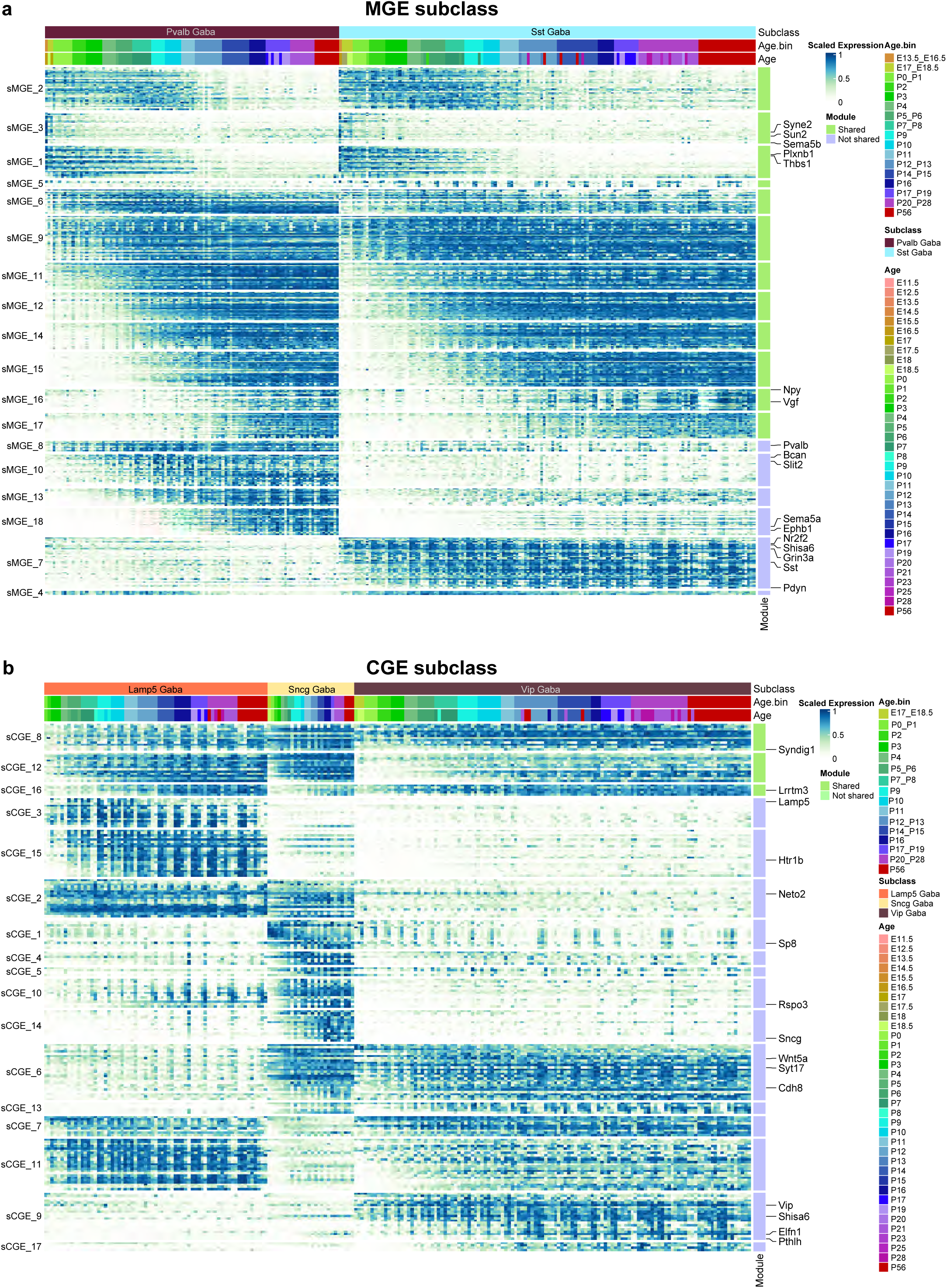
Gene modules across ages of GABAergic subclasses. (a-b) Expression of DE genes for each subclass of CTX-MGE (a) and CTX-CGE (b) neurons, organized in gene co-expression modules shown as colored bars on the right of the heat map. Green and blue bars denote shared and subclass-specific modules, respectively. Module IDs are shown on the left, exemplary DE genes are shown on the right.

**Extended Data Figure 12.**
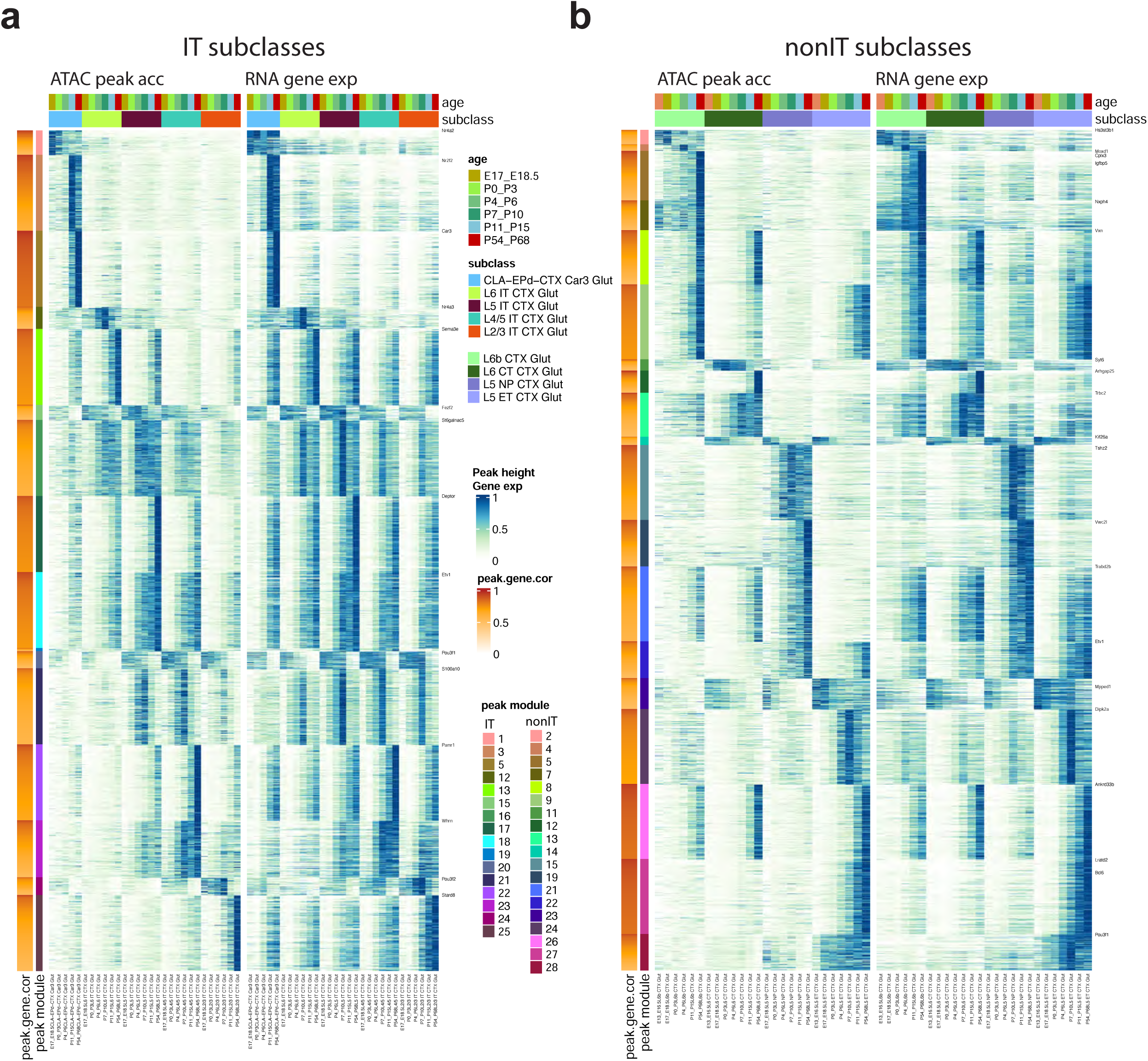
Correspondence of chromatin accessibility and gene expression across glutamatergic neuron types and ages during development. (a-b) Heatmap representation of corresponding peak accessibility and gene expression in IT subclasses (a) and nonIT subclasses (b). In each panel, each row corresponds to a peak/gene pair, ordered by peak module and peak/gene correlation, and each column corresponds to a cell category defined by subclass and age group. The left heatmap shows the average peak accessibility in each subclass-by-age-group category. Accessibility values are normalized, with maximum value of 1 per peak and 0 indicating no accessibility. The right heatmap shows the average gene expression in each subclass-by-age-group category. Expression values are normalized, with maximum value of 1 per gene and 0 indicating no expression.

**Extended Data Figure 13.**
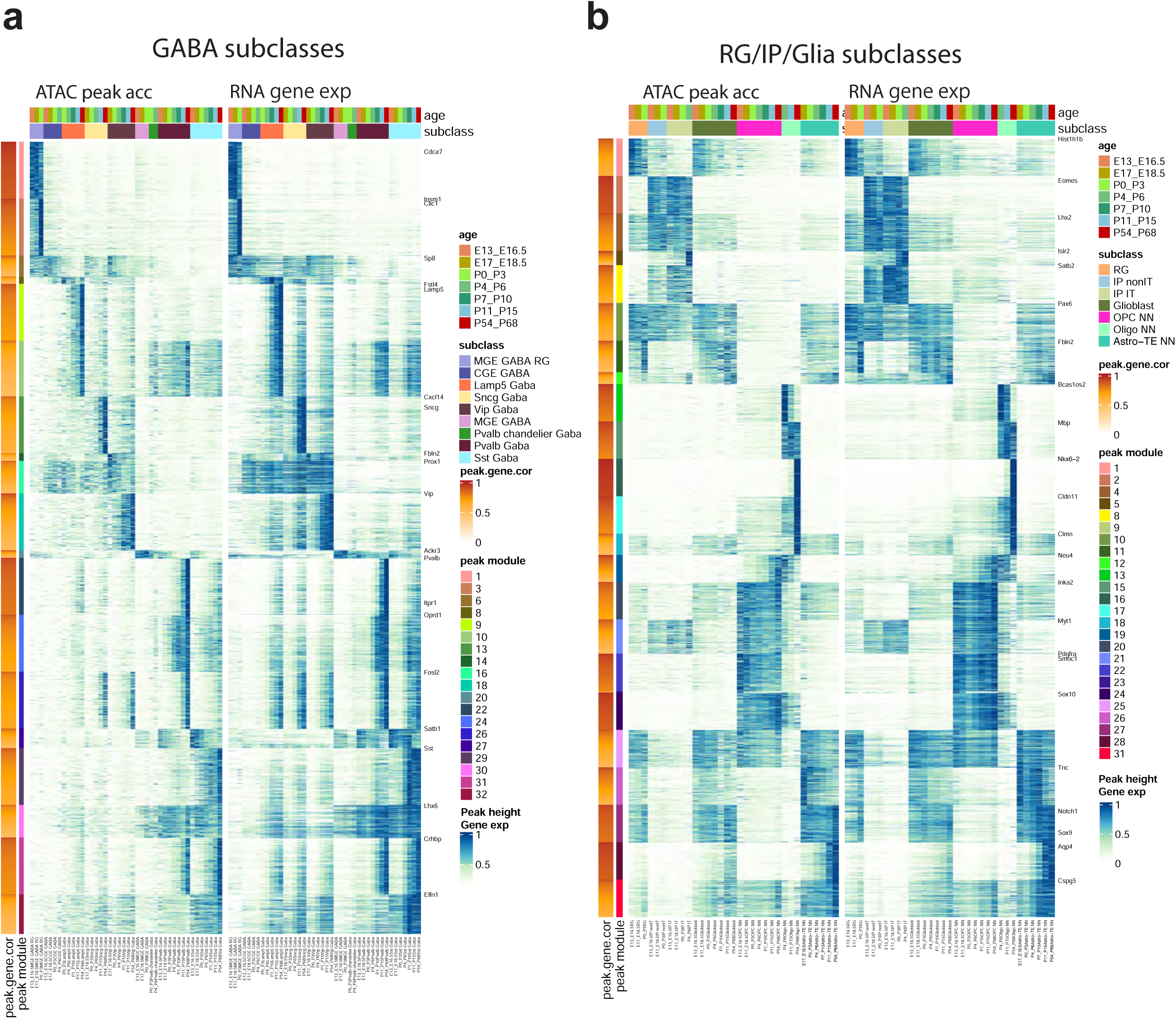
Correspondence of chromatin accessibility and gene expression across GABAergic and glial cell types and ages during development. (a-b) Heatmap representation of corresponding peak accessibility and gene expression in GABA subclasses (a) and glia subclasses (b). In each panel, each row corresponds to a peak/gene pair, ordered by peak module and peak/gene correlation, and each column corresponds to a cell category defined by subclass and age group. The left heatmap shows the average peak accessibility in each subclass-by-age-group category. Accessibility values are normalized, with maximum value of 1 per peak and 0 indicating no accessibility. The right heatmap shows the average gene expression in each subclass-by-age-group category. Expression values are normalized, with maximum value of 1 per gene and 0 indicating no expression.

**Extended Data Figure 14.**
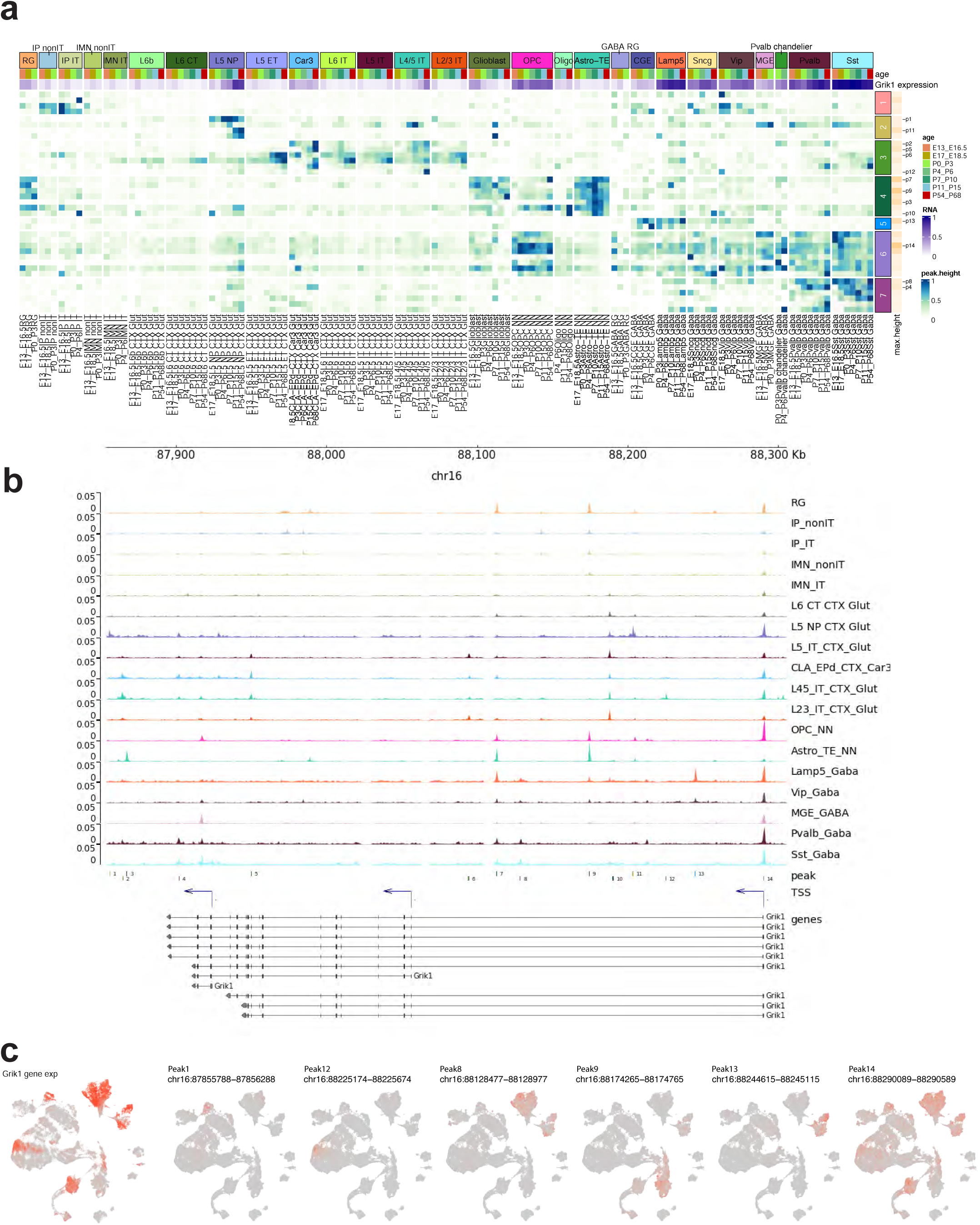
Differential accessibility peaks associated with the *Grik1* gene in different cell types or different developmental ages. **(a)** Heatmap representation of accessibility of differentially accessible peaks located in *Grik1* gene body and 50 Kb upstream. Each row corresponds to a peak, ordered by peak module, and each column corresponds to a cell category defined by subclass and age group. The *Grik1* gene expression level is shown in purple at the top. The heatmap color represents the average peak accessibility (height) in each subclass-by-age-group category, normalized with 1 indicating the maximum value for each peak and 0 indicating no accessibility. The peak module and maximum peak height are shown for each peak to the right. Specific peaks are numbered and labeled. **(b)** The accessibility tracks per subclass surrounding the *Grik1* gene, along with the genomic locations of labeled peaks in (a). TSS, transcription start site. **(c)** UMAP representation of Multiome cells, colored by *Grik1* expression and accessibility of a subset of peaks labeled in (a).

**Extended Data Figure 15.**
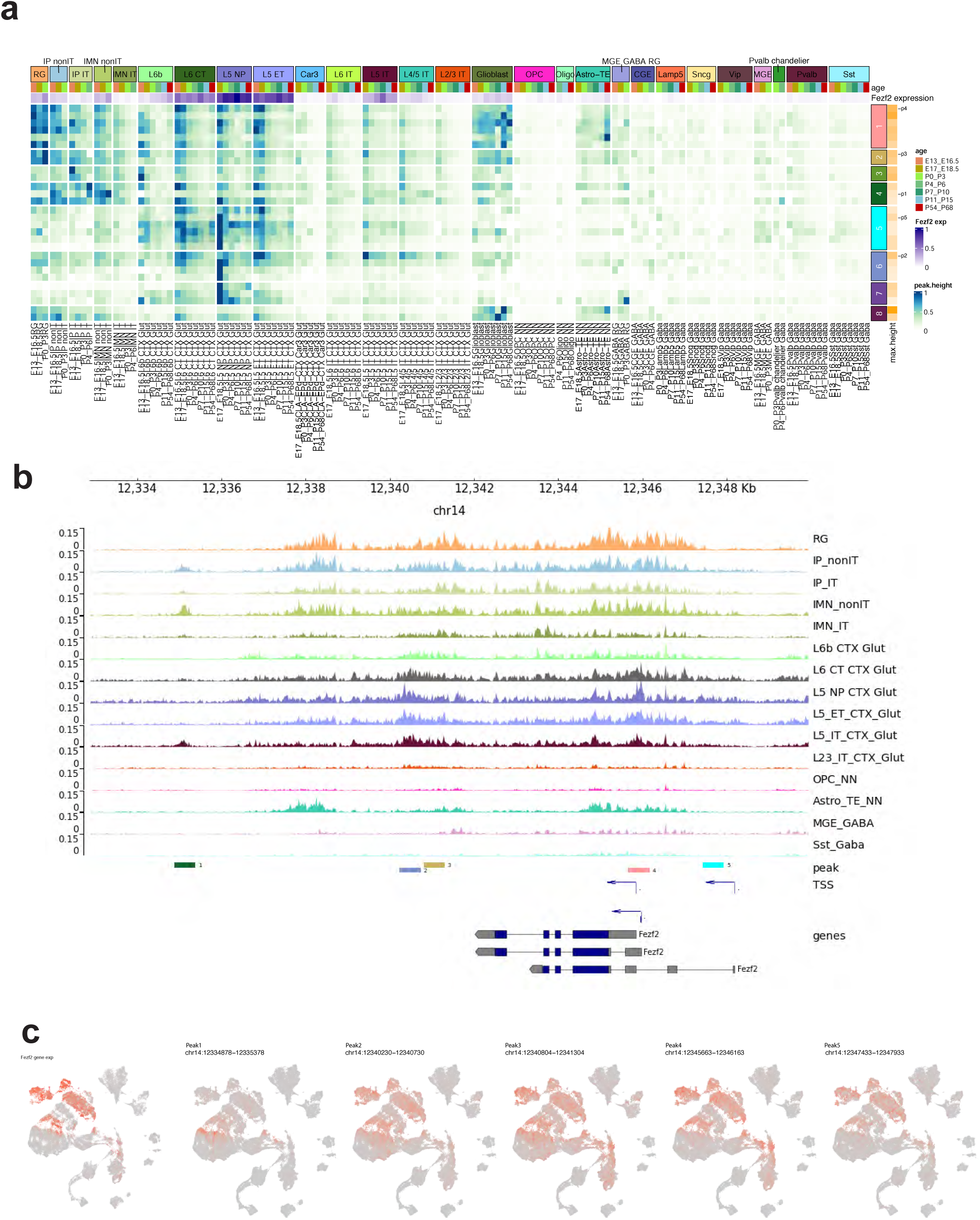
Differential accessibility peaks associated with the *Fezf2* gene in different cell types or different developmental ages. **(a)** Heatmap representation of accessibility of differentially accessible peaks located in *Fezf2* gene body and 50 Kb upstream. Each row corresponds to a peak, ordered by peak module, and each column corresponds to a cell category defined by subclass and age group. The *Fezf2* gene expression level is shown in purple at the top. The heatmap color represents the average peak accessibility (height) in each subclass-by-age-group category, normalized with 1 indicating the maximum value for each peak and 0 indicating no accessibility. The peak module and maximum peak height are shown for each peak to the right. Specific peaks are numbered and labeled. **(b)** The accessibility tracks per subclass surrounding the *Fezf2* gene, along with the genomic locations of labeled peaks in (a). TSS, transcription start site. **(c)** UMAP representation of Multiome cells, colored by *Fezf2* expression and accessibility of a subset of peaks labeled in (a).

